# High-resolution HIV-1 m^6^A epitranscriptome reveals isoform-dependent methylation clusters and unique 2-LTR transcript modifications

**DOI:** 10.1101/2025.03.06.641803

**Authors:** Delphine Naquin, Sandra Blanchet, Erwin van Dijk, Lisa Bertrand, Sylvie Grégoire, Cecilia Bertha Ramirez, Rania Ouazahrou, Yan Jaszczyszyn, Arnaud Moris, Claude Thermes, Céline Hernandez, Olivier Namy

## Abstract

The *N*^6^-methyladenosine (m^6^A) modification of HIV-1 has been widely studied but the number and precise positions of the m^6^A sites remain unclear due to the lack of precision of detection methods. Using the latest Nanopore chemistry and direct m^6^A base-calling, we identified 18 m^6^A: 14 at the 3’ end and 4 in central regions of the genome. Our data reveal differential methylation of these positions between splicing isoforms. Eleven of these sites are clustered in two short segments with peak-shaped methylation profiles. Single-molecule analysis revealed that a very small number of transcripts were unmethylated in both clusters. We also identified a ∼732 nt RNA species resulting from the transcription of non-integrated viral DNA circles closed by two long terminal repeats. These transcripts started in the first LTR, terminated at the polyA site of the second LTR and harbored six m^6^A sites. Five of these sites were present in other transcripts and, remarkably, had the highest methylation rates. The sixth site was methylated only in this transcript, suggesting a role for this RNA in HIV-1 infection. These findings reveal a new landscape of HIV m^6^A transcriptome modifications and pave the way for studies deciphering their role in the viral life cycle.

## INTRODUCTION

Epitranscriptomic RNA modifications play crucial roles in diverse cellular and viral processes and diseases ^1^. The N6-methyladenosine (m^6^A) modification occurs frequently, in all types of cellular RNA, and consists of the covalent addition of a methyl group to the N6 position of adenosine. This methyl group addition is catalyzed by a multi-subunit methyltransferase complex with a catalytic core composed of methyltransferase-like 3 (METTL3) and methyltransferase-like 14 (METTL14) ^2^. This writer complex preferentially targets adenosines in the consensus sequence 5′-DRACH-3′ (D = A/G/U, R = A/G, H = U/A/C) in mammalian mRNAs ^3^. DRACH motifs are frequent in mRNA but are not always targeted, as only about 10% of these motifs were found to be m^6^A-methylated in cells. This suggests that there are probably other context-dependent signals for the methylation of specific DRACH motifs. The m^6^A modification is reversible and can be removed by erasers, such as the fat mas and obesity-associated protein (FTO) ^4^ or alkB homolog 5 (ALKBH5) ^5^. Reader proteins recognize m^6^A and determine the functional consequences of the modification for the fate of the RNA. The readers capable of binding m^6^A include the YTH domain-containing family of proteins, the most frequently studied members of which are YTHDC-1, YTHDF1,2,3 ^6,7^. However, discrepancies remain concerning the function of these proteins ^8^. In mammals, m^6^A is the most prevalent modification involved in gene expression regulation and RNA metabolism, including mRNA transport, localization, stability, translation efficiency and processing ^9–13^, but this modification is also present in viral RNA ^14^. Several studies have detected methylated regions (m^6^A) in the HIV-1 genome with techniques based on methylated RNA immunoprecipitation ^15–20^. However, it was not possible to localize the methylated positions with single-nucleotide precision because the resolution of the techniques used ranged from 20 nt to more than 200 nt. There were also differences between studies, with some reporting peaks distributed along most of the length of the genome ^20^, whereas others detected peaks only at the 5’ and 3’ ends ^15,19^ or only at the 3’ end ^18^. These discrepancies may reflect differences between the cell types or infection protocols (cell transfection with proviral plasmids or infection with viruses) used or a lack of reproducibility of antibody-based methods ^21^. In principle, Oxford Nanopore Technologies (ONT) direct RNA sequencing ^22^ would allow the detection of modified RNA nucleotides at single-molecule level. Several studies have provided diverse bioinformatics tools for the detection of RNA modifications based on various strategies (base-call errors induced by modifications, altered current intensity, modification-aware base-calling models, reviewed in ^23^). However, improvements in data quality and recovery rates for full-length transcripts are required to enable direct base-calling for the modified nucleotides ^24^. Here, we considered the full-length genome, including the 5’LTR and 3’LTR sequences. A recent study using the RNA002 kit resulted in the development of a modified library protocol and major improvements in the recovery of full-length RNA molecules ^25^. Combining several bioinformatics tools based on different properties of the ionic current, the authors identified three m^6^A sites at the 3’ end of the viral genome. We made use of an advance developed by ONT — RNA004 chemistry (new nanopore and new motor protein) — that greatly improves overall yield and data quality ^26^, and new base-calling models making it possible to call m^6^As directly with the four standard bases. Using these developments, we characterized the HIV-1 m^6^A methylation pattern of HIV-1 mRNAs in infected CD4^+^ T cells, including a CD4^+^ T cell line (SupT1) and, importantly, primary CD4^+^ T cells. We extended the detection to 18 m^6^A sites at single-nucleotide resolution, single-molecule level and in different splicing isoforms, providing the most complete and precise description of the landscape of HIV-1 m^6^A methylation to date.

## MATERIALS AND METHODS

### Cell infection

VSV-G pseudotyped HIV_NL4-3 XCS_ was produced by transfection of 293T cells using CAPHOS kit (Sigma-Aldrich) as in^54^. Viruses were pseudotyped to favor viral entry and to achieve a high infection rate within a short time. SupT1 CD4^+^ T cells were cultured with RPMI GlutaMax 1640 (Gibco) complemented with 10% FBS (Dutscher) and 1% penicillin/streptomycin.

2 x 10^8^ SupT1 CD4^+^ T cells were infected with 800 ng of VSV-G-HIV_NL4-3 XCS_ HIV-Gagp24 for 3h in IMDM plus 10mM HEPES (Gibco/ThermoFisher Scientific) supplemented with 2 µg/ml of DEAE-dextran (Sigma). Two biological replicates were performed corresponding to 100 million cells infected at time 0. Eighteen hours post-infection a fraction of the cells (2 x 10^5^) was harvested, stained with CD4-Vio770 antibody (Miltenyi) and a Live/Dead (LVD) fixable violet dye (Invitrogen), fixed with 4% paraformaldehyde (ChemCruz) and permeabilized (PBS1X, BSA 0,5%, saponin 0,05%). Cells were then stained with an HIV-1 Gag-p24 specific antibody (KC57-PE, Beckman Coulter). Sample acquisition and analysis were performed using a BD LSRFortessa and FlowJo v10.8 Software, respectively (both from BD Life Sciences). Mock-infected cells were used as negative control. Twenty and thirty hours post-infection, cells were harvested, washed in cold PBS and lysed at 4°C in RLT (from RNeasy Mini Kit (Qiagen) buffer containing ß-mercaptoethanol and 1% NP40.

PBMCs from anonymous blood donors (n=2, donor 864 and 880) were purchased from the blood bank at EFS (Établissement Français du sang) under the agreement number 15/EFS/022. PBMCs were isolated using a Ficoll gradient and monocytes depleted by overnight incubation in lying cell culture flasks containing RPMI+10%FBS supplemented with 10 IU/ml of recombinant human IL-2 (rhIL-2, Miltenyi Biotec). CD4^+^ T cells were then isolated using CD4^+^ magnetic beads (Miltenyi Biotec) and activated with Phytohemagglutinin (PHA, 1 µg/ml, PAA Laboratories) in RPMI GlutaMax 1640 complemented with 10% FBS, 1% penicillin/streptomycin, nonessential amino acids and sodium pyruvate (all from Gibco/ThermoFisher Scientific), so called T-cell medium. 48 h post-activation, CD4^+^ T cells were harvested and resuspended in T-cell medium containing 100 IU/ml of recombinant human IL-2 (rhIL-2, Miltenyi Biotec). 7-days post activation, 300 x 10^6^ and 130 x 10^6^ CD4^+^ T cells, were infected (for donor 864 and donor 880, respectively) at a multiplicity of infection (moi) 0,01 with VSV-G-HIV_NL4-3 XCS_ (representing around 1000 ng of HIVGagp24 / 10^7^ cells) for 3h in IMDM plus 10mM HEPES supplemented with 2 µg/ml of DEAE-dextran. CD4^+^ T cells were then maintained in T-cell medium containing 100 IU/ml of rhIL-2 and HIV infection was monitored using flow cytometry, as for SupT1 CD4^+^ T cells. For donor 864 and donor 880, cells were harvested and lysed 48 h and 72 h post-infection, respectively.

### In vitro transcription

HIV-1 genomic RNA was transcribed in two parts, from the beginning of 5’LTR to the end of Pol (5124 bases) and from the end of Pol to the end of 3’LTR (4829 bases), with 225 overlapping bases. Each part of the genome was PCR amplified from pNL4.3XCS^55^ and cloned in a pUC19 vector containing T7 promoter, using NEBuilder cloning kit (New England Biolabs). The DNA fragment corresponding to the 2LTR (synthesized by IDT) was cloned into the same plasmid. Owing to synthesis constraints, the 3′ R repeat at the distal end of the 2LTR element was omitted, yielding a truncated 2LTR sequence of 638 nt. The plasmids were then linearized with SmaI restriction enzyme during 3 hours at 30°C, to perform run-off transcription. In vitro transcription was performed using RiboMAX T7 Large Scale RNA Production System (Promega) with incubation 4 hours at 37°C, followed by DNAseI treatment during 15 min and RNA column clean-up (ZymoResearch).

### RNA purification

Total RNA was extracted from the cell pellets using RNeasy Mini Kit (Qiagen), according to the manufacturer’s protocol. Briefly, cells were lysed in RLT buffer containing b-mercaptoethanol and homogenized using QIAshredder homogenizer. The lysate was then purified on RNeasy spin columns and RNA was eluted in RNase-free water. PolyA RNA were then purified from the total RNA using two rounds on Oligo(dT)_25_

Dynabeads (Thermo Fisher Scientific). In vitro transcripts were polyadenylated using E. coli PolyA polymerase kit from NEB and purified on Oligo(dT)_25_ Dynabeads.

### Library preparation and sequencing

The polyA RNA was used for ONT direct RNA sequencing using the RNA004 kit. In the case of demethylase treatment (see below), RNAs were first treated and carefully purified before library preparation. The reverse transcription (RT) step was modified as described by ^25^; a pool of 111 oligonucleotides complementary to the HIV-1 genome was added during library preparation in an estimated 30-fold molar excess to the viral RNA, which resulted in a much more efficient reverse transcription process.

Sequencing was processed on P2solo through the GridION (MinKNOW version: 24.02.10 for SupT1_20h_pi rep1, 24.02.16 for SupT1_30h_pi rep1 and IVT, 24.06.15 for SupT1_20h_pi+demeth rep1 and rep2, SupT1_20h_pi rep2, SupT1_30h_pi rep2, CD4_48h_pi and CD4_72h_pi, IVT). After sequencing, base-calling has been done with dorado-0.7.2 and rna004_130bps_sup@v5.0.0 model with –modified-bases m^6^A parameter (for all samples).

### Identification of HIV-1 splicing isoforms

Pass reads were mapped with minimap2 (v2.17-r941) (with -x splice and –secondary=no options) on the PNL4-3XCS HIV-1 genome. BamToBed tools with -splice option have been used to select full-length transcripts using a homemade script python. With this script, each read is analyzed and its splice junctions must match (with a tolerance of one base) the splice junctions of a specific isoform. In each full-length read we identified the donor (D) and acceptor (A) sites that match the already known HIV-1 splice sites (eight donor and eleven acceptor sites, Nguyen Quang et al., Emery et al., Ocwieja et al., O’Reilly et al., Widera et al.) (Table S2A). Some rare isoforms were detected with only few reads and only those detected in at least five of the six samples were considered. Finally, 51 isoforms were identified by their succession of splice junctions and were named as in (Nguyen Quang et al. 2020) (Table S2B). Between 4.4% (CD4_48h_pi) and 8.0% (SupT1_20h_pi rep1) of full-length reads did not fully match these sites and were not considered.

### Detection of m^6^A sites, computation of methylation rates

Nucleotide position of m^6^A’s was obtained thanks to the specific base-calling process that can detect m^6^A’s (version 5.1.0). For each sample, the methylation rate is computed by modkit pileup with –filter-threshold A:0.8, --mod-thresholds a:0.9 and –edge-filter 20. The nucleotides that are called as m^6^A are counted at each position in a set of reads to determine the corresponding modification rate. The methylation rates computed at each position of the transcripts in a sample were compared to those computed at the corresponding positions of the IVT sample using <u>DMR model and scoring details - Modkit</u> that computes a final modification rate and a corresponding p-value at each position. The positions presenting p-values p ≤ 10^-5^ and a number of methylated reads ≥ 30 were retained as putative m^6^A sites. For the clustering analyses in Figure 3 and Figure S4, only the isoforms identified by more than 100 full-length reads were retained.

To detect m^6^A between modified and unmodified oligonucleotides, the sequence was different in order to distinguish between them (by a tag of 5 nucleotides (Materials and Methods)). The DMR model could therefore not be used, and the methylation rate was calculated by subtracting the modified from the unmodified values.

### Single molecule analysis of methylated m^6^A sites

Modkit extract enables access to read-level base modification calls. Parameters –mapped-only and –include-bed were used (bed file containing eight positions from cluster1 and three positions from cluster2). The output tsv files were parsed with the R language. During this analysis, the threshold of probability of modification to call 6mA was 0.9 (as with modkit pileup).

### Synthetic oligonucleotides

We used two synthetic RNA oligonucleotides identical to HIV-1 regions with methylated As and the same two oligonucleotides differing only at extremities, without methylated As.

Oli1-mod harbors m^6^A at positions 8532 et 8563 + 5nt tag at 3’ end (**TATGC)**: CCGCTTGAGAG(m^6^A)CTTACTCTTGATTGTAACGAGGATTGTGGA(m^6^A)CTTCTGGGAC**TATGC** (58 nt-long).

Oli1-nmod harbors non-modified A’s and 5nt tag at 3’ end (**GTACG**): CCGCTTGAGAGACTTACTCTTGATTGTAACGAGGATTGTGGAACTTCTGGGAC**GTACG** (58 nt-long).

Oli2-mod harbors m^6^A at positions 9400, 9428 et 9442: CTTCAAGA(m^6^A)CTGCTGACATCGAGCTTGCTACAAGGG(m^6^A)CTTTCCGCTGGGG(m^6^A)CTTTCCAGG (60 nt-long).

Oli2-nmod harbors non-modified A’s and 6nt tag at 3’ end (**TCATAG**): CTTCAAGAACTGCTGACATCGAGCTTGCTACAAGGGACTTTCCGCTGGGGACTTTCCAGG**TCATAG** (66 nt-long).

Oli3-mod harbors m^6^A at positions 8436 and 8442: AAGGAATAGAAGAAGAAGGTGGAGAGAGAG(m^6^A)CAGAG(m^6^A)CAGATCC (44nt-long)

Oli3-nmod harbors non-modified A’s and 5nt tag at 3’ end (**TATGC**): AAGGAATAGAAGAAGAAGGTGGAGAGAGAGACAGAGACAGATCC**TATGC** (49 nt-long)

Oli4-mod harbors m^6^A at position 9163: ACACACAAGGCTACTTCCCTGATTGGCAGA(m^6^A)CTACACA (38 nt-long)

Oli4-nmod harbors non-modified A’s and 5nt tag at 3’ end (**GTAGC**): ACACACAAGGCTACTTCCCTGATTGGCAGAACTACACA**GTAGC** (43 nt-long)

### Oligonucleotides demethylation and sequencing

As recommended by the manufacturer, less than 3 µM of m^6^A was incubated with 1 µM ALKBH5 at room temperature. The composition of the reaction mixture was as follows: 3 µL enzyme (0.9 µg, 19.8 pmol), a mix of four oligos at 0.13 µM each (oli1-mod, oli1-nmod, oli2-mod and oli2-nmod). A total of (3 x 0.13) + (2 x 0.13) = 0.65 µM m^6^A was used in this step; the buffer contained 100 mM KCl, 2 mM MgCl2, 2 mM L-ascorbic acid, 300 µM alpha-ketoglutarate, 150 µM (NH4)2Fe((SO4)2 and 50 mM Hepes buffer (pH 6.5), in a total volume of 20 µL. The reactions were stopped after 1.5 h incubation by adding EDTA to a final concentration of 5 mM; the oligos were purified using SPRIselect beads following a modified protocol as follows. Water was added to a 25 µL total volume, followed by the addition of 45 µL SPRI beads and mixing. After 5 minutes of room temperature incubation, the beads were magnetized and the supernatant containing the small RNA was transferred to a new tube containing 45 µL SPRI beads and 135 µL isopropanol, and mixed. After another 5 minutes of room temperature incubation, the beads were magnetized and the supernatant was discarded. The beads were washed twice with 86% ethanol, allowed to dry for 5 minutes and resuspended in 10 µL water. The RNA oligos were then polyadenylated to make them suitable for direct RNA sequencing as follows. They were incubated with 5 U E. coli poly(A) polymerase (PAP) (New England Biolabs) in 20 µL reactions containing 1 mM ATP and 1x PAP buffer at 37 °C for 1.5 minutes. Then EDTA was added to 10 mM final concentrations and RNA was again purified using SPRI beads as described above. The purified and polyadenylated oligos were then used for ONT direct RNA004 library preparation and sequenced on two MinION flow cells (MinKNOW version: 24.06.14). Distinction between oligos has been made by mapping to the reference with these four oligos added by eight As in 3’.

As an alternative to ALKBH5, RNA was also treated with the FTO RNA demethylase (New England Biolabs) following the manufacturer’s recommendations. 100 ng RNA was incubated with 1 µL (1µg) FTO enzyme in a 25 µL reaction containing 75 µM Fe(II) and 1 x FTO buffer for 1 hour at 37 °C. After the incubation the same procedure was followed as done for ALKBH5 to stop de reactions, purifiy and polyadenylate the RNAs.

### Linearity of the m^6^A detection method

To test the linearity of the m^6^A detection method, different mixtures of the modified and non-modified RNA oligonucleotides (oli-mod+oli-nmod) were prepared and sequenced and the detected methylation rates were compared with the expected rates. For this, an equimolar pool of methylated oligos (oli1-mod + oli2-mod + oli3-mod + oli4-mod) and an equimolar pool of non-methylated oligos (oli1-nmod + oli2-nmod + oli3-nmod + oli4-nmod) were prepared; these pools were subsequently mixed in 3 different ratios to obtain the expected methylation levels, namely 25 %, 50 % and 75 %. These mixtures were then used for ONT direct RNA004 library preparation and sequenced on three MinION flow cells (MinKNOW version: 25.03.9). Distinction between oligos was performed as described in the previous section leading to distinct sets of modified and non-modified oligos that were used to compute the observed methylation rates corresponding to the expected values of 0% and 100%,

### Poly(A) RNA demethylation and sequencing

Biological polyA RNA was treated with demethylases ALKBH5 or FTO in the same way as described for the oligos with the following few modifications; 300 ng or 100 ng RNA was used per reaction for ALKBH5 or FTO respectively. RNA purification was done by adding 2x reaction volumes of SPRI beads followed by ethanol wash and elution in 10 µL water.

## RESULTS

### Nanopore sequencing of full-length HIV-1 transcripts and identification of mRNA isoforms

For the detection of HIV-1 m^6^A sites, cellular transcripts from SupT1 and primary CD4^+^ T cells infected with HIV-1 were extracted and sequenced by ONT direct RNA sequencing (DRS, Materials and Methods). It has been reported that it is very difficult to obtain reads corresponding to full-length HIV RNA molecules with the standard DRS library preparation protocol ^25^. In our previous DRS experiments with the standard protocol, we obtained less than 0.008% and 0.06% of total reads for our first two samples, corresponding to the ∼9.6 kb unspliced HIV-1 genome. In this study, we used a modified protocol described elsewhere ^25^ to increase recovery rates for full-length transcripts. We used two biological replicates of SupT1 cells (rep1 and rep2) harvested 20 h and 30 h post-infection and primary CD4^+^ T cells isolated from the blood of two different blood donors collected 48 h and 72 h post-infection (Figure S1). PolyA RNA was isolated from each of these six infected samples and sequenced to generate 7.3 to 28.9 million total reads (Table S1) and between 33,587 and 164,360 reads corresponding to full-length HIV transcripts (Table S1). Full-length reads were examined for the presence of splice junctions. We identified eight known donor sites (four major sites — D1, D2, D3, D4 — together with D1c, D2b, D4a and D4b) and eleven acceptor sites (four major sites — A1, A2, A5, A7 — together with A1b, A3, A4a, A4b, A4c, A5a and A5c) ^27–30^ (Table S2A). The different isoforms were identified by the succession of donor and acceptor sites (Materials and Methods); we identified 51 isoforms, as previously described ^27,31^ (Table S2B). These isoforms are typically classified into three groups on the basis of size: (i) unspliced ∼9 kb transcripts (US), (ii) partially spliced transcripts (∼4 kb, PS, lacking intron 1 and retaining intron 4) including Vif2, Vpr3,4, Env/Vpu and Tat5, and (iii) completely spliced transcripts (∼2 kb, CS, lacking introns 1 and 4) including Vif1, Vpr1,2, Tat1-4, Rev and Nef. The relative abundances of the various isoforms were similar in all samples (correlation R>0.86 in all comparisons, Table S3).

### Detection of m^6^A sites in all HIV-1 transcripts

The first step was the detection of m^6^A sites in all HIV-1 reads (Material and Methods). The positions of N6-methyladenosine nucleotides were determined directly at read level during the base-calling process, providing a first estimate of methylation rates at each position in a set of transcripts. These methylation rates were compared with those for the corresponding positions in the unmodified RNAs transcribed *in vitro* (Figure 1B). We performed this analysis for all HIV-1 transcripts from the six samples, resulting in a set of 14 putative m^6^A sites, all located in the 3’ region of the HIV-1 genome common to all transcripts (Tables 1, S4; Figure 1A). The numbering used here corresponds to that of the full-length genome, including the 5’LTR and 3’LTR sequences (transcription start site in position +454). The six samples all present a similar methylation landscape showing two clusters of sites. The first cluster consisted of eight sites (A8436 to A8632) with two highly methylated central sites (64-80%), and the second cluster consisted of three sites (A9400, A9428, A9442) with a highly methylated central site (79-84%). Three additional sites displaying low rates of methylation were detected between these two clusters. Remarkably, the segment containing the first cluster (8436 to 8632) was found to harbor five DRACH sequences ^3,32^, all of which were methylated. The other three sites of this segment correspond to 1-nt variants of the DRACH motif (found here at positions A8487, A8555, and A8572) interspersed with the canonical DRACH sites (Tables 1, S4; Figure 1A). It should be noted that m^6^As have also been detected in non-DRACH motifs, especially the variant GGAC<u>G</u> (found here at positions A8487 and A8572), which has been found to be highly methylated in human mRNAs ^33,34^.

**Figure 1.**
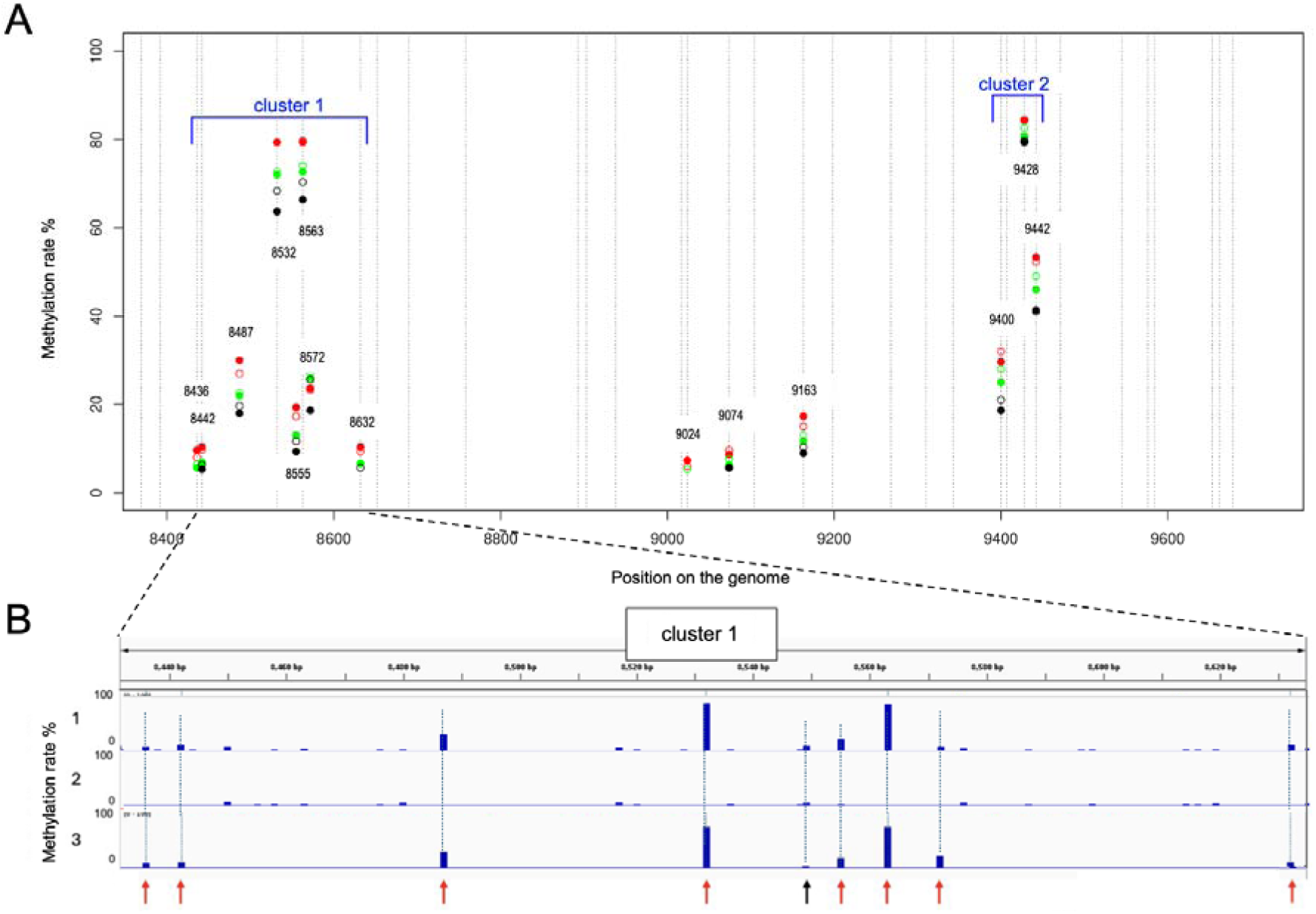
Detection of methylation signals at A nucleotides in all HIV-1 transcripts in the six samples. **(A)** Methylation profile of all HIV-1 transcripts in the 3’ region (8360-9700). Closed red circles, SupT1_20h_pi rep1; empty red circles, SupT1_30h_pi rep1; closed green circles, SupT1_20h_pi rep2; empty green circles, SupT1_30h_pi rep2; closed black circles, CD4_48h_pi; empty black circles, CD4_72h_pi. Vertical lines, positions of As in the 33 DRACH sequences of the region. **(B)** The IGV screenshot shows the methylation rate measured in the cluster 1 region of the HIV-1 genome including sites A8436 to A8632. Line 1, raw methylation rate measured directly during the m^6^A base-call of SupT1_20h_pi rep1. Line 2, raw methylation rate measured directly for the in vitro sample in the same region. Line 3, Final methylation rate computed as the difference (line 1) - (line 2); red arrows indicate the eight detected m^6^A sites; negative values are set to zero; sites with rates < 3% were not retained (black arrow).

### Efficiency of m^6^A detection in synthetic oligonucleotides

We assessed the efficiency of m^6^A site detection by subjecting two synthetic RNA oligonucleotides with sequences corresponding to HIV-1 segments but harbouring m^6^A at positions 8532 and 8563 (oli1-mod) and at positions 9400, 9428 and 9432 (oli2-mod) to the detection procedure. As controls, we used and analyzed two other synthetic HIV RNA oligonucleotides with the same sequence but without m^6^A modifications (oli1-nmod, oli2-nmod). Remarkably, all but one (position A9400) of the methylated As were identified (Figure 2A,B, blue bars). The m^6^A site at position A9400 is located too close (9nt) to the 5’ end of oli2-mod, which prevents its accurate detection ^24^. The methylated A8563, A9428 and A9442 positions were detected with a methylation rate of ∼100% in both oligomers (Figure 2A,B). The lower methylation rate (∼75%) observed for A8532 is probably due to its position close (13nt away) to the 5’ end of oli1-mod. We note that in these oligonucleotides, no other A was detected as significantly methylated in modified and non-modified oligos (Table S5). These observations confirm the specificity and high level of efficiency of m^6^A detection with the RNA004 chemistry and the m^6^A base-calling option. We further assessed the quality of m^6^A detection by treating oli1-mod and oli2-mod harboring m^6^A modifications with the ALKBH5 and FTO demethylases and then performing the detection process. Methylation rates were significantly lower at all m^6^A sites and no other A was detected as significantly methylated after treatment of modified and non-modified oligos, confirming the quality of the m^6^A detection process (Figure 2A,B; Table S5). We then assessed the linearity of the measurement of the methylation rates. Different mixtures of modified and non-modified RNA oligonucleotides (oli-mod and oli-nmod) were prepared and sequenced (Materials and Methods). The measured methylation rates were compared with the rates expected for these mixtures, for the five positions A8436, A8442, A9163, A9428 and A9442, showing the linearity of the rate measurement in the full range of values, from non-methylated (<3%) to fully methylated sites (>96%) (Figure S2).

**Figure 2.**
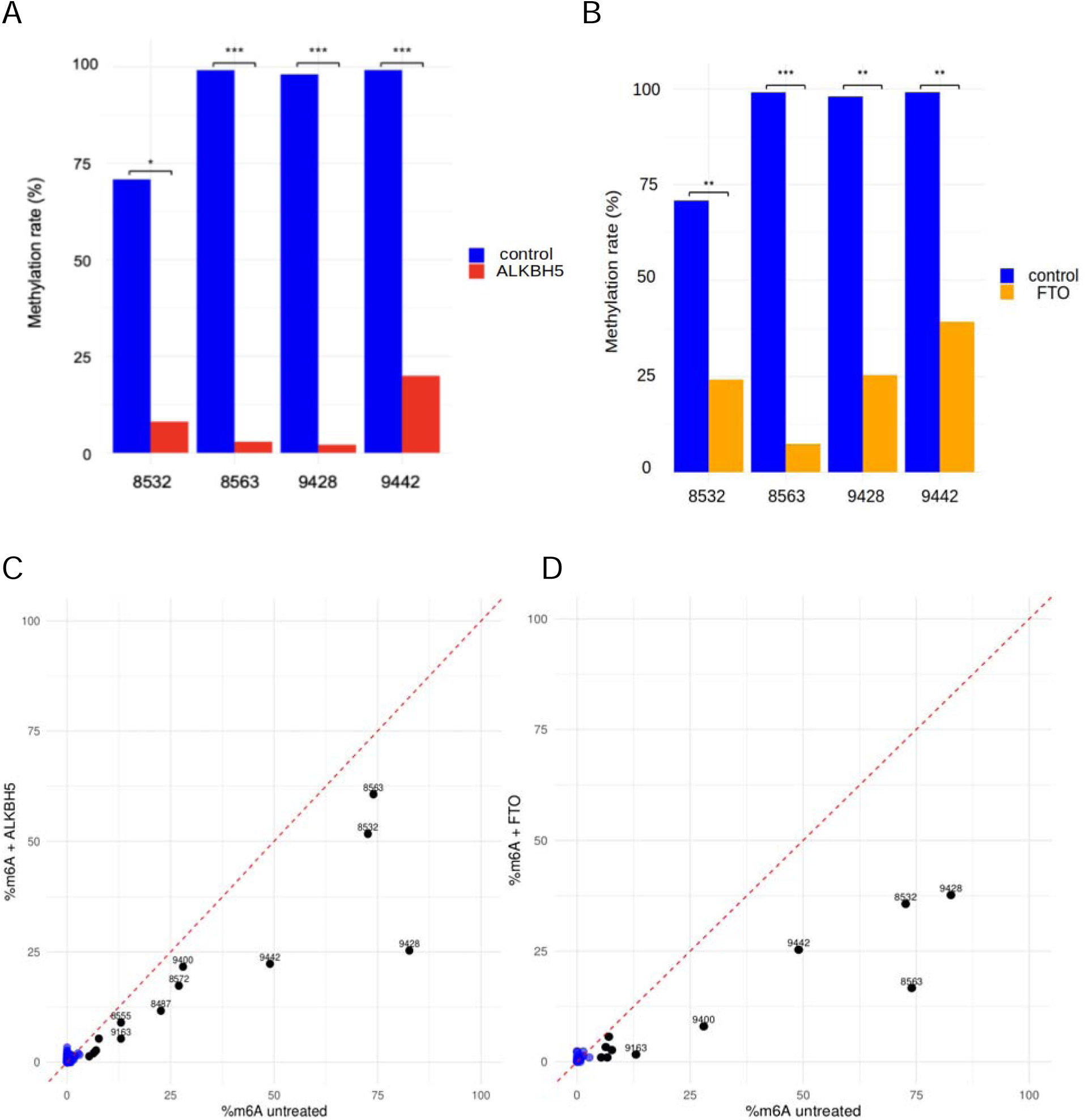
(A, **B)** Efficiency of m^6^A site detection performed with methylated synthetic oligonucleotides. In blue, methylation rates measured at the indicated m^6^A positions, comparing unmodified (oli1-nmod and oli2-nmod) and m^6^A-modified (oli1-mod and oli2-mod) oligonucleotides. In red or orange, the same measurement after the demethylation of synthetic m^6^A-modified oligonucleotides with ALKBH5 or FTO, respectively (Materials and Methods). *, p-value <10^-6^; ***, p-value <10^-149^. **(C, D)** Validation of m^6^A detection in the SupT1_20h_pi rep2 sample. In abscissa, methylation rate without treatment; in ordinate, methylation rate after treatment with the indicated demethylase. Black points represent the 14 detected m^6^A sites, blue points represent all other As in the reference genome; negative values are set to zero; in both panels we plotted only the positions covered by >100 reads and >40 modified reads.

### Validation of the candidate m^6^A sites with the ALKBH5 and FTO demethylases

For further validation of the detection of m^6^A sites in viral RNAs from HIV-1-infected cells, SupT1 cells infected for 20 h were treated with two demethylases (ALKBH5 for the SupT1_20h_pi rep1 and rep2 samples, and FTO for the SupT1_20h_pi rep2 sample), sequenced and the results compared with those of the untreated samples. Remarkably, all 14 sites had significant lower methylation rates after treatment with both demethylases except for positions A8487 and A8555 that presented low coverage after FTO treatment (Table S6). Despite degradation due to the demethylase treatment (Table S1), the most abundant isoforms could be used to validate most sites (Table S7). Interestingly, Note that ALKBH5 and FTO demethylases had lower efficiency with HIV-1 RNAs (Table S6) than with RNA oligonucleotides (Table S5). This difference is likely due to conformational differences presented by a RNA sequence when embedded either in oligonucleotides or in native HIV-1 transcripts, that may ultimately impact the demethylase activity ^35^. Overall, these results, obtained with synthetic oligonucleotides and viral mRNA from HIV-1 infected cells, highlight the quality of our detection method and confirm that the 14 m^6^A methylated sites identified in HIV-1 transcripts are *bona fide* m^6^A modification sites.

### Detection of m^6^A sites in HIV-1 splicing isoforms

We then investigated the distribution of m^6^A sites in each of the different splicing isoforms. The 14 m^6^A sites found in the pool of all transcripts were also detected in each of the 51 isoforms identified in the six samples (Figure 3A, Table S8, Figures S3, S4). The methylation rate of these sites was similar to that for all transcripts, characterized by cluster 1 centered around the two highly methylated A8532 and A8563 sites and cluster 2 centered around the highly methylated A9428. An unsupervised clustering of viral transcripts based on the methylation rates observed at the 14 m^6^A sites showed that the isoforms clustered into three groups. The first group corresponded to the US transcripts. The second group consisted mostly of PS transcripts and the third group consisted of the CS transcripts (Figure 3A, Figure S4). Methylation rates differed significantly between these classes and were highest for CS and lowest for US (CS rates > PS rates > US rates, Figure 3B, Figure S5). These results are consistent with previous observations (CS rates > PS rates > US rates) ^15,19,25^.

In parallel to the 14 sites common to all viral mRNA isoforms, we detected four additional m^6^A sites specific to only a few isoforms. Three of these sites, A5613, A5654 and A5703, were detected in the Vif and Vpr coding regions (between the D3 and A3 splice sites) and are specific to the isoforms encoding Vif and Vpr. The fourth site, A5887, was detected in the Tat coding region (between A3 and A4c) and is specific to the Vif, Vpr, Tat and SORF1 isoforms (Figure 4A, Table S9). The size of the corresponding RNAs depends strongly on the absence (CS, -i4) or presence (PS, +i4) of intron 4 (i4: D4A7, 2325 nt-long). As for the global trend observed for the 14 sites common to all isoforms presented in Figure 3, the methylation rates of these four additional m^6^A sites was higher for the CS (-i4) than for the PS (+i4) isoforms, but this difference was more pronounced (Table S9). For example, in the SupT1_20h_pi (rep1 and rep2) samples, the methylation rates of the four sites in the Vif1 isoform (-i4) were 3 to 9 times higher than those in the Vif2 isoform (+i4) (Table S9). Similarly, the methylation rates in Vpr1 and Vpr2 were much higher than those in Vpr3 and Vpr4, respectively. There was one notable exception to this tendency: the rate of A5887 methylation was significantly higher in Tat 5 (PS, +i4) than in Tat1 (CS, -i4) (Figure 4B). Importantly, all four sites were unmethylated or had very low rates of methylation in unspliced transcripts (≤ 1.3%). A similar trend was observed for the other four SupT1 and CD4 samples, for the CS and PS transcripts of the Vif, Vpr and Tat isoforms (Figure S6, Table S9).

**Figure 3.**
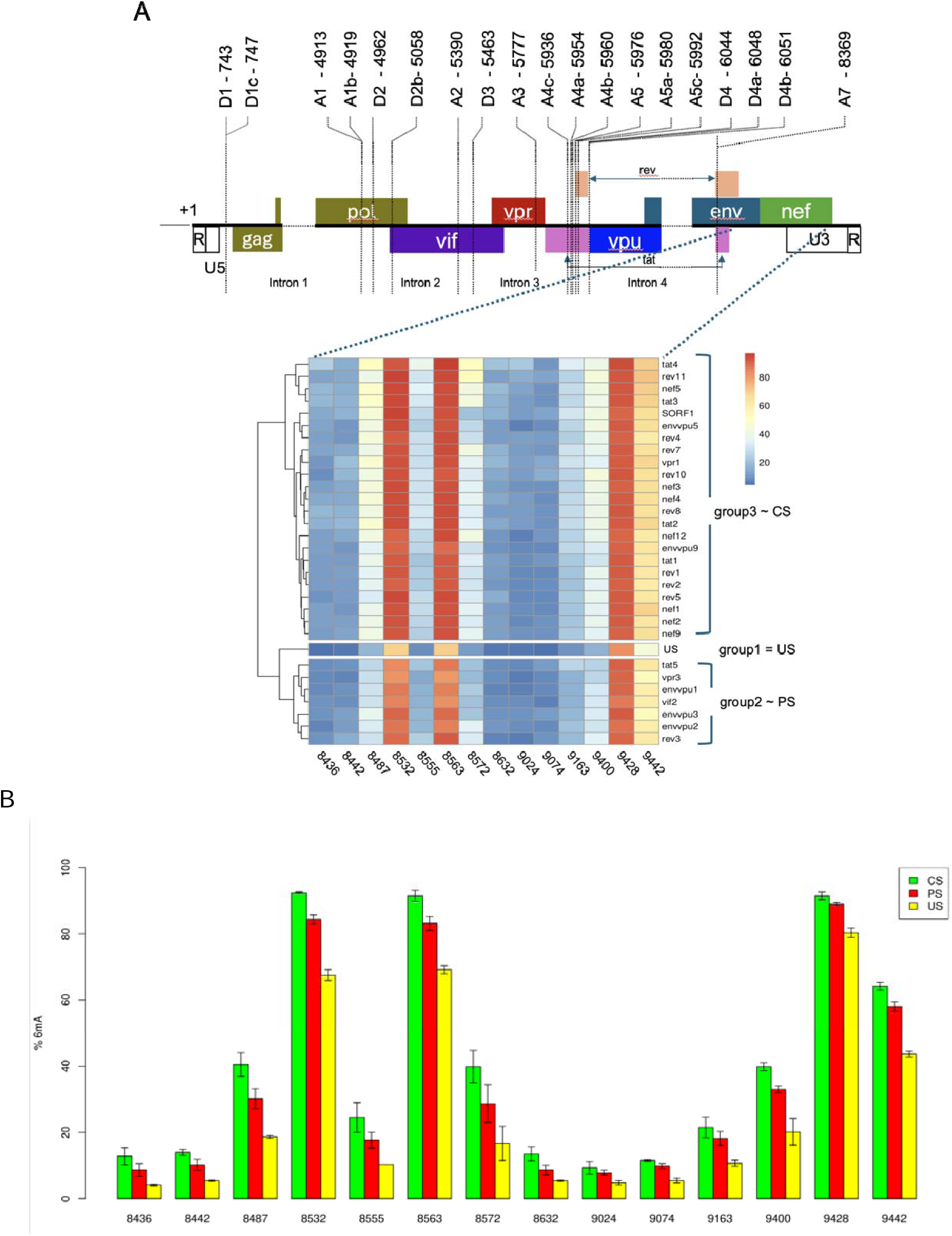
Methylation rates for the 14 m^6^A sites detected in SupT1 cells_20h_pi. **(A)** Unsupervised heatmap of the methylation rates of HIV-1 transcripts (Materials and Methods) in SupT1_20h_pi rep1. Isoforms are clustered based on the methylation rates of the 14 m^6^A sites. The clusters essentially correspond to US, PS and CS transcripts. **(B)** Methylation rates of m^6^A sites in the CS, PS and US transcripts of the SupT1_20h_pi rep1 and rep2 (mean values ± SD).

**Figure 4.**
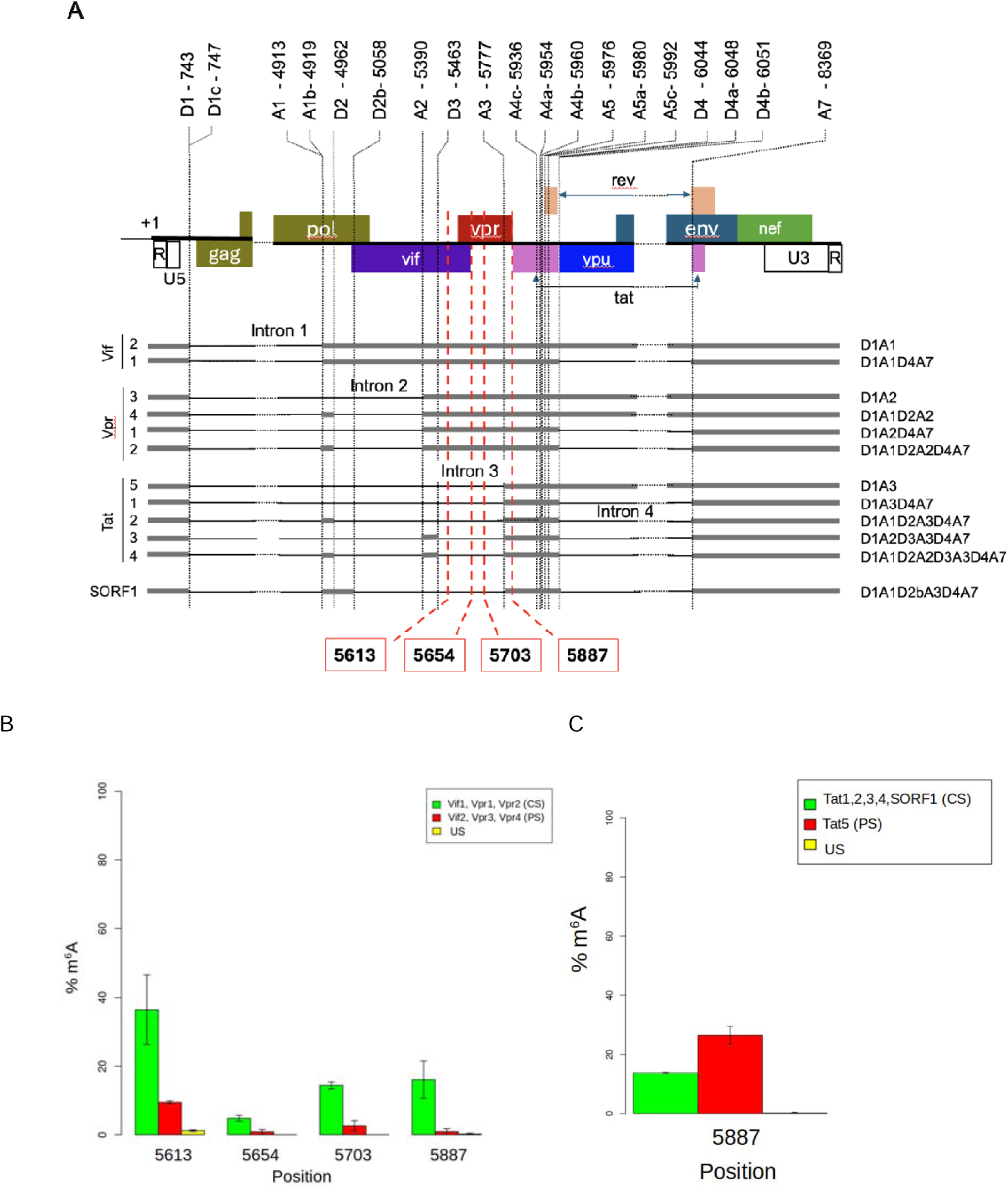
Position and methylation rates of the four isoform-specific m^6^A sites on SupT1_20h_pi rep1 and rep2. **(A)** The A5613, A5654 and A5703 sites are common to the Vif and Vpr isoforms; A5887 is common to the Vif, Vpr, Tat and SORF1 isoforms. **(B)** Methylation rates in SupT1_20h_pi rep1 and rep2, calculated for the four sites in the CS (- intron 4, Vif1, Vpr1, Vpr2), PS (+ intron 4, Vif2, Vpr3, Vpr4) and US isoforms (mean ± SD). **(C)** Methylation rates calculated for the A5887 site for the CS (Tat1, 2, 3, 4, SORF1), PS (Tat5) and US isoforms (mean ± SD).

### Single-molecule analysis of m^6^A sites

We investigated the distribution of the methylated m^6^A sites in individual transcripts by counting, for US, PS and CS transcripts, the numbers of methylated As in the eight sites of cluster 1 (from 0 to 8 methylated As) (Figure 5A, S7A,C) and the three sites of cluster 2 (from 0 to 3 methylated As) (Figure 5B, S7B,D) of the 3’UTR region of HIV-1, as defined in Figure 1. We examined the two clusters either separately (Figure 5A, 5B, Figure S7) or simultaneously on each read (Figure 5C-E, Figure S8, Table S10). The distribution of m^6^A sites was similar for rep1 and rep2 samples, for the SupT1 samples at 20 h and 30 h post-infection, and also in SupT1 and primary CD4^+^ T cells. We found that the number of sites modified per molecule increased from US to PS to CS. When considering the two clusters simultaneously, we observed that the peaks for the US transcripts corresponded to two methylated sites in cluster 1 and one or two methylated sites in cluster 2 (Figure 5A,B, Figure S7). By contrast, for the PS and CS transcripts, the peaks corresponded to two or three methylated sites in cluster 1 and two methylated sites in cluster 2. In the six samples, a small proportion (3.7% to 5.4%) of US transcripts were totally unmethylated simultaneously in the two clusters. The proportion of unmethylated transcripts decreased to 1.5% for PS transcripts and 0.3% for CS transcripts (Table S10).

**Figure 5.**
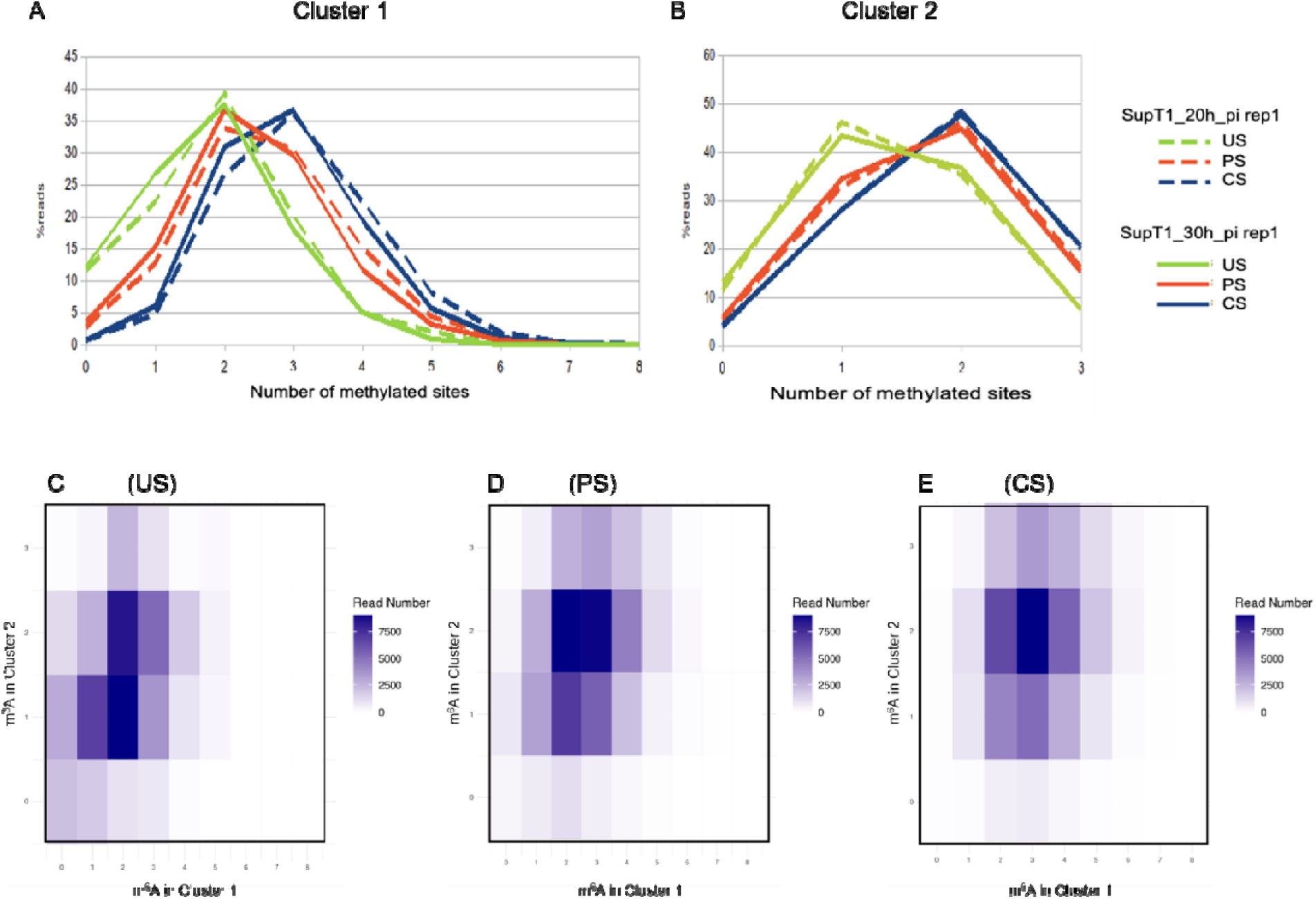
Single-molecule analysis of methylated m^6^A sites. **(A)** Cluster 1. **(B)** Cluster 2. **(C-E)** Correlation between the number of methylated sites in cluster 1 and in cluster 2 of the same transcript from the SupT1_20h_pi rep1 sample. The intensity of the color in the heatmaps indicates the number of reads with the indicated numbers of methylated As in cluster 1 (0 to 8 m^6^As) and cluster 2 (0 to 3 m^6^As) (Materials and Methods).

### Detection and analysis of 2-LTR transcripts

During the analysis of the splicing isoforms, we identified a class of RNA transcripts mapping to the long terminal repeats (LTR, constituted by the U3-R-U5 segments) located at either end of the viral genome. They result from the transcription of 2-LTR circles, a non-integrated circular form of the reverse-transcribed viral DNA formed early after infection during the conversion of the viral genome into DNA by the junction of the two LTRs located at the ends of the genome ^36^ (Figure 6A, 6B). They begin at the promoter at the 5’ end of the R segment of the 3’ LTR and terminate at the polyA site at the 3’ end of the R segment of the 5’ LTR. They create a precise join between the 3’ end of the RU5 segment of the 3’ LTR and the 5’ end of the U3R segment of the 5’ LTR. Studies of the 2-LTR U5-U3 junction (the junction between the two LTRs) have shown this junction to be highly heterogeneous in infected cells and between infected individuals, frequently due to insertions ^37,38^. Transcripts encompassing this junction have been detected by RT-PCR, but were not precisely identified. A recent study of the HIV-1 transcriptome by Nanopore cDNA sequencing identified RNA species that could correspond to 2-LTR transcripts and were detected in infected CD4^+^ T and HeLa cells ^27^. Here, the 2-LTR transcripts were detected in both infected SupT1 and infected primary CD4^+^ T cells (0.16% to 1.69 % of HIV-1 transcripts, Table S2B). They displayed either the precise U5-U3 junction or a number of variants mostly carrying short (1-nt to 10-nt) insertions in agreement with previous studies ^37,38^. The detection of m^6^A sites in these 2-LTR transcripts revealed the presence of six sites that were detected in all six samples (Figure 6B-D, Tables 2, S11). The first site, 2L125, is homologous to the A578 position of the viral genome (Figure 6B,C) common to all HIV-1 transcripts. Remarkably, this 2L125 site was methylated in ∼33 % of 2-LTR transcripts (Table 2) but was not detected as methylated in any other HIV-1 transcript, in all six samples. The other five sites are located in the U3 segment of the 5’ LTR and are homologous to m^6^A sites A9074, A9163, A9400, A9428 and A9442, which are common to all splicing isoforms (Figure 6B-D). Remarkably, in the 2-LTR transcripts, the methylation rates of all five of these sites were higher than those of the homologous sites in the viral genome. For example, the 2L272 site has a methylation rate of 29.2 ± 2.1 % in SupT1_20h_pi samples (Table 2), whereas the homologous site A9163 had a rate of 15.15 ± 3 % in the corresponding samples (Table 1). We note that the sequence variants observed at the position 2L183 (U5/U3 junction) may alter the methylation rate. The rate measured here is global and includes the rates of all variants. Future studies will be required to classify these variant sequences and to determine to what extent the methylation of this position differs with each sequence context.

**Figure 6.**
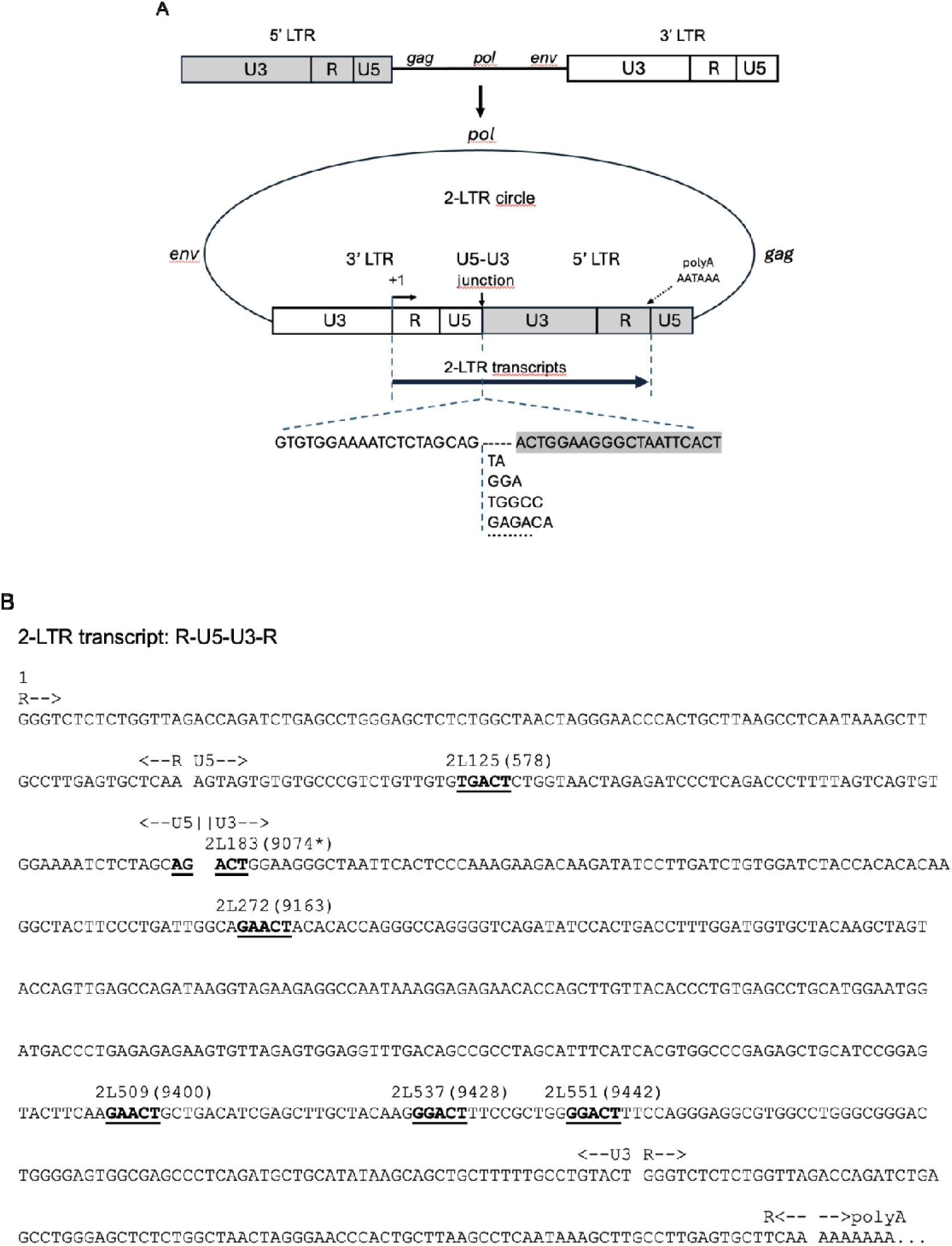
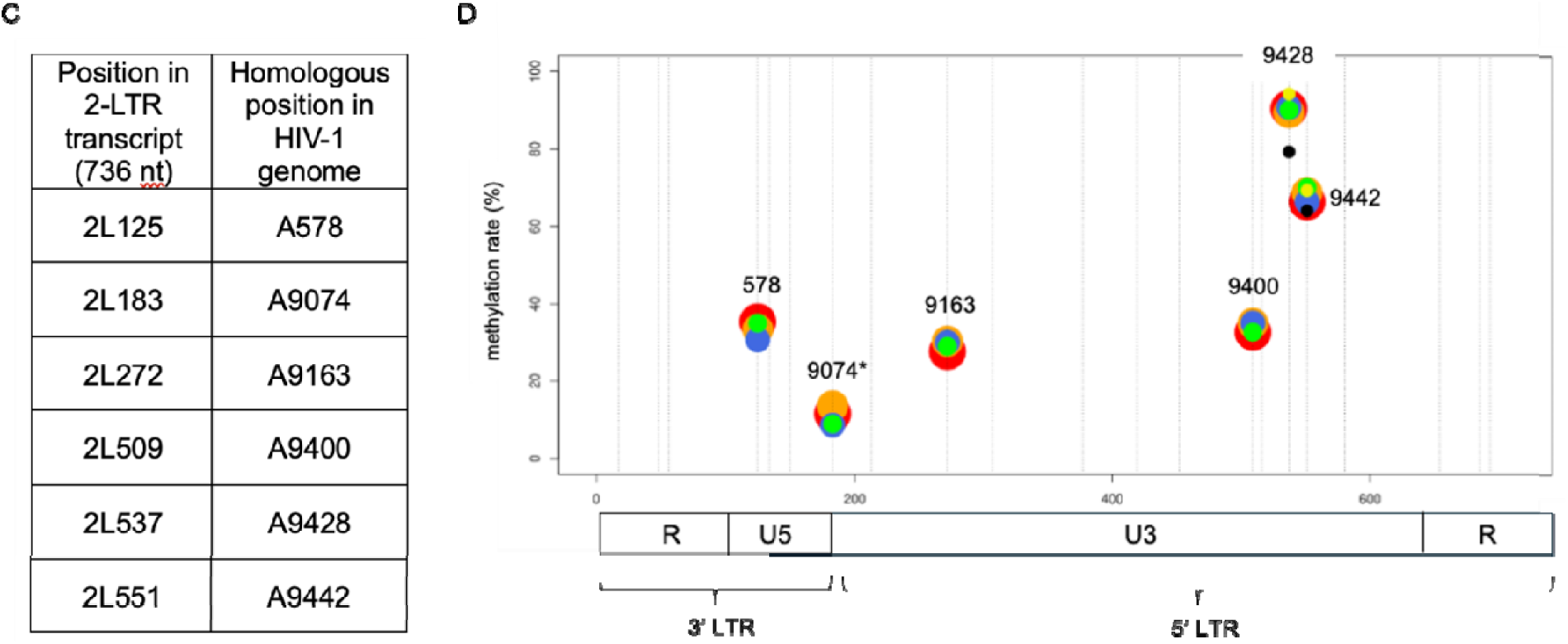
Transcription and methylation of the 2-LTR transcripts. **(A)** Diagram of the 2-LTR circles and of the transcripts joining the RU5 segment of the 3’ LTR to the U3R segment of the 5’ LTR. Several examples of insertion variants at the U5-U3 junction are shown. **(B)** Sequence of the 2-LTR transcripts. The successive transcribed regions — R, U5, U3, R — are indicated; m^6^A sites are shown in bold typeface and underlined and are named according to the position of the methylated A in the 2-LTR transcript; in brackets, the homologous position in the HIV-1 reference genome; for example, for 2L125(578), 125 is the position of the m^6^A in the 2-LTR transcript and (578) is the homologous position in the HIV-1 reference genome; * indicates that the sequence context of 2L183 in the 2-LTR transcript (AG<u>A</u>CT, see B) differs from that at the homologous position 9074 in the reference genome (GG<u>A</u>CT). **(C)** Correspondence table for a position in the 2-LTR transcript and the homologous position in the reference genome. **(D)** Sites detected in the 2-LTR transcripts of infected SupT1 and CD4^+^ T-cell samples (Materials and Methods). Large red circles, SupT1_20h_pi rep1; orange circles, SupT1_30h_pi rep1; blue circles, SupT1_20h_pi rep2; green circles, SupT1_30h_pi rep2; yellow circles, CD4_48h_pi; black circles, CD4_72h_pi. Vertical lines indicate the positions of the As in all the DRACH motifs (21) of the transcript.

**Table 1.** Methylation rates of the 14 m^6^A sites detected in the 3’ region (8360-9700) common to all HIV-1 transcripts measured in SupT1_20h_pi and SupT1_30h_pi samples (mean of rep1 and rep2 ± SD). The underlined nucleotides are single-nucleotide variants of the consensus DRACH sequence.

| m<sup>6</sup>A position | Sequence | SupT1_20h_pi | SupT1_30h_pi |
| --- | --- | --- | --- |
| A8436 | AGACA | 8 ± 2.4 | 6.85 ± 1.6 |
| A8442 | AGACA | 8.65 ± 2.3 | 8.2 ± 2.1 |
| A8487 | GGAC<u>U</u> | 26.35 ± 5.2 | 24.5 ± 3.5 |
| A8532 | AGACU | 76 ± 4.7 | 75.65 ± 5.2 |
| A8555 | GGA<u>U</u>U | 16.15 ± 4.5 | 15.15 ± 3.0 |
| A8563 | GAACU | 76.65 ± 3.8 | 76.2 ± 4.9 |
| A8572 | GGAC<u>G</u> | 25.35 ± 2.3 | 24.65 ± 1.9 |
| A8632 | GAACU | 8.5 ± 2.6 | 8 ± 1.8 |
| A9024 | UGACU | 6.3 ± 1.4 | 5.5 ± 0.7 |
| A9074 | GGACU | 8.2 ± 0.7 | 8 ± 2.4 |
| A9163 | GAACU | 15.15 ± 3.0 | 13.35 ± 2.3 |
| A9400 | GAACU | 28.85 ± 1.2 | 28.5 ± 4.9 |
| A9428 | GGACU | 83.5 ± 1.1 | 82.5 ± 2.5 |
| A9442 | GGACU | 51.15 ± 3.0 | 49.15 ± 4.5 |

**Table 2.** Methylation rates of the six m^6^A sites detected in the 2-LTR transcripts measured in SupT1_20h_pi and SupT1_30h_pi samples (mean of rep1 and rep2 ± SD).

| m<sup>6</sup>A position | Sequence | SupT1_20h_pi | SupT1_30h_pi |
| --- | --- | --- | --- |
| A578 (2L125) | UGACU | 33 ± 3,3 | 34,15 ± 1,2 |
| A9074 (2L183) | AGACU | 10,35 ± 1,9 | 11,35 ± 3,3 |
| A9163 (2L272) | GAACU | 29,2 ± 2,1 | 29,7 ± 0,9 |
| A9400 (2L509) | GAACU | 35 ± 3,3 | 33,85 ± 1,6 |
| A9428 (2L537) | GGACU | 90,8 ± 0,7 | 89,65 ± 0,5 |
| A9442 (2L551) | GGACU | 66,5 ± 0,3 | 69,35 ± 0,9 |

To further validate the m^6^A sites in 2LTR transcripts, m^6^A detection has been performed in the sample SupT1_20h_rep2 treated by ALKBH5 and FTO. This analysis used the *in vitro* genomic RNA as control sequence (Table S11) although it differs from the 2LTR sequence. To take this difference into account, a new analysis has been performed using as in vitro control a 2LTR RNA transcribed in vitro (638 pb, Materials and Methods). The results obtained in this second analysis (Table 3) are very similar to those obtained when using the *in vitro* genome sequence as control.

**Table 3.**
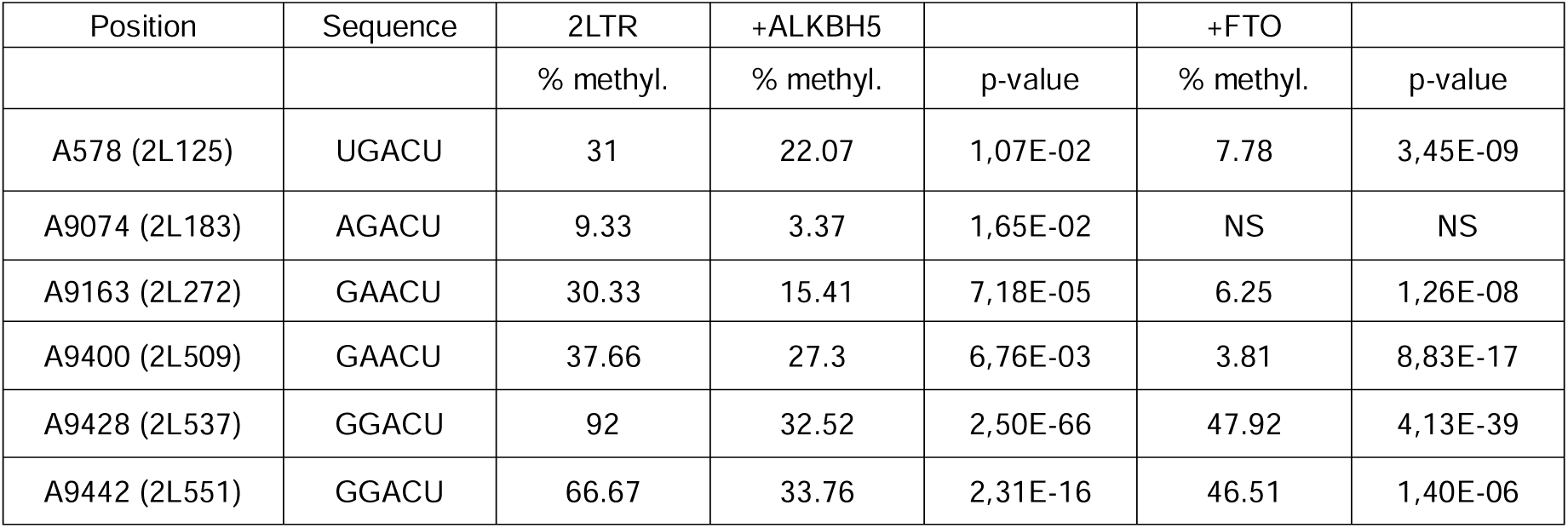
Demethylation and detection m6A on 2LTRs of SupT1_20h_pi rep2 sample. NS corresponds to a “non-significant value” equivalent to a total read count of less than 100.

## DISCUSSION

We report here the detection of m^6^A nucleotides in HIV-1 transcripts from infected human cells, with comparison to previous studies ^15,18–20,25^. We included cells of two different origins: CD4^+^ T cells from the SupT1 cell line and CD4^+^ T cells from the peripheral blood mononuclear cells (PBMC) of blood donors. These cells, which are natural HIV-1 targets, were infected for different periods of time (20 h and 30 h for SupT1 cells; 48 h and 72 h for blood donor cells). Using the new RNA004 chemistry and a new base-calling model, and stringent validation with two demethylases (ALKBH5 and FTO), we detected 14 m^6^A sites common to the unspliced HIV-1 RNA and to all the isoforms encoding viral proteins. Surprisingly, however, we found no detectable differences in the overall m6A pattern between SupT1 cells and primary CD4=:J T cells. This unexpected similarity reinforces the value of the SupT1 cell line as a robust and physiologically relevant model for studying HIV-1 RNA epitranscriptomic regulation.

The m^6^A positions were clustered in two regions of the 3’ end of the HIV-1 genome, both having a methylation profile featuring highly methylated central sites. These sites are partly included in the methylated regions identified in previous studies based on the immunoprecipitation of m^6^A-containing RNA (reviewed in^9^) or on m^6^A-SAC-seq ^39^. Unlike these studies, we found no methylation sites in the 5’ region of the reference HIV-1 genome, either in the unspliced RNA or in the diverse splicing isoforms. Overall, we observed a remarkable specificity of the 14 m^6^A sites in the 245 DRACH sites of the HIV-1 genome. The saturation of the five DRACH sites located in the 200 nt cluster 1 and the additional methylation of three single-nucleotide DRACH variants in the same region further increase the specificity of HIV-1 methylation. However, despite the proposal of hypotheses involving the recruitment of m^6^A writers by RNA-binding proteins or by RNA polymerase II close to DRACH sites during transcription ^6,40^, there is currently no explanation for the specificity of the methylation of some DRACH sites but not others. Interestingly, on examination of the secondary structure of the HIV-1 RNA, most of the 14 m^6^A sites were found to be located in single-stranded portions of the structure, or within the final nucleotides at the 3’ end of a helix (Figure 7). This finding is fully consistent with studies showing that DRACH sequences located in a double-stranded RNA duplex prevent efficient m^6^A methylation by the METTL3–METTL14 complex ^41^.

**Figure 7.**
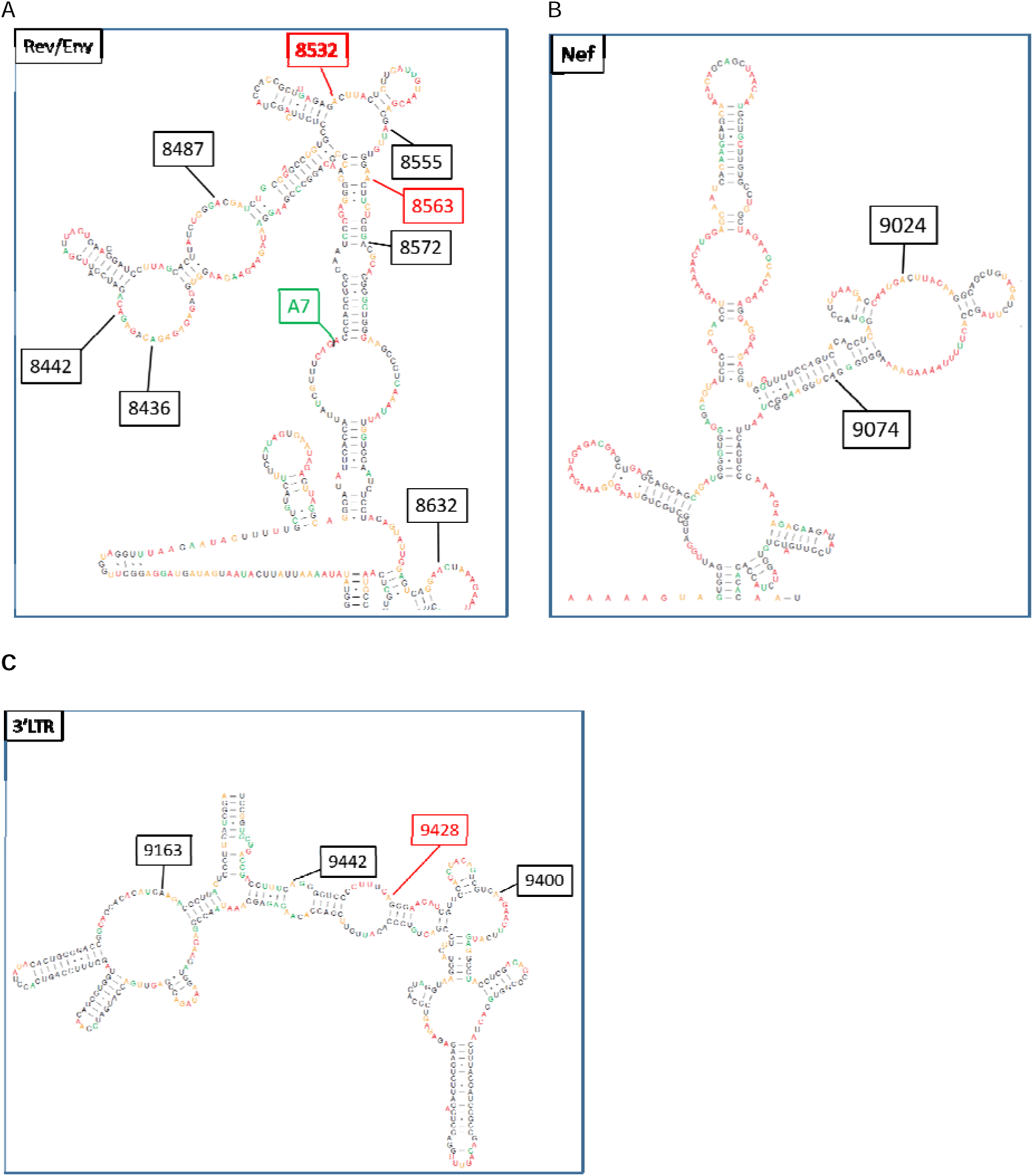
Secondary structures of HIV-1 RNA around the 14 m^6^A sites detected in all isoforms. **(A)** Region of the Rev/Env sequence containing the eight m^6^A sites of cluster 1; this structure is common to the US and PS transcripts as both retain the end of the unspliced intron 4 at the A7 acceptor site, in green (intron 3’ end). In red, the two highly methylated sites (64-80%). **(B)** Region in the Nef sequence containing the methylated sites A9024 and A9074. **(C)** Region in the 3’LTR containing the three m^6^A sites of cluster 2. In red, the highly methylated site (79-84%). The structure predictions are adapted from ^42^.

The 14 m^6^A sites include three sites (A8532, A9428, A9442) previously detected as peaks in RNA immunoprecipitation experiments ^18,19^ and as methylated nucleotides (m^6^As) by direct RNA sequencing ^25^. This last study also identified A8563 as a weakly methylated m^6^A site, contrasting with our results, as A8563 was one of the highly methylated sites of cluster 1 (Figure 1). We found that the 14 m^6^A sites had higher methylation rates in CS isoforms than in PS isoforms and US transcripts (Figure 3B, Figure S5). At single-molecule level, we observed that most CS transcripts had three methylated sites in cluster 1, and two in cluster 2. By contrast, most US transcripts had two methylated sites in cluster 1, and one in cluster 2. In addition, only very small proportions of CS transcripts had no methylated site in either cluster (0.3%). This proportion increased only slightly, from 3.7% to 5.4%, in US transcripts. These observations are consistent with a previous study ^25^ of three methylation sites (A8532, A9428, A9442) reporting that 2.5% and 8.5% of CS and US transcripts, respectively, were unmethylated. These differences emphasize isoform-dependent variation in methylation, although a direct causal role in splicing regulation would require further investigations.

Four additional sites were detected specifically in several isoforms (Figure 4A, Tables 2, S9). The methylation rates of these additional sites were higher for CS than for PS isoforms, but the ratio of CS transcript methylation to PS transcript methylation was much higher than for the 14 sites common to all isoforms (Figure 4B and Figure 3B). This suggests that m^6^A sites located at different positions within a specific isoform may be involved in different processes, such as translation or the maintenance of stability. Further experiments involving the mutation of one or several of these sites would help to address these questions. Interestingly, we observed that the two demethylases (ALKBH5 and FTO) exhibited different specificities depending on the RNA substrate (Figure 2C,D). Although both treatments led to substantial RNA degradation, sequencing coverage at potential m^6^A sites remained sufficient to reveal a significant reduction in methylation levels, thereby confirming the chemical nature of the modifications at these positions.

In addition to the splicing isoforms, we detected ≈736 nt-long RNAs resulting from the transcription of 2-LTR unintegrated DNA circles joining the R segments of the two LTRs (RU5-U3R, Figure 6A). These transcripts contained six m^6^A sites, five of which were common to all spliced isoforms and had higher methylation rates than any of the other transcripts. An additional site, 2L125, was identified in the U5 segment, with a methylation rate ranging from 24% to 35%. We observed that (i) positions homologous to this position (2L125) in the other transcripts were not considered methylated by our detection process and (ii) the other five sites had methylation rates higher than that at the homologous sites in all the other transcripts. These results strongly suggest that, despite not encoding canonical viral proteins, 2-LTR transcripts play a functional role in the viral life cycle. Consistent with this idea, 2-LTR circles were initially thought to be dead-end products in the viral life cycle ^43^, but many studies subsequently showed that they could be transcribed and might constitute a reserve supply of HIV-1 genomes for proviral integration ^44–48^. In addition, the large difference in 2L125 methylation rate between 2-LTR transcripts and the other transcripts (same sequence context) confirms that this site is not an artifact of the detection process but, indeed, a *bona fide* methylation site.

It is generally thought that methylation occurs cotranscriptionally and that exclusion of m^6^A from splice-site proximal regions by the exon junction complex results in low methylation rates close to splice junctions ^34,49–52^. However, our observation that CS transcripts have higher methylation levels than PS and US transcripts suggests that other mechanisms may be at play. Notably, recent studies have shown that m^6^A methylation can occur posttranscriptionally, especially in transcripts that are highly methylated ^53^. This raises the possibility that the methylated sites observed at highly specific positions within HIV-1 transcripts are methylated posttranscriptionally and are potentially associated with an epitranscriptomic role in the virus life cycle.

In conclusion, we provide an unbiased assessment of m^6^A modifications in HIV-1 transcripts isolated from infected CD4^+^ T cells. We reveal the existence of novel m^6^A sites in the 3’ end common to all transcripts, in specific isoforms, and in 2-LTR transcripts. Importantly, the use of two independent m^6^A-specific demethylases (ALKBH5 and FTO) provides strong validation of the existence of these modifications. Our study highlights isoform-dependent variation in m^6^A levels and confirms the selective nature of HIV-1 methylation. These results pave the way for efforts to decipher the role of m^6^A modifications in the HIV life cycle.

## DATA AVAILABILITY

Data are deposited at GEO NCBI database under the accession number GSE291168

## Supplementary Data Statement

Supplementary Data are available at NAR Genomics and Bioinformatics Online

## Author contributions

Conceptualization: ON, AM, CH; Formal analysis: DN, CT; Investigation: SB, EVD, LB, SG, CBR, RO, YJ; Funding acquisition: ON, AM, SB, CH; Writing original draft: DN, SB, AM, CT, CH, ON.

## ACKNOWLEDGMENTS

This work was granted by the Agence Nationale de Recherche sur le SIDA (ANRS-Maladies infectieuses émergentes) for fundings to A.M. and O.N., S.B salary was supported by ANRS. L.B PhD salary was supported by ANRS-MIE. This study was initiated thanks to fundings from the cross-disciplinary thematic axes of I2BC through the epiRNA programs. The HTS facility of the I2BC acknowledges the support of France Génomique (funded by the French National Program "Investissement d’Avenir" ANR-10-INBS-09).

