## supplemental data for "High-resolution HIV-1 m^6^A epitranscriptome reveals isoform-dependent methylation clusters and unique 2-LTR transcript modifications"

### SUPPLEMENTARY FIGURES

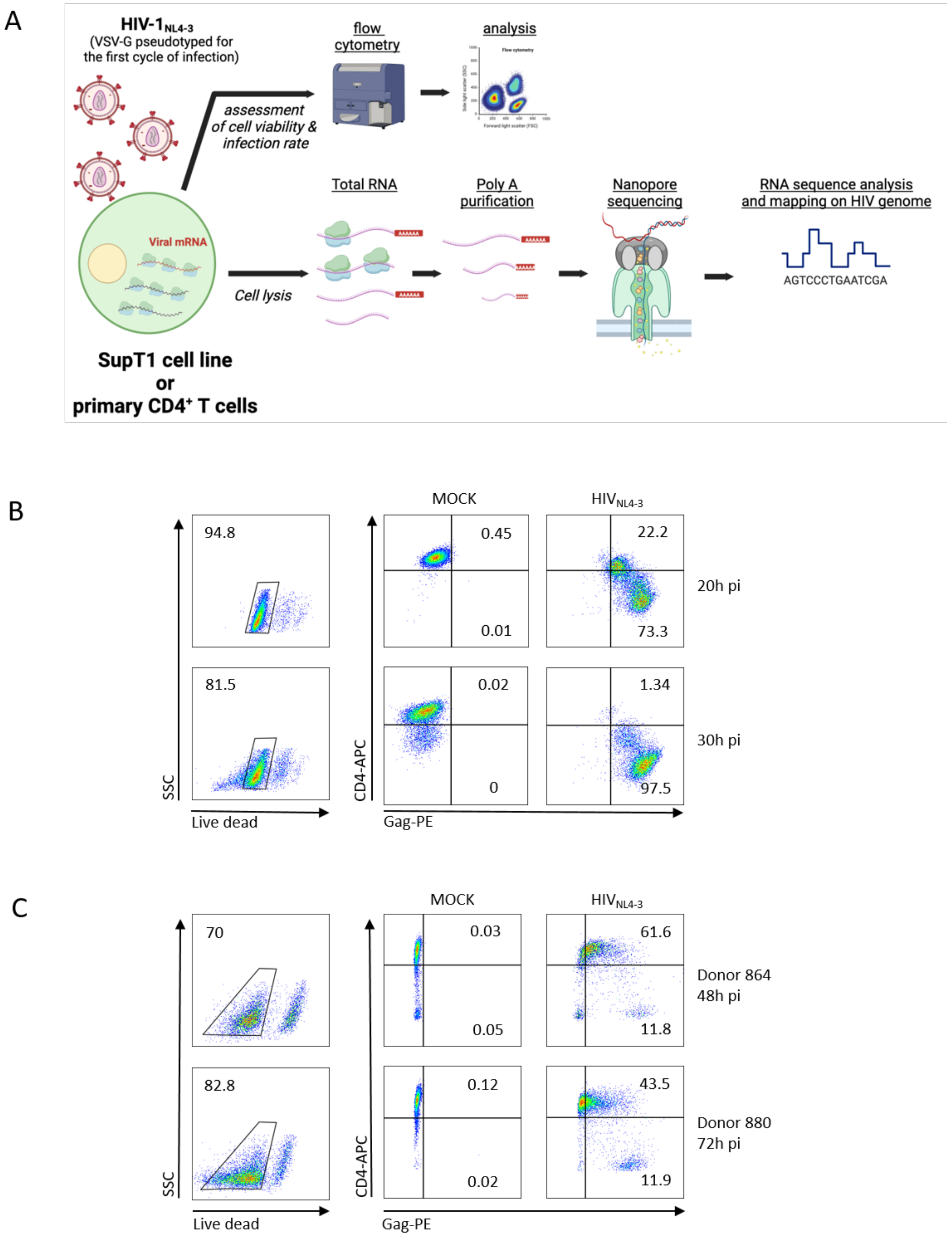

**Figure S1. (A) Outline of the infection and Nanopore sequencing protocols**. SupT1 T cells or primary CD4$^+$ T cells were infected with HIV$_{NL4-3}$ pseudotyped with VSV-G to enhance viral infection. At 20 or 30 hours post-infection (pi) for HIV $_{NL4-3}$-infected SupT1 cells and at 48 h or 72 h pi for HIV $_{NL4-3}$-infected primary CD4$^+$ T cells, we collected a fraction of the cell culture. The viability and percentages of HIV-Gag$^+$ and CD4$^+$ cells were assessed by flow cytometry. During HIV infection, cells start expressing HIV-Gag protein and progressively lose their membrane expression of CD4 due to expression of the HIV accessory proteins Nef and Vpu, and HIV-Env. The cells were harvested and polyA RNA was purified and sequenced with Nanopore technology. Reads were mapped onto the HIV$_{NL4-3}$ genome sequence. Created with BioRender. Moris, A. (2025) https://BioRender.com/o40u811 **(B) Analysis of the infection rate of HIV$_{NL4-3}$-infected CD4$^+$ SupT1 cells.** After 20 h or 30 h, cells infected with HIV$_{NL4-3}$ or mock-treated were labeled with a viability dye and stained with antibodies specific for CD4 and HIV-Gag. Left panels: the percentage viable cells. Middle and right panels: percentages of CD4$^-$HIV-Gag$^+$ and CD4$^+$HIV-Gag$^+$ cells among mock-treated and HIV$_{NL4-3}$-infected CD4$^+$ SupT1 cells, respectively. **(C) Analysis of the infection rate of HIV$_{NL4-3}$-infected primary CD4$^+$ T cells.** After 48 h or 72 h, cells from donors 864 and 880 were harvested and stained as in (B). Left panels: the percentage viable cells. Middle and right panels: percentages of CD4$^-$HIV-Gag$^+$ and CD4$^+$HIV-Gag$^+$ cells among mock-treated and HIV$_{NL4-3}$-infected primary CD4$^+$ T cells, respectively, from donors 864 (top panels) and 880 (bottom panels). Numbers indicated the percentage of cells within the quadrant. MOCK: cells mock-treated without infection used as a negative control.

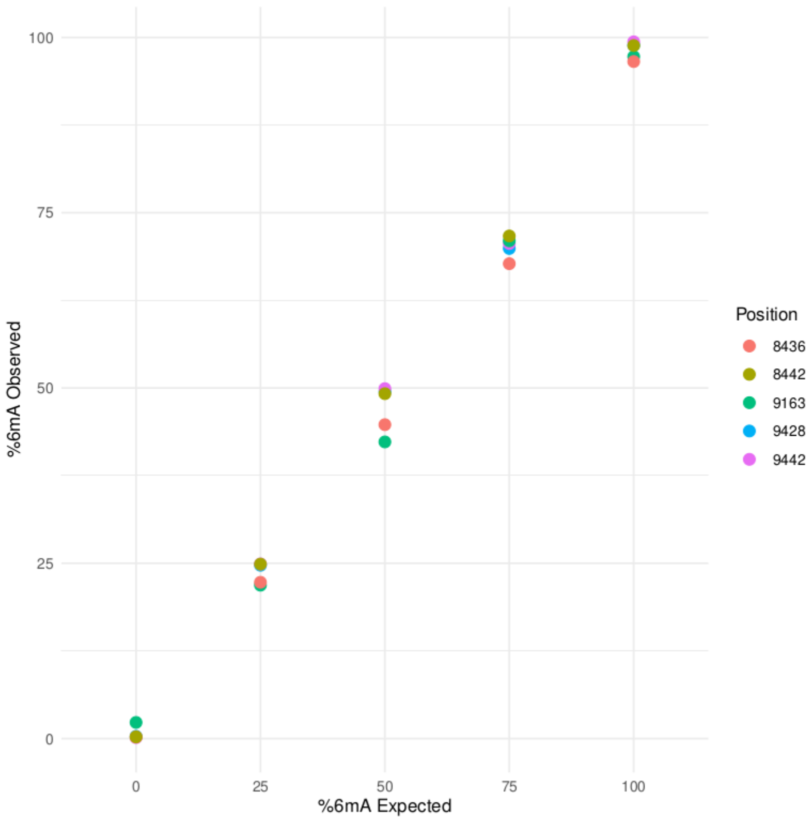

**Figure S2.** Linearity of the measurement of m$^6$A methylation rates. Different mixtures of modified and non-modified RNA oligonucleotides (oli-mod and oli-nmod) were prepared and sequenced (Materials and Methods) allowing to measure the methylation rates for the expected rates 0%, 25%, 50%, 75% and 100%, for the five positions A9428 and A9442 (oli1mod and oli1-nmod), A8436 and A8442 (oli3mod and oli3-nmod) and A9163 (oli4mod and oli4-nmod). The diagram shows the linearity of the rate measurement from non-methylated (<3%) to fully methylated sites (>96%); correlation between expected and observed rates, r = 0.998 (Pearson).

A

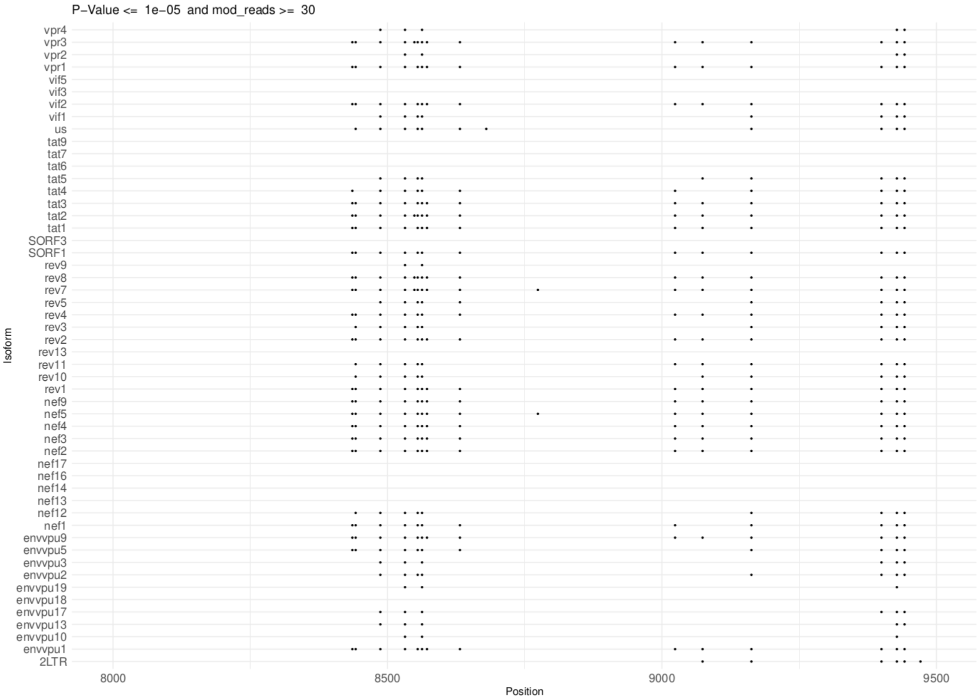
P−Value <= 1e−05 and mod_reads >= 30
Isoform
vpr4
vpr3
vpr2
vpr1
vif5
vif3
vif2
vif1
us
tat9
tat7
tat6
tat5
tat4
tat3
tat2
tat1
SORF3
SORF1
rev9
rev8
rev7
rev5
rev4
rev3
rev2
rev13
rev11
rev10
rev1
nef9
nef5
nef4
nef3
nef2
nef17
nef16
nef14
nef13
nef12
nef1
envvpu9
envvpu5
envvpu3
envvpu2
envvpu19
envvpu18
envvpu17
envvpu13
envvpu10
envvpu1
2LTR
8000
8500
9000
9500
Position

B

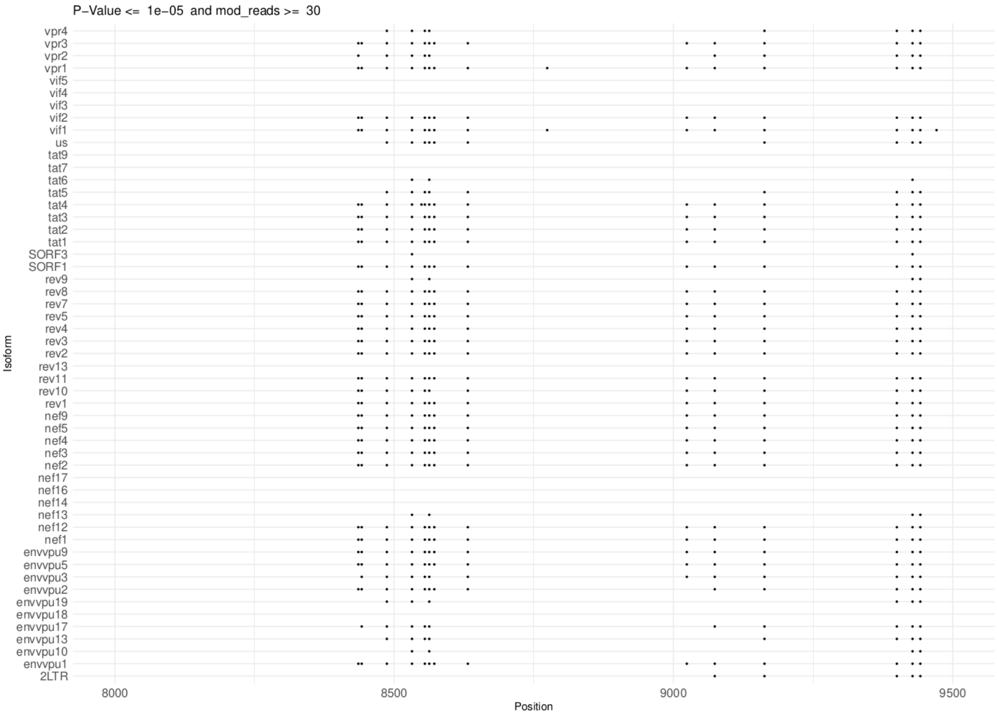
P−Value <= 1e−05 and mod_reads >= 30
Isoform
vpr4
vpr3
vpr2
vpr1
vif5
vif4
vif3
vif2
vif1
us
tat9
tat7
tat6
tat5
tat4
tat3
tat2
tat1
SORF3
SORF1
rev9
rev8
rev7
rev5
rev4
rev3
rev2
rev13
rev11
rev10
rev1
nef9
nef5
nef4
nef3
nef2
nef17
nef16
nef14
nef13
nef12
nef1
envvpu9
envvpu5
envvpu3
envvpu2
envvpu19
envvpu18
envvpu17
envvpu13
envvpu10
envvpu1
2LTR
8000
8500
9000
9500
Position

C

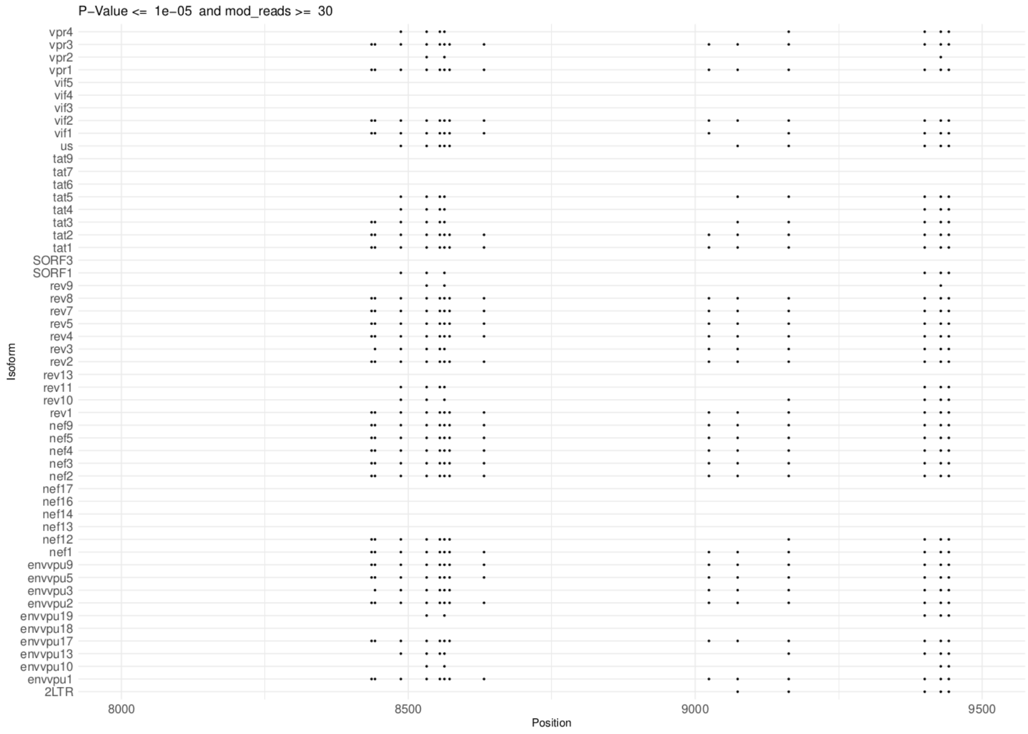
P−Value <= 1e−05 and mod_reads >= 30
Isoform
vpr4
vpr3
vpr2
vpr1
vif5
vif4
vif3
vif2
vif1
us
tat9
tat7
tat6
tat5
tat4
tat3
tat2
tat1
SORF3
SORF1
rev9
rev8
rev7
rev5
rev4
rev3
rev2
rev13
rev11
rev10
rev1
nef9
nef5
nef4
nef3
nef2
nef17
nef16
nef14
nef13
nef12
nef1
envvpu9
envvpu5
envvpu3
envvpu2
envvpu19
envvpu18
envvpu17
envvpu13
envvpu10
envvpu1
2LTR
8000
8500
9000
9500
Position

D

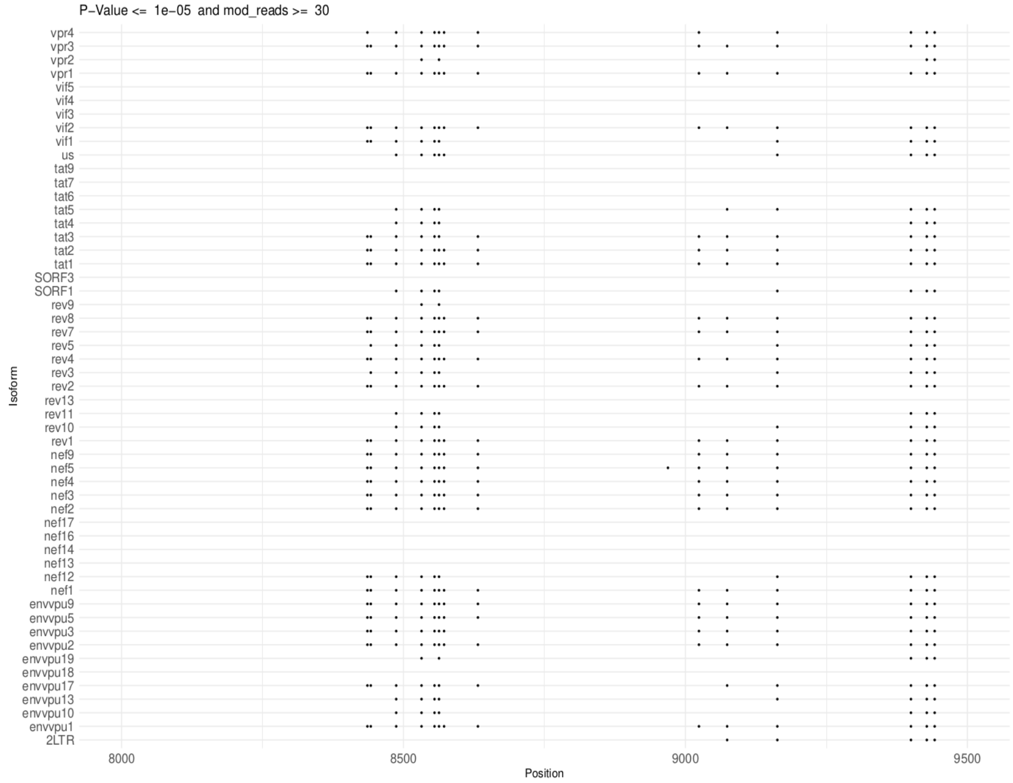
P−Value <= 1e−05 and mod_reads >= 30
Isoform
vpr4
vpr3
vpr2
vpr1
vif5
vif4
vif3
vif2
vif1
us
tat9
tat7
tat6
tat5
tat4
tat3
tat2
tat1
SORF3
SORF1
rev9
rev8
rev7
rev5
rev4
rev3
rev2
rev13
rev11
rev10
rev1
nef9
nef5
nef4
nef3
nef2
nef17
nef16
nef14
nef13
nef12
nef1
envvpu9
envvpu5
envvpu3
envvpu2
envvpu19
envvpu18
envvpu17
envvpu13
envvpu10
envvpu1
2LTR
8000
8500
9000
9500
Position

E

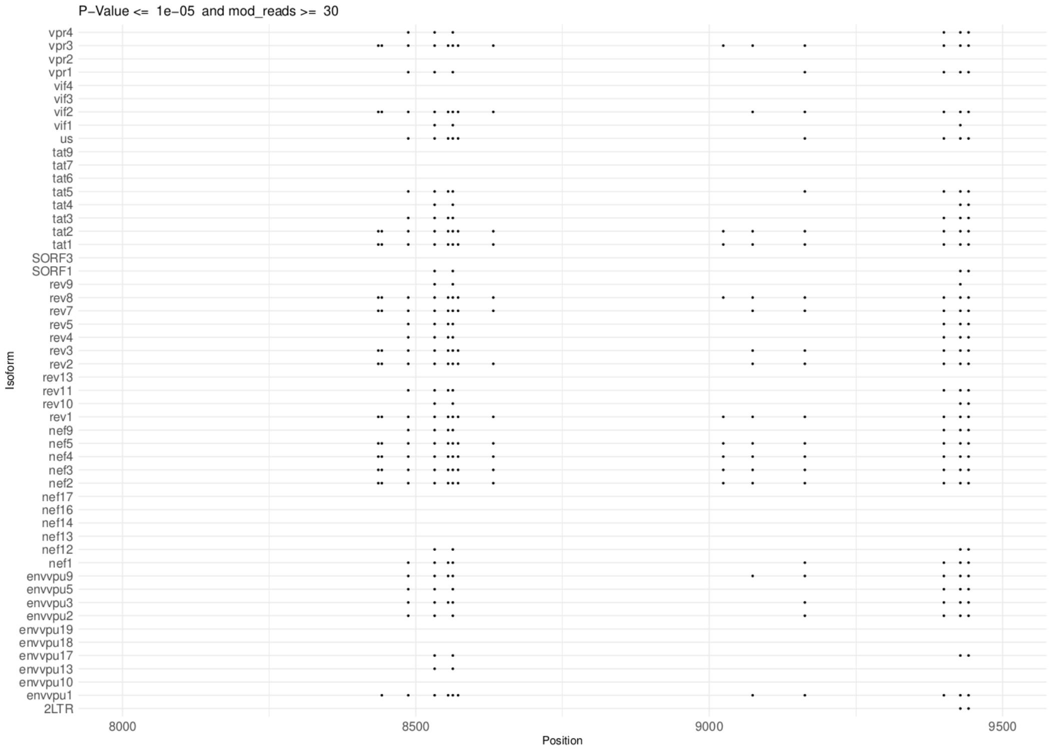

F

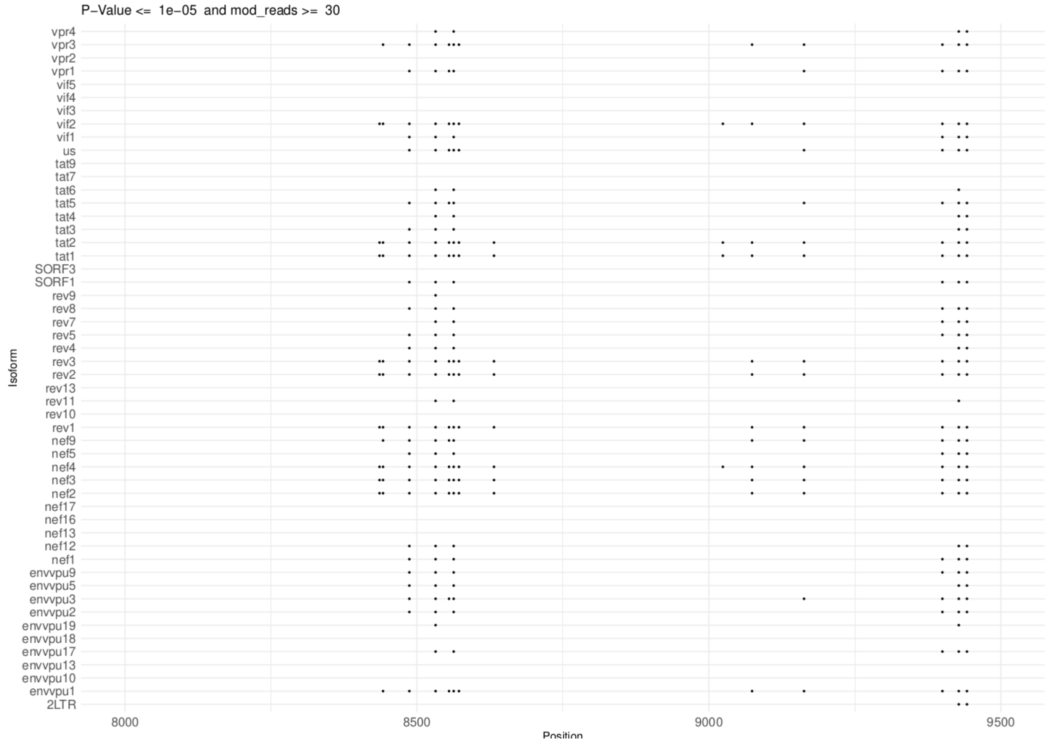

**Figure S3.** m$^6$A sites detected in splicing isoforms. **(A)** SupT1_20h_pi rep 1 sample; **(B)** SupT1_30h_pi rep1 sample; **(C)** SupT1_20h_pi rep2 sample; **(D)** SupT1_30h_pi rep2 sample; **(E)** CD4_48h_pi sample; **(F)** CD4_72h_pi sample. Only positions with *p*-values ≤ 1 x 10$^{-5}$ and a number of modified reads ≥ 30 are displayed.

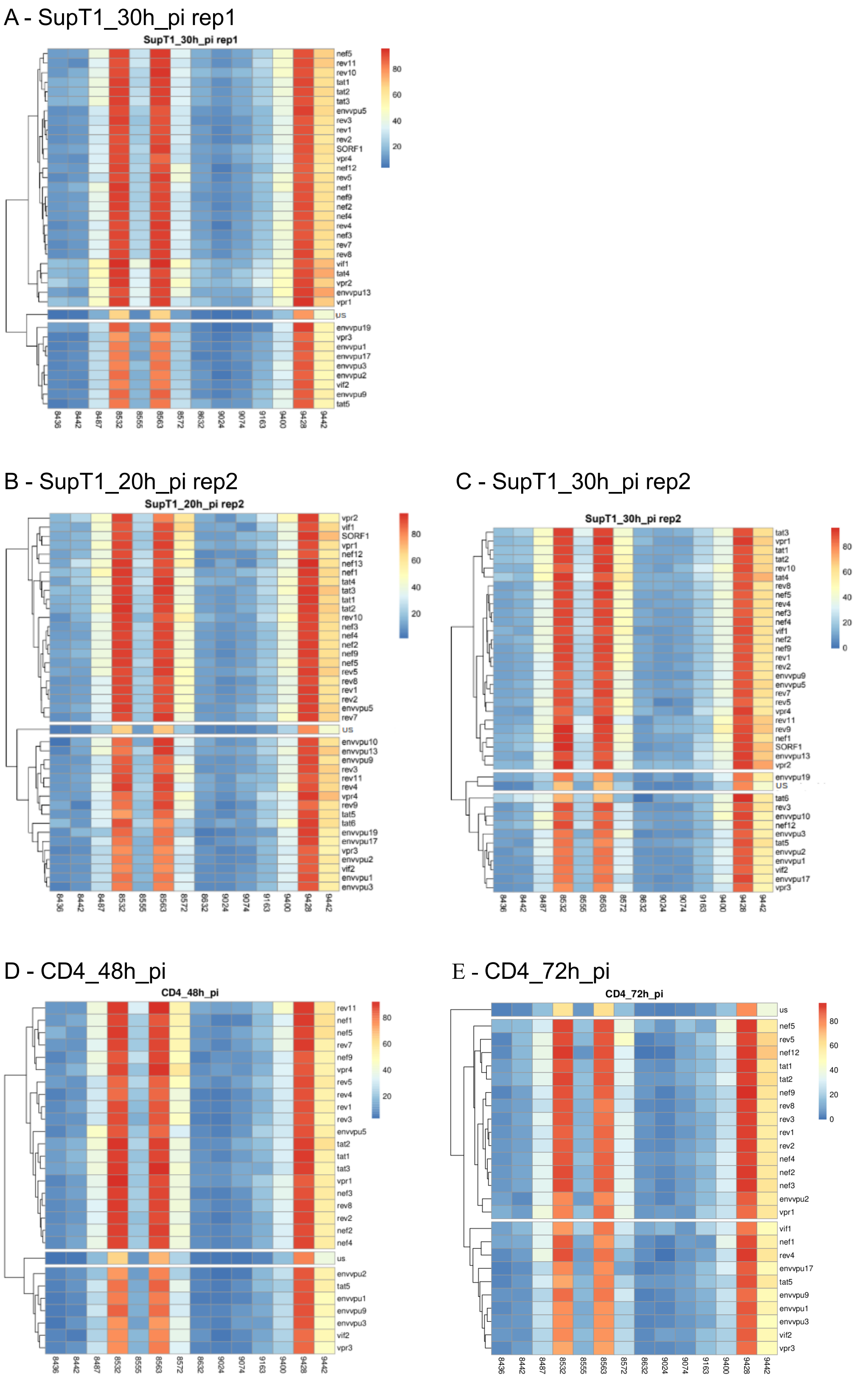

**Figure S4.** Heatmap of the methylation levels of the 14 m$^6$A sites detected in the various splicing isoforms. **(A-E)** Results for the samples indicated. Methylation level was calculated for each set of full-length reads corresponding to each isoform represented by ≥100 reads (Materials and Methods). Isoforms are clustered based on the methylation level of the 14 $m^6A$ sites and the clusters obtained correspond essentially to US, PS and CS isoforms.

A – SupT1_30h_pi

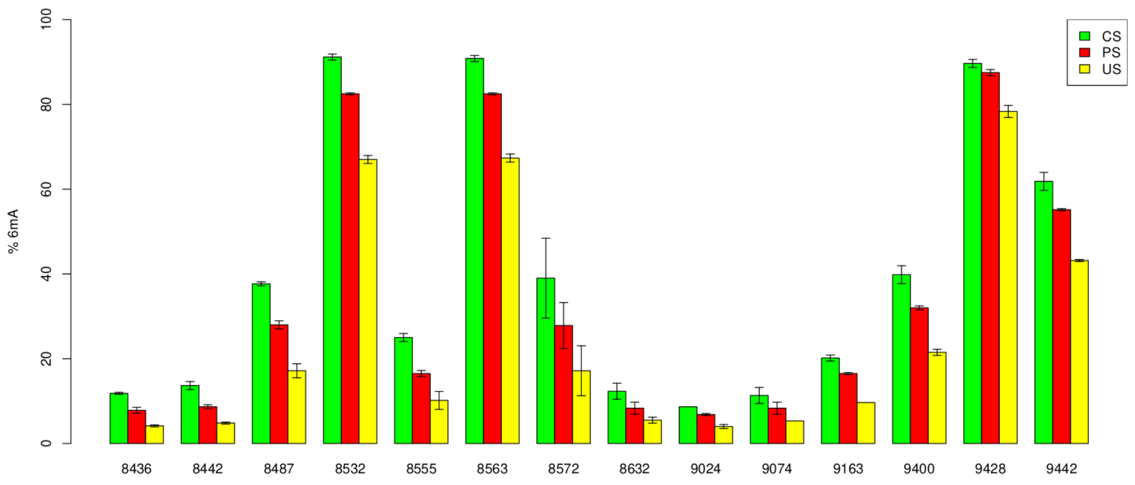

B - CD4_48h_pi

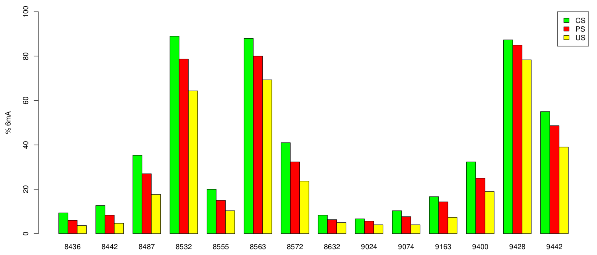

**C** - CD4_72h_pi

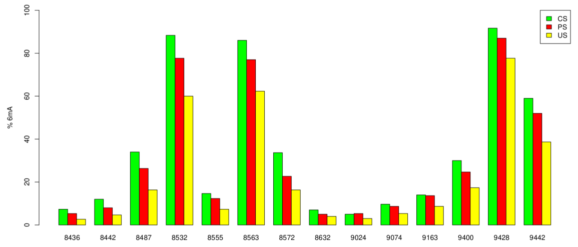

**Figure S5**. Methylation rates of m[6]A sites in CS, PS and US transcripts. **(A)** Methylation rates measured in SupT1_30h_pi rep1 and rep2 (mean ± SD). **(B,C)** Methylation rates measured in the indicated samples.

#### SupT1_30h_pi (rep1 and rep2)

A

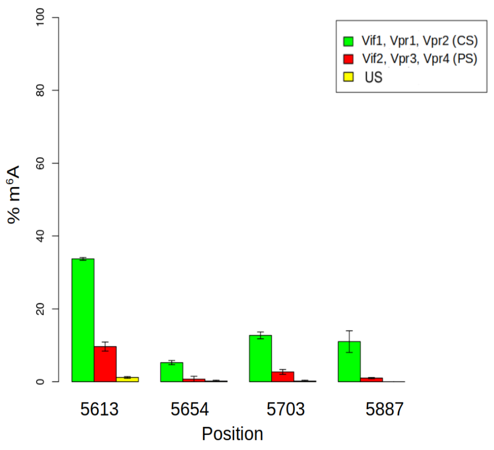

B

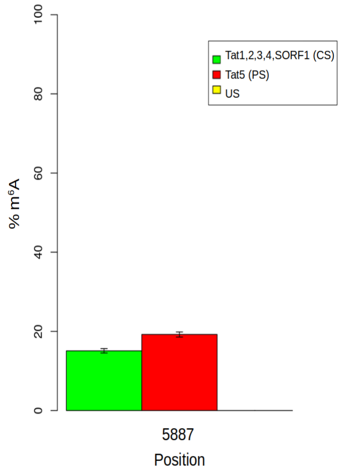

## CD4_48h_pi

C

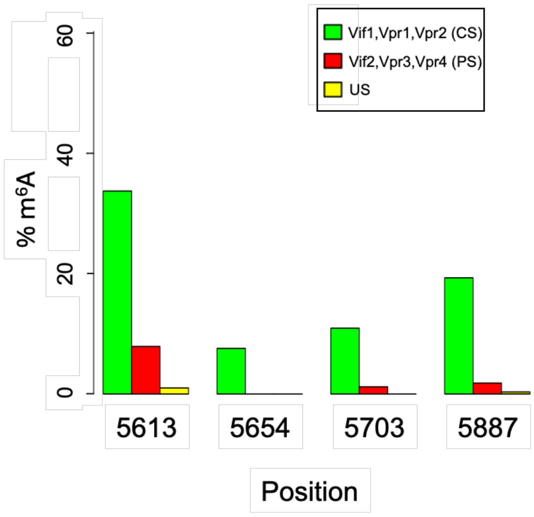

D

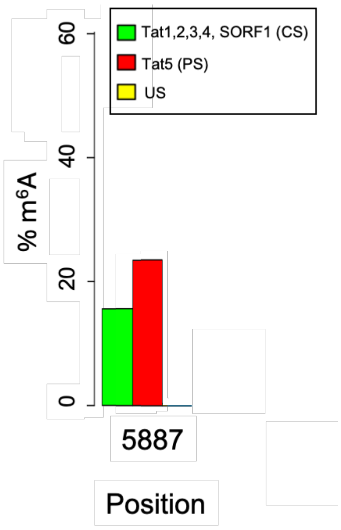

## CD4_72h_pi

E

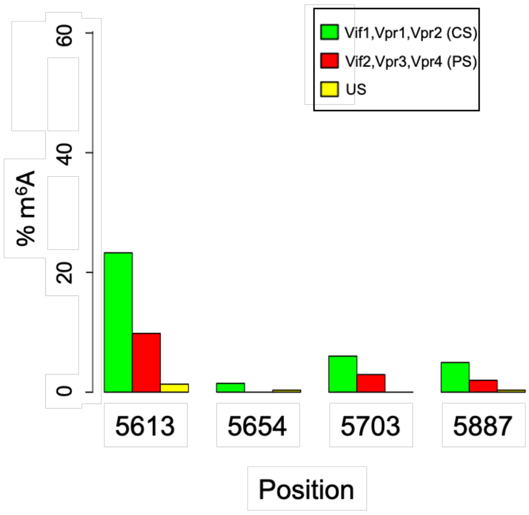

F

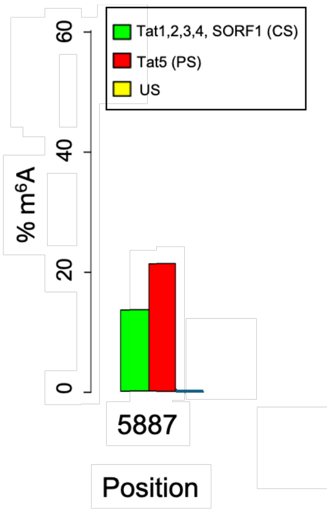

**Figure S6.** Methylation rates of the four isoform-specific m$^6$A sites in the samples indicated. The A5613, A5654 and A5703 sites are common to the Vif and Vpr isoforms; A5887 is common to the Vif, Vpr, Tat and SORF1 isoforms. **(A,C,E)** Methylation rates calculated for the four sites in CS (- intron 4, Vif1, Vpr1, Vpr2), PS (+ intron 4, Vif2, Vpr3, Vpr4) and US isoforms. **(B,D,F)** Methylation rates calculated for the A5887 site for the CS (Tat1, 2, 3, 4, SORF1), PS (Tat5) and US isoforms.

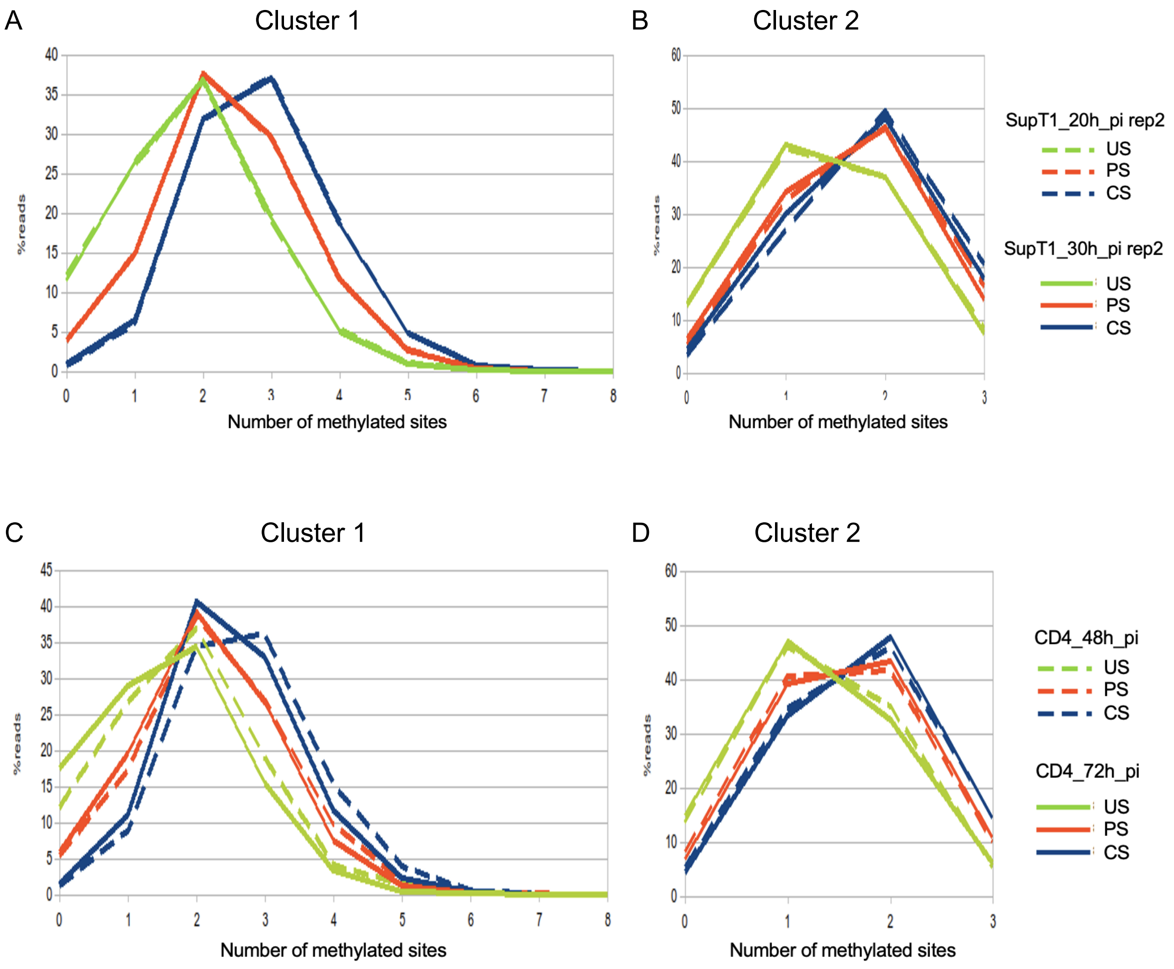

**Figure S7**. Single-molecule analysis of methylated m$^6$A sites. **(A)** Cluster 1 and **(B)** cluster 2 of the indicated SupT1 samples. **(C)** Cluster 1 and **(D)** cluster 2 of the indicated CD4 samples.

**SupT1_30h_pi rep1**

A (US)

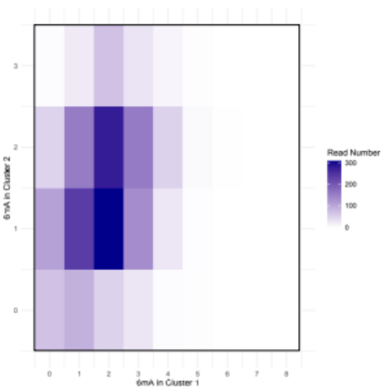

B (PS)

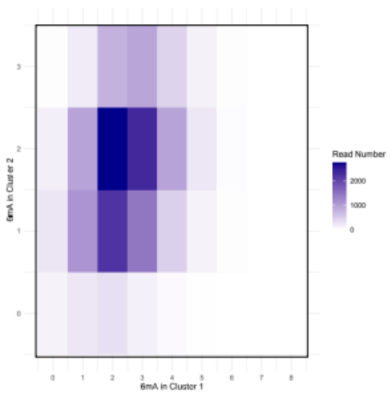

C (CS)

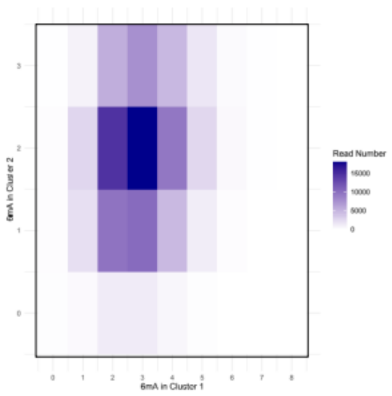

**SupT1_20h_pi rep2**

D (US)

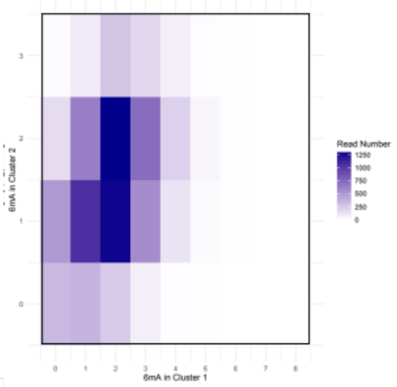

E (PS)

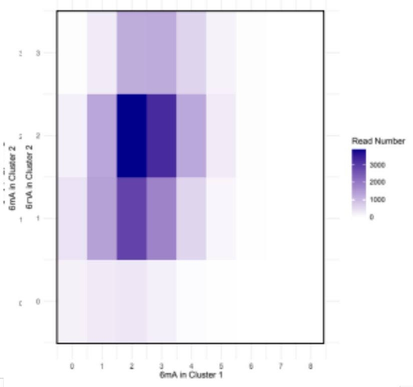

F (CS)

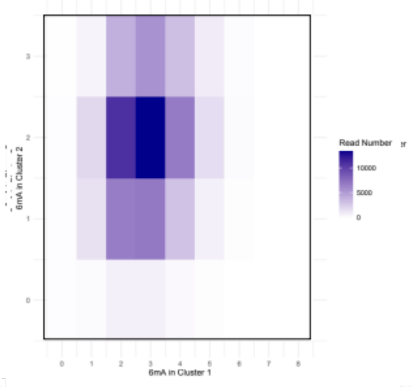

**SupT1_30h_pi rep2**

G (US)

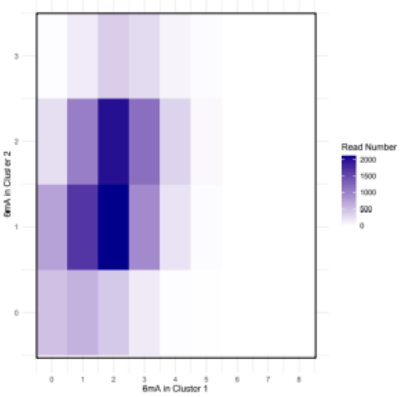

H (PS)

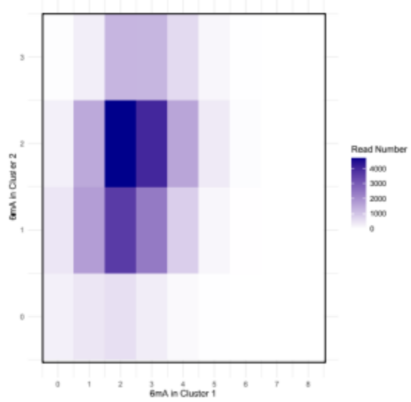

I (CS)

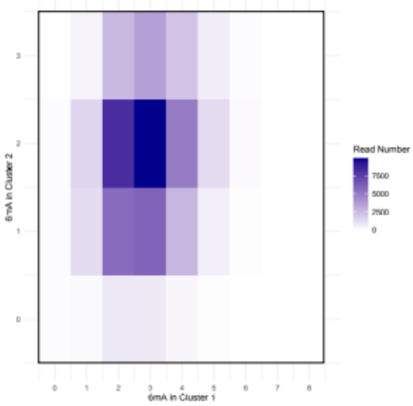

**CD4_48h_pi**

J (US) K (PS) L (CS)

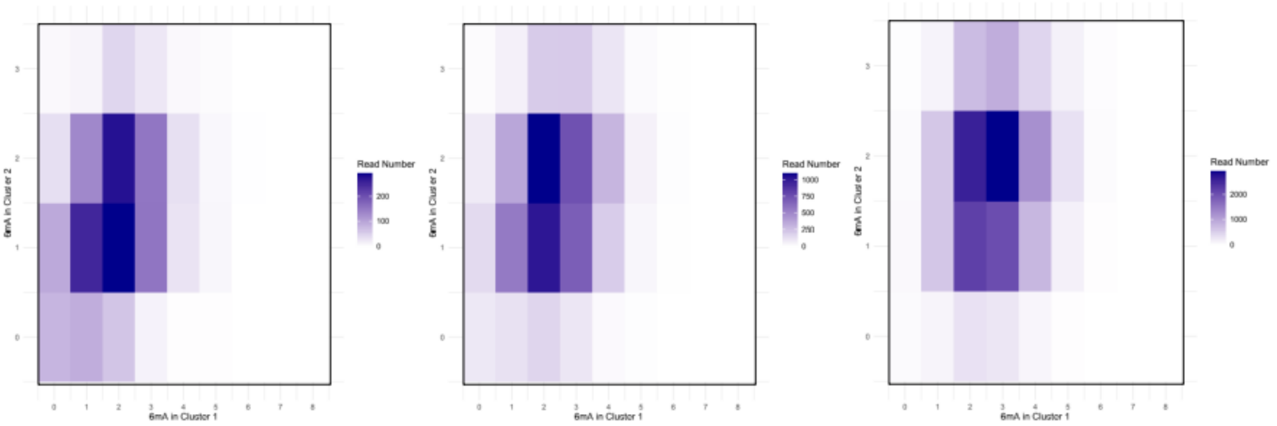

**CD4_72h_pi**

M (US) N (PS) O (CS)

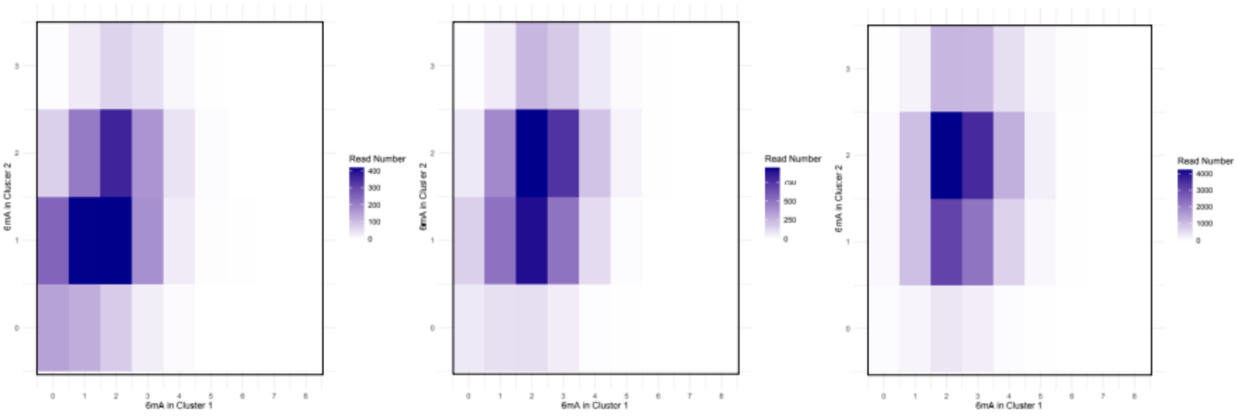

**Figure S8.** Single-molecule analysis of methylated m$^6$A sites illustrating the correlation between the methylation patterns for clusters 1 and 2 of the same transcripts. The intensity of the color on the heatmap indicates the number of reads, with the indicated numbers of methylated As in cluster 1 (0 to 8 m$^6$As) and cluster 2 (0 to 3 m$^6$As). US, unspliced; PS partially spliced; CS, completely spliced. One m$^6$A of one read is considered methylated if the probability of base modification is ≥ 0.5 (value in modkit extract), Materials and Methods).

### SUPPLEMENTARY TABLES

| | SupT1_ 20h_pi rep1 | SupT1_ 20h_pi rep2 | SupT1_ 30h_pi rep1 | SupT1_ 30h_pi rep2 | CD4_4 8h_pi | CD4_7 2h_pi | SupT1_ 20h_pi rep1 + ALKBH5 | SupT1_ 20h_pi rep2 + ALKBH5 | SupT1_ 20h_pi rep2 + FTO |
|---|---|---|---|---|---|---|---|---|---|
| Total reads ($10^6$) | 7.3 | 27.2 | 15.7 | 27.2 | 26.1 | 28.9 | 28.6 | 36.5 | 30.8 |
| HIV-1 reads | 386852 | 287479 | 1074370 | 345351 | 166733 | 393666 | 971749 | 313069 | 375951 |
| Infection rate (%) | 5.5 | 1.31 | 7.1 | 1.53 | 0.77 | 1.9 | 5.58 | 1.32 | 1.83 |
| Full-length reads | 80230 | 134086 | 164360 | 127964 | 33587 | 39278 | 4302 | 5825 | 10580 |
| Full length (%) | 20.74 | 46.64 | 15.3 | 37.05 | 20.14 | 9.98 | 0.44 | 1.86 | 2.81 |
| Mean length of mapped reads on VIH (kb) | 1.2 | 2.2 | 1.3 | 2.4 | 1.7 | 1.6 | 0.3 | 0.5 | 0.6 |

**Table S1.** Run statistics. Infection rate, number of full-length HIV reads divided by the total number of HIV and human reads, mean length of mapped reads on the HIV-1 genome (kb).

**A**

| Donor sites | Acceptor sites |
|---|---|
| D1: 743 | |
| D1c: 747 | |
| | A1: 4913 |
| | A1b: 4919 |
| D2: 4962 | |
| D2b: 5058 | |
| | A2: 5390 |
| D3 : 5463 | |
| | A3: 5777 |
| | A4c: 5936 |
| | A4a: 5954 |
| | A4b: 5960 |
| | A5: 5976 |
| | A5a: 5980 |
| | A5c: 5992 |
| D4: 6044 | |
| D4a: 6048 | |
| D4b: 6051 | |
| | A7: 8369 |

**B**

| | | SupT1_ 20h_pi rep1 | % | SupT1_ 20h_pi rep2 | % | SupT1_ 30h_pi rep1 | % | SupT1_ 30h_pi rep2 | % | CD4_ 48h_pi | % | CD4_ 72h_pi | % |
|---|---|---|---|---|---|---|---|---|---|---|---|---|---|
| US | | 610 | 0.83 | 10494 | 8.21 | 2387 | 1.57 | 16566 | 13.75 | 2286 | 7.12 | 3280 | 8.76 |
| envvpu1 | D1A5 | 3971 | 5.38 | 17206 | 13.47 | 12515 | 8.22 | 16904 | 14.04 | 3699 | 11.52 | 4054 | 10.83 |
| envvpu2 | D1A4b | 214 | 0.29 | 757 | 0.59 | 659 | 0.43 | 901 | 0.75 | 216 | 0.67 | 151 | 0.4 |
| envvpu3 | D1A4a | 139 | 0.19 | 462 | 0.36 | 370 | 0.24 | 451 | 0.37 | 269 | 0.84 | 348 | 0.93 |
| envvpu5 | D1A1D2A5 | 281 | 0.38 | 756 | 0.59 | 672 | 0.44 | 943 | 0.78 | 111 | 0.35 | 93 | 0.25 |
| envvpu9 | D1A2D3A5 | 619 | 0.84 | 1345 | 1.05 | 1304 | 0.86 | 2367 | 1.97 | 397 | 1.24 | 195 | 0.52 |
| envvpu10 | D1A2D3A4b | 46 | 0.06 | 61 | 0.05 | 54 | 0.04 | 143 | 0.12 | 30 | 0.09 | 9 | 0.02 |
| envvpu13 | D1A1D2A2D3A5 | 76 | 0.1 | 147 | 0.12 | 140 | 0.09 | 250 | 0.21 | 38 | 0.12 | 21 | 0.06 |
| envvpu17 | D1A5a | 92 | 0.12 | 488 | 0.38 | 350 | 0.23 | 466 | 0.39 | 91 | 0.28 | 100 | 0.27 |
| envvpu18 | D1cA4b | 2 | 0 | 6 | 0 | 3 | 0 | 9 | 0.01 | 3 | 0.01 | 2 | 0.01 |
| envvpu19 | D1cA5 | 49 | 0.07 | 127 | 0.1 | 120 | 0.08 | 140 | 0.12 | 30 | 0.09 | 40 | 0.11 |
| nef1 | D1A7 | 286 | 0.39 | 463 | 0.36 | 663 | 0.44 | 492 | 0.41 | 175 | 0.55 | 228 | 0.61 |
| nef2 | D1A5D4A7 | 29546 | 40.01 | 49676 | 38.88 | 65277 | 42.86 | 34533 | 28.67 | 8482 | 26.42 | 13658 | 36.47 |
| nef3 | D1A1D2A5D4A7 | 6934 | 9.39 | 7760 | 6.07 | 13894 | 9.12 | 7378 | 6.13 | 1066 | 3.32 | 1214 | 3.24 |

| | | | | | | | | | | | | | |
|---|---|---|---|---|---|---|---|---|---|---|---|---|---|
| nef4 | D1A2D3A5D4A7 | 7816 | 10.59 | 6156 | 4.82 | 11672 | 7.66 | 7242 | 6.01 | 1591 | 4.95 | 931 | 2.49 |
| nef5 | D1A1D2A2D3A5D4A7 | 2290 | 3.1 | 1512 | 1.18 | 3781 | 2.48 | 1897 | 1.58 | 376 | 1.17 | 162 | 0.43 |
| nef9 | D1A5aD4A7 | 690 | 0.93 | 1382 | 1.08 | 1540 | 1.01 | 857 | 0.71 | 205 | 0.64 | 357 | 0.95 |
| nef12 | D1cA5D4A7 | 214 | 0.29 | 331 | 0.26 | 496 | 0.33 | 268 | 0.22 | 65 | 0.2 | 105 | 0.28 |
| nef13 | D1A5D4aA7 | 25 | 0.03 | 35 | 0.03 | 58 | 0.04 | 27 | 0.02 | 8 | 0.02 | 10 | 0.03 |
| nef14 | D1A2D3A5D4aA7 | 8 | 0.01 | 8 | 0.01 | 11 | 0.01 | 8 | 0.01 | 2 | 0.01 | 0 | 0 |
| nef16 | D1cA5D4A7 | 10 | 0.01 | 10 | 0.01 | 23 | 0.02 | 6 | 0 | 1 | 0 | 3 | 0.01 |
| nef17 | D1A5D4bA7 | 8 | 0.01 | 9 | 0.01 | 22 | 0.01 | 9 | 0.01 | 2 | 0.01 | 2 | 0.01 |
| rev1 | D1A4bD4A7 | 3290 | 4.46 | 5743 | 4.5 | 7876 | 5.17 | 4360 | 3.62 | 1645 | 5.12 | 1350 | 3.61 |
| rev2 | D1A4aD4A7 | 3748 | 5.08 | 6522 | 5.11 | 8662 | 5.69 | 4546 | 3.77 | 3284 | 10.23 | 3979 | 10.63 |
| rev3 | D1A4cD4A7 | 213 | 0.29 | 348 | 0.27 | 539 | 0.35 | 252 | 0.21 | 396 | 1.23 | 584 | 1.56 |
| rev4 | D1A1D2A4bD4A7 | 363 | 0.49 | 504 | 0.39 | 830 | 0.55 | 445 | 0.37 | 169 | 0.53 | 110 | 0.29 |
| rev5 | D1A1D2A4aD4A7 | 199 | 0.27 | 326 | 0.26 | 464 | 0.3 | 218 | 0.18 | 137 | 0.43 | 111 | 0.3 |
| rev7 | D1A2D3A4bD4A7 | 786 | 1.06 | 641 | 0.5 | 1111 | 0.73 | 742 | 0.62 | 387 | 1.21 | 90 | 0.24 |
| rev8 | D1A2D3A4aD4A7 | 717 | 0.97 | 524 | 0.41 | 1006 | 0.66 | 626 | 0.52 | 507 | 1.58 | 190 | 0.51 |
| rev9 | D1A2D3A4cD4A7 | 33 | 0.04 | 38 | 0.03 | 52 | 0.03 | 35 | 0.03 | 51 | 0.16 | 34 | 0.09 |
| rev10 | D1A1D2A2D3A4bD4A7 | 191 | 0.26 | 154 | 0.12 | 315 | 0.21 | 169 | 0.14 | 81 | 0.25 | 23 | 0.06 |
| rev11 | D1A1D2A2D3A4aD4A7 | 202 | 0.27 | 161 | 0.13 | 335 | 0.22 | 151 | 0.13 | 130 | 0.4 | 42 | 0.11 |
| rev13 | D1cA4bD4A7 | 23 | 0.03 | 29 | 0.02 | 38 | 0.02 | 29 | 0.02 | 8 | 0.02 | 11 | 0.03 |
| SORF1 | D1A1D2bA3D4A7 | 218 | 0.3 | 145 | 0.11 | 358 | 0.24 | 155 | 0.13 | 66 | 0.21 | 84 | 0.22 |
| SORF3 | D1A1D2bA2D3A3D4A7 | 7 | 0.01 | 10 | 0.01 | 35 | 0.02 | 13 | 0.01 | 6 | 0.02 | 4 | 0.01 |
| tat1 | D1A3D4A7 | 4534 | 6.14 | 3746 | 2.93 | 4210 | 2.76 | 2101 | 1.74 | 1584 | 4.93 | 2379 | 6.35 |
| tat2 | D1A1D2A3D4A7 | 2354 | 3.19 | 2181 | 1.71 | 3641 | 2.39 | 1914 | 1.59 | 745 | 2.32 | 1019 | 2.72 |
| tat3 | D1A2D3A3D4A7 | 484 | 0.66 | 262 | 0.21 | 568 | 0.37 | 332 | 0.28 | 171 | 0.53 | 97 | 0.26 |
| tat4 | D1A1D2A2D3A3D4A7 | 171 | 0.23 | 109 | 0.09 | 294 | 0.19 | 120 | 0.1 | 73 | 0.23 | 53 | 0.14 |
| tat5 | D1A3 | 230 | 0.31 | 278 | 0.22 | 251 | 0.16 | 270 | 0.22 | 250 | 0.78 | 281 | 0.75 |
| tat6 | D1A1D2A3 | 16 | 0.02 | 31 | 0.02 | 39 | 0.03 | 32 | 0.03 | 34 | 0.11 | 39 | 0.1 |
| tat7 | D1A2D3A3 | 7 | 0.01 | 15 | 0.01 | 10 | 0.01 | 13 | 0.01 | 12 | 0.04 | 7 | 0.02 |
| tat9 | D1cA3D4A7 | 11 | 0.01 | 8 | 0.01 | 6 | 0 | 10 | 0.01 | 9 | 0.03 | 6 | 0.02 |
| vif1 | D1A1D4A7 | 98 | 0.13 | 267 | 0.21 | 389 | 0.26 | 280 | 0.23 | 39 | 0.12 | 112 | 0.3 |
| vif2 | D1A1 | 677 | 0.92 | 3199 | 2.5 | 2108 | 1.38 | 5234 | 4.35 | 547 | 1.7 | 591 | 1.58 |
| vif3 | D1cA1b | 2 | 0 | 4 | 0 | 10 | 0.01 | 7 | 0.01 | 2 | 0.01 | 3 | 0.01 |
| vif4 | D1cA1 | 0 | 0 | 11 | 0.01 | 9 | 0.01 | 22 | 0.02 | 2 | 0.01 | 2 | 0.01 |
| vif5 | D1A1b | 3 | 0 | 10 | 0.01 | 15 | 0.01 | 19 | 0.02 | 0 | 0 | 3 | 0.01 |
| vpr1 | D1A2D4A7 | 332 | 0.45 | 290 | 0.23 | 435 | 0.29 | 426 | 0.35 | 139 | 0.43 | 186 | 0.5 |
| vpr2 | D1A1D2A2D4A7 | 61 | 0.08 | 50 | 0.04 | 148 | 0.1 | 76 | 0.06 | 21 | 0.07 | 16 | 0.04 |

| | | | | | | | | | | | | | |
|---|---|---|---|---|---|---|---|---|---|---|---|---|---|
| vpr3 | D1A2 | 880 | 1.19 | 2793 | 2.19 | 2386 | 1.57 | 5651 | 4.69 | 2340 | 7.29 | 1022 | 2.73 |
| vpr4 | D1A1D2A2 | 64 | 0.09 | 166 | 0.13 | 142 | 0.09 | 335 | 0.28 | 131 | 0.41 | 54 | 0.14 |
| Total isoforms | | 73840 | | 127756 | | 152293 | | 120440 | | 32109 | | 37445 | |
| 2LTR* | | 1273 | 1.69 | 566 | 0.44 | 2391 | 1.55 | 878 | 0.72 | 52 | 0.16 | 109 | 0.29 |

**Table S2. (A)** List of the HIV-1 donor and acceptor splice sites. **(B)** Absolute and relative amounts of the HIV-1 transcript isoforms. %, amount calculated relative to all identified splicing isoforms (not including 2LTR); *, amount calculated relatively to all HIV-1 isoforms including 2LTR. Gray lines, the seven most abundant isoforms, accounting for about 80% of all isoforms.

| | SupT1_20h_pi rep2 | SupT1_30h_pi rep1 | SupT1_30h_pi rep2 | CD4_48h_pi | CD4_72h_pi |
|---|---|---|---|---|---|
| SupT1_20h_pi rep1 | 0.946 | 0.991 | 0.861 | 0.890 | 0.925 |
| SupT1_20h_pi rep2 | | 0.972 | 0.964 | 0.952 | 0.978 |
| SupT1_30h_pi rep1 | | | 0.891 | 0.911 | 0.946 |
| SupT1_30h_pi rep2 | | | | 0.941 | 0.933 |
| CD4_48h_pi | | | | | 0.971 |

**Table S3.** Pearson correlation coefficients for the relationship between the relative amounts of the splicing isoforms (Table S2B).

| m$^6$A position | Sequence | SupT1-20h-pi rep1 | SupT1-20h-pi rep2 | SupT1-30h-pi rep1 | SupT1-30h-pi rep2 | CD4-48h-pi | CD4-72h-pi |
|---|---|---|---|---|---|---|---|
| A8436 | AGACA | 9.7 | 6.3 | 8 | 5.7 | 4.7 | 3.7 |
| A8442 | AGACA | 10.3 | 7.0 | 9.7 | 6.7 | 6.3 | 5.3 |
| A8487 | GGACG | 30 | 22.7 | 27 | 22 | 19.7 | 18 |
| A8532 | AGACU | 79.3 | 72.7 | 79.3 | 72 | 68.3 | 63.7 |
| A8555 | GGAUU | 19.3 | 13 | 17.3 | 13 | 11.7 | 9.3 |
| A8563 | GAACU | 79.3 | 74 | 79.7 | 72.7 | 70.3 | 66.3 |
| A8572 | GGACG | 23.7 | 27 | 23.3 | 26 | 25.7 | 18.7 |
| A8632 | GAACU | 10.3 | 6.7 | 9.3 | 6.7 | 5.7 | 3.7 |
| A9024 | UGACU | 7.3 | 5.3 | 6.0 | 5.0 | 4.3 | 3.3 |
| A9074 | GGACU | 8.7 | 7.7 | 9.7 | 6.3 | 5.7 | 5.7 |
| A9163 | GAACU | 17.3 | 13 | 15 | 11.7 | 10.3 | 9.0 |
| A9400 | GAACU | 29.7 | 28.0 | 32.0 | 25.0 | 21.0 | 18.7 |
| A9428 | GGACU | 84.3 | 82.7 | 84.3 | 80.7 | 79.7 | 79.3 |
| A9442 | GGACU | 53.3 | 49 | 52.3 | 46 | 41 | 41.3 |

**Table S4.** Rates of methylation of the 14 m$^6$A sites detected in all HIV-1 transcripts in each of the 6 indicated samples (Materials and Methods). Underlined, 1-nt variants of the consensus DRACH sequences. Gray, the three most methylated sites, A8532, A8563, A9428.

| Oligos | Corresponding A positions | % modif | % modif ALKBH5 | % modif FTO |
|---|---|---|---|---|
| Oli1-mod | A8563 | 99.27 | 3.04 | 3.75 |
| Oli1-nmod | A8563 | 0.14 | 0.20 | 0.11 |
| Oli2-mod | A9428 | 99.55 | 3.24 | 30.81 |
| Oli2-nmod | A9428 | 1.50 | 1.13 | 0.53 |
| Oli2-mod | A9442 | 99.35 | 20.10 | 44.81 |
| Oli2-nmod | A9442 | 0.19 | 0.16 | 0.17 |
| All oligos | All other As * | ND | ND | ND |

Table S5. Validation of the methylated sites detected in synthetic RNA oligonucleotides. In $m^6A$-modified RNA oligonucleotides oli1-mod and oli2-mod methylated at corresponding positions of the HIV-1 reference genome (column 2), methylated sites were detected as described in Materials and Methods (column 3). For validation, the methylation rates were also measured after treatment with ALKBH5 and FTO demethylases (columns 4 and 5) showing a significant reduction of methylation rates at $m^6A$ sites. * indicates that at no modified read is detected (ND) at all other A positions, in modified and non-modified oligos (except for one position in oli2-nmod, A9407 detected with methylation rates <2.4%).

| Position | Sequence | SupT1_20h_pi rep1 | SupT1_20h_pi+ ALKBH5 rep1 | | SupT1_20h_pi rep2 | SupT1_20h_pi+ ALKBH5 rep2 | | SupT1_20h_pi rep2+FTO | |
|---|---|---|---|---|---|---|---|---|---|
| | | % methyl. | % methyl. | p-value | % methyl. | % methyl. | *p*-value | % methyl. | *p*-value |
| A8436 | AGACA | 9.7 | 1 | 2.00E+101 | 6.3 | 2 | 9.60E-38 | 3.3 | 4.41e-18 |
| A8442 | AGACA | 10.3 | 1.7 | 7.90E-147 | 7 | 2.7 | 2.40E-36 | 5.7 | 1.5e-4 |
| A8487 | GGACG | 30 | 7.3 | 1.60E-265 | 22.7 | 11.7 | 3.90E-75 | NS | NS |
| A8532 | AGACU | 79.3 | 49.3 | 0 | 72.7 | 51.7 | 6.70E-191 | 35.7 | 0 |
| A8555 | GGAUU | 19.3 | 11.3 | 4.00E-56 | 13 | 9 | 2.50E-14 | NS | NS |
| A8563 | GAACU | 79.3 | 60 | 5.30E-174 | 74 | 60.7 | 8.60E-82 | 16.7 | 0 |
| A8572 | GGACG | 23.7 | 12.3 | 1.80E-24 | 27 | 17.3 | 1.70E-15 | 3.0 | 1.25e-23 |
| A8632 | GAACU | 10.3 | 3 | 1.10E-80 | 6.7 | 2.3 | 3.30E-36 | 1 | 7.02e-76 |
| A9024 | UGACU | 7.3 | 1.3 | 3.00E-87 | 5.3 | 1.3 | 1.00E-43 | 1 | 8.33e-55 |
| A9074 | GGACU | 8.7 | 5 | 1.10E-22 | 7.7 | 5.3 | 3.20E-09 | 2.7 | 2.91e-304 |
| A9163 | GAACU | 17.3 | 3.3 | 1.00E-165 | 13 | 5.3 | 4.30E-58 | 1.7 | 9.6e-140 |
| A9400 | GAACU | 29.7 | 18.3 | 3.20E-93 | 28 | 21.7 | 6.90E-24 | 8 | 8.43e-241 |
| A9428 | GGACU | 84.3 | 22.3 | 0 | 82.7 | 25.3 | 0 | 37.7 | 0 |
| A9442 | GGACU | 53.3 | 18.3 | 0 | 49 | 22.3 | 1.5e-311 | 25.3 | 4.90e-232 |

**Table S6.** Demethylation efficiency for the 14 $m^6A$ sites detected in all transcripts of SupT1_20h_pi rep1 and SupT1_20h_pi rep2. SupT1_20h_pi+ALKBH5 corresponds to the SupT1_20h_pi sample demethylated with ALKBH5, and SupT1_20h_pi rep2+FTO to the SupT1_20h_pi rep2 sample demethylated with FTO (Materials and Methods). *P*-values correspond to the difference between the numbers of methylated reads in SupT1_20h_pi and in SupT1_20h_pi+demethylase (ALKBH5 or FTO) (Materials and Methods); NS, not significant.

| SupT1_20h _rep2 | | | | | | | | | | | |
|---|---|---|---|---|---|---|---|---|---|---|---|
| Position | Sequence | **nef2** | +ALKBH5 | | +FTO | | **nef3** | +ALKBH5 | | +FTO | |
| | | % methyl. | % methyl. | p-value | % methyl. | p-value | % methyl. | % methyl. | p-value | % methyl. | p-value |
| A8436 | AGACA | 11 | 5,67 | 5,18E-14 | 4,33 | 5,90E-34 | 9,67 | 5,67 | 2,90E-02 | 3,66 | 1,49E-06 |
| A8442 | AGACA | 13,67 | 8,33 | 1,67E-12 | 6,66 | 1,17E-30 | 12,33 | 10 | 0,21 | 9 | 1,73E-02 |
| A8487 | GGACG | 38,33 | 9,67 | 1,78E-163 | NS | NS | 39 | 14,67 | 5,63E-20 | NS | NS |
| A8532 | AGACU | 92 | 74 | 4,01E-140 | 43,33 | 0 | 92 | 61,67 | 3,90E-87 | 49,33 | 7,33E-231 |
| A8555 | GGAUU | 21 | 14 | 4,39E-15 | NS | NS | 23 | 14,33 | 2,27E-04 | NS | NS |
| A8563 | GAACU | 90,67 | 71,3 | 1,43E-140 | 20 | 0 | 91,33 | 74 | 1,91E-28 | 19 | 0 |
| A8572 | GGACG | 45,67 | 24,33 | 7,00E-20 | 4,07 | 1,06E-26 | 43,67 | 29,4 | 6,70E-03 | NS | NS |
| A8632 | GAACU | 11 | 4 | 1,79E-28 | 1,33 | 8,61E-85 | 11,33 | 3,67 | 9,32E-06 | 1,33 | 2,65E-14 |
| A9024 | UGACU | 7,67 | 2 | 1,23E-23 | 1 | 1,83E-53 | 8,67 | 2,33 | 2,50E-05 | 1 | 8,62E-11 |
| A9074 | GGACU | 11 | 6 | 4,96E-13 | 3 | 1,72E-51 | 11,67 | 7 | 7,80E-03 | 2,33 | 7,61E-12 |
| A9163 | GAACU | 18,33 | 7 | 1,41E-39 | 2,66 | 4,15E-111 | 19 | 9,33 | 2,28E-05 | 2 | 1,19E-20 |
| A9400 | GAACU | 39,67 | 26,67 | 1,43E-33 | 10,33 | 7,22E-227 | 42 | 31,67 | 1,03E-04 | 10 | 1,37E-44 |
| A9428 | GGACU | 91,67 | 26,67 | 0 | 39,66 | 0 | 90,67 | 24 | 0 | 37 | 1,19E-281 |
| A9442 | GGACU | 61,67 | 26,33 | 4,44E-214 | 28,33 | 1,11E-255 | 62,67 | 23,67 | 2,56E-48 | 30,66 | 1,01E-49 |

| Position | Sequence | **nef4** | +ALKBH5 | | +FTO | | **rev1** | +ALKBH5 | | +FTO | |
|---|---|---|---|---|---|---|---|---|---|---|---|
| | | % methyl. | % methyl. | p-value | % methyl. | p-value | % methyl. | % methyl. | p-value | % methyl. | p-value |
| A8436 | AGACA | 10 | 3,78 | 2,80E-03 | 5,66 | 2,97E-03 | 9,67 | 7,94 | 0,4 | 5,66 | 3,62E-03 |
| A8442 | AGACA | 11,33 | 9,25 | 0,39 | 8,66 | 7,93E-02 | 11,33 | 6,55 | 1,80E-02 | 11,66 | 8,68E-01 |
| A8487 | GGACG | 37,33 | 17,39 | 1,55E-10 | NS | NS | 34,67 | 13,13 | 3,11E-13 | NS | NS |
| A8532 | AGACU | 91,33 | 33,5 | 2,61E-192 | 40 | 2,52E-257 | 91 | 15,63 | 0 | 40 | 3,53E-241 |
| A8555 | GGAUU | 22,67 | 9,78 | 1,49E-06 | ND | ND | 20,67 | 13,59 | 6,37E-03 | NS | NS |
| A8563 | GAACU | 91 | 72,9 | 7,83E-22 | 17,66 | 0 | 89,33 | 64,54 | 5,66E-37 | 19,66 | 1,20E-259 |
| A8572 | GGACG | 42,67 | 27,3 | 1,10E-02 | NS | NS | 45,67 | 22,75 | 6,22E-05 | NS | NS |
| A8632 | GAACU | 10,67 | 2,27 | 1,20E-05 | 1,33 | 5,54E-12 | 10,33 | 3,14 | 1,20E-04 | 1,33 | 1,14E-10 |
| A9024 | UGACU | 8 | 1,95 | 6,30E-04 | 1 | 7,15E-09 | 7,67 | 0,7 | 2,87E-05 | 0,66 | 5,27E-09 |
| A9074 | GGACU | 10,67 | 7,15 | 9,80E-02 | 4 | 2,51E-06 | 11 | 4,98 | 2,20E-03 | 3,33 | 6,01E-08 |
| A9163 | GAACU | 17,67 | 6,28 | 7,85E-06 | 1,33 | 1,73E-18 | 19,67 | 5,13 | 1,74E-08 | 2,33 | 3,11E-17 |
| A9400 | GAACU | 40,67 | 28,79 | 1,86E-04 | 10,66 | 4,45E-36 | 39,67 | 28,76 | 3,85E-04 | 9,66 | 8,61E-35 |
| A9428 | GGACU | 90,67 | 21,36 | 1,99E-244 | 39 | 1,26E-223 | 92,33 | 20,06 | 0 | 39,66 | 3,40E-252 |
| A9442 | GGACU | 62 | 19,9 | 8,37E-41 | 30,66 | 9,49E-42 | 61,67 | 16,33 | 5,17E-50 | 33,66 | 5,03E-31 |

| Position | Sequence | **rev2** | +ALKBH5 | | +FTO | | **tat1** | +ALKBH5 | | +FTO | |
|---|---|---|---|---|---|---|---|---|---|---|---|
| | | % methyl. | % methyl. | p-value | % methyl. | p-value | % methyl. | % methyl. | p-value | % methyl. | p-value |
| A8436 | AGACA | 9 | 7,33 | 0,39 | 6,66 | 8,40E-02 | 14,33 | 12,26 | 0,61 | 13,22 | 7,62E-01 |
| A8442 | AGACA | 11,67 | 9,33 | 0,22 | 9,66 | 1,35E-01 | 17 | 12,81 | 0,24 | 15,66 | 6,41E-01 |
| A8487 | GGACG | 33,33 | 13,67 | 1,19E-13 | NS | NS | 40,67 | 20,74 | 4,17E-06 | NS | NS |
| A8532 | AGACU | 91 | 24 | 9,71E-307 | 39,66 | 1,03E-271 | 93 | 33,47 | 5,66E-136 | 39,39 | 7,10E-148 |
| A8555 | GGAUU | 20,33 | 8,67 | 4,82E-07 | NS | NS | 21,33 | 14 | 5,49E-02 | ND | ND |
| A8563 | GAACU | 90 | 68,67 | 8,46E-34 | 20 | 0 | 91,33 | 67,58 | 1,43E-20 | 23,24 | 2,11E-142 |
| A8572 | GGACG | 46 | 23,1 | 6,36E-06 | NS | NS | 42,33 | NS | NS | NS | NS |

| | | | | | | | | | | | |
|---|---|---|---|---|---|---|---|---|---|---|---|
| A8632 | GAACU | 11 | 3,67 | 5,92E-05 | 0,33 | 1,43E-15 | 11,67 | 1,82 | 6,10E-04 | 0,33 | 4,04E-07 |
| A9024 | UGACU | 8 | 1 | 1,08E-05 | 1,33 | 1,55E-08 | 9 | 1,13 | 2,30E-03 | 0,11 | 6,93E-06 |
| A9074 | GGACU | 11 | 7 | 3,00E-02 | 4 | 1,59E-07 | 12,67 | 12,64 | 1 | 3,57 | 2,20E-04 |
| A9163 | GAACU | 18,67 | 9,33 | 5,77E-05 | 2,33 | 5,28E-18 | 22 | 12,01 | 1,40E-02 | 4,59 | 8,68E-08 |
| A9400 | GAACU | 38 | 28,67 | 7,30E-04 | 10,66 | 3,47E-33 | 40,67 | 35,3 | 2,40E-01 | 11,16 | 1,87E-15 |
| A9428 | GGACU | 91,67 | 24,67 | 0 | 39 | 6,34E-268 | 92,67 | 27,34 | 3,38E-156 | 45,08 | 1,02E-110 |
| A9442 | GGACU | 61,67 | 19,33 | 5,04E-50 | 28,33 | 8,83E-48 | 65 | 17,71 | 3,06E-28 | 32,13 | 8,60E-20 |

Table S7 : Detection of $m^6A$ on the SupT1_20h_pi rep2 sample without demethylase treatment and after treatment with demethylase (ALKBH5 and FTO). NS corresponds to a total read count < 100 (non-significant); ND indicates that at no modified read is detected.

**A - SupT1_20h_pi rep1**

| | 8436 | 8442 | 8487 | 8532 | 8555 | 8563 | 8572 | 8632 | 9024 | 9074 | 9163 | 9400 | 9428 | 9442 |
|---|---|---|---|---|---|---|---|---|---|---|---|---|---|---|
| envvpu1 | 9.00 | 11.00 | 31.00 | 85.33 | 18.00 | 84.67 | 24.33 | 8.67 | 7.67 | 9.00 | 18.00 | 32.33 | 88.67 | 59.00 |
| envvpu2 | 8.12 | 13.37 | 33.63 | 86.14 | 18.83 | 84.77 | 35.48 | 12.00 | 10.58 | 9.33 | 21.21 | 31.54 | 87.42 | 59.77 |
| envvpu3 | 11.45 | 14.48 | 37.81 | 89.22 | 18.87 | 88.22 | 22.58 | 11.31 | 10.83 | 12.97 | 18.33 | 33.51 | 91.33 | 52.72 |
| envvpu5 | 14.94 | 11.89 | 43.11 | 93.81 | 24.92 | 92.54 | 28.20 | 12.85 | 7.77 | 10.32 | 26.73 | 42.67 | 90.46 | 65.72 |
| envvpu9 | 14.33 | 12.00 | 36.67 | 89.00 | 26.67 | 88.33 | 30.82 | 11.33 | 9.00 | 9.67 | 27.67 | 33.67 | 89.67 | 62.67 |
| nef1 | 14.18 | 12.26 | 39.08 | 92.82 | 28.73 | 92.46 | 36.00 | 13.40 | 10.44 | 9.64 | 22.14 | 36.62 | 90.65 | 68.79 |
| nef12 | 13.61 | 15.46 | 42.78 | 88.73 | 27.75 | 93.39 | 41.97 | 11.77 | 7.23 | 12.10 | 26.31 | 38.24 | 89.33 | 61.10 |
| nef2 | 14.00 | 14.00 | 41.33 | 92.67 | 27.33 | 92.33 | 33.67 | 14.33 | 9.33 | 11.00 | 22.00 | 38.00 | 91.00 | 63.00 |
| nef3 | 14.33 | 16.00 | 44.67 | 93.67 | 30.33 | 93.00 | 36.00 | 14.33 | 11.00 | 11.33 | 25.67 | 41.33 | 90.33 | 66.33 |
| nef4 | 16.00 | 15.67 | 45.67 | 93.67 | 31.67 | 93.33 | 37.33 | 16.33 | 12.67 | 10.67 | 27.67 | 40.00 | 91.00 | 66.33 |
| nef5 | 18.33 | 17.00 | 49.67 | 94.67 | 35.67 | 94.00 | 42.33 | 19.33 | 14.67 | 11.67 | 29.00 | 42.00 | 92.00 | 69.33 |
| nef9 | 13.00 | 15.00 | 41.67 | 92.00 | 25.00 | 92.67 | 37.44 | 13.67 | 11.33 | 11.33 | 23.00 | 36.33 | 90.33 | 66.33 |
| rev1 | 12.67 | 14.00 | 37.00 | 92.00 | 24.33 | 90.67 | 32.00 | 13.00 | 9.33 | 9.00 | 22.67 | 37.67 | 91.00 | 62.00 |
| rev10 | 10.75 | 18.48 | 41.39 | 92.68 | 31.41 | 90.54 | 33.72 | 15.48 | 13.17 | 16.88 | 27.25 | 45.02 | 91.37 | 65.40 |
| rev11 | 15.10 | 15.91 | 41.05 | 95.12 | 31.18 | 94.15 | 52.88 | 14.10 | 16.66 | 15.40 | 26.08 | 44.85 | 91.55 | 71.86 |
| rev2 | 12.67 | 12.67 | 37.33 | 90.33 | 24.33 | 90.67 | 31.67 | 12.00 | 9.67 | 10.67 | 21.33 | 36.67 | 90.00 | 60.67 |
| rev3 | 10.95 | 16.66 | 30.68 | 88.50 | 24.78 | 90.60 | 33.52 | 8.55 | 6.79 | 11.39 | 19.84 | 35.14 | 88.76 | 61.13 |
| rev4 | 13.67 | 12.33 | 45.00 | 90.67 | 30.00 | 94.00 | 29.37 | 12.67 | 13.67 | 12.67 | 25.33 | 39.00 | 90.00 | 60.67 |
| rev5 | 11.64 | 13.28 | 33.21 | 94.26 | 28.68 | 93.90 | 33.63 | 16.46 | 9.68 | 9.06 | 23.33 | 37.76 | 90.90 | 61.83 |
| rev7 | 14.67 | 16.00 | 41.00 | 93.00 | 29.67 | 93.33 | 42.52 | 18.00 | 13.00 | 12.33 | 30.33 | 38.33 | 91.00 | 65.00 |
| rev8 | 18.33 | 17.33 | 42.33 | 93.67 | 32.67 | 93.67 | 34.26 | 13.33 | 13.00 | 11.00 | 28.67 | 39.67 | 92.00 | 64.67 |
| SORF1 | 15.61 | 17.37 | 45.83 | 95.40 | 27.48 | 92.54 | 21.55 | 17.93 | 13.46 | 14.60 | 31.49 | 42.18 | 89.19 | 65.47 |
| tat1 | 13.33 | 13.33 | 38.67 | 90.67 | 22.00 | 91.33 | 34.00 | 13.67 | 10.00 | 12.33 | 23.67 | 31.33 | 91.00 | 62.33 |
| tat2 | 18.00 | 18.33 | 47.67 | 93.67 | 32.33 | 93.33 | 35.67 | 16.00 | 13.33 | 11.33 | 28.67 | 39.67 | 91.00 | 67.67 |
| tat3 | 19.00 | 17.67 | 49.00 | 94.33 | 32.33 | 93.33 | 43.70 | 18.00 | 12.67 | 11.00 | 27.67 | 41.33 | 92.00 | 68.67 |
| tat4 | 25.31 | 16.43 | 51.76 | 91.88 | 39.37 | 96.89 | 51.67 | 21.15 | 23.18 | 12.30 | 34.00 | 40.95 | 93.42 | 69.13 |
| tat5 | 10.23 | 10.98 | 27.91 | 82.54 | 22.81 | 85.05 | 21.41 | 9.95 | 9.60 | 13.55 | 26.76 | 26.27 | 89.54 | 61.77 |
| us | 4.33 | 5.67 | 19.00 | 68.67 | 10.33 | 70.00 | 12.99 | 5.67 | 5.33 | 5.00 | 11.33 | 17.33 | 81.33 | 44.33 |
| vif2 | 11.33 | 10.33 | 30.67 | 83.67 | 18.00 | 82.00 | 21.81 | 10.33 | 10.00 | 8.00 | 18.00 | 33.00 | 85.33 | 59.67 |
| vpr1 | 12.00 | 20.67 | 47.00 | 92.33 | 30.00 | 92.67 | 37.22 | 18.33 | 14.00 | 16.33 | 30.73 | 35.13 | 91.29 | 67.36 |
| vpr3 | 9.33 | 8.33 | 29.33 | 80.33 | 16.33 | 80.00 | 19.42 | 10.33 | 6.33 | 9.00 | 17.33 | 27.00 | 88.67 | 53.00 |

**B - SupT1_30h_pi rep1**

| | 8436 | 8442 | 8487 | 8532 | 8555 | 8563 | 8572 | 8632 | 9024 | 9074 | 9163 | 9400 | 9428 | 9442 |
|---|---|---|---|---|---|---|---|---|---|---|---|---|---|---|
| envvpu1 | 7.00 | 8.67 | 27.00 | 82.33 | 16.00 | 82.67 | 23.00 | 8.33 | 6.67 | 9.33 | 15.00 | 32.00 | 88.00 | 55.67 |
| envvpu13 | 13.71 | 10.78 | 44.99 | 89.45 | 36.43 | 91.19 | 41.19 | 15.18 | 14.74 | 15.49 | 25.68 | 49.71 | 92.21 | 75.56 |
| envvpu17 | 7.33 | 10.00 | 27.67 | 81.67 | 13.33 | 82.00 | 23.12 | 7.00 | 7.00 | 10.67 | 15.00 | 30.67 | 86.67 | 54.33 |
| envvpu19 | 12.84 | 12.06 | 29.28 | 87.28 | 18.09 | 86.83 | 19.76 | 8.01 | 3.64 | 5.53 | 8.43 | 33.11 | 91.81 | 53.41 |
| envvpu2 | 8.33 | 8.67 | 28.00 | 84.33 | 15.67 | 81.00 | 23.06 | 11.33 | 5.33 | 8.33 | 17.00 | 34.67 | 86.67 | 51.33 |

| | | | | | | | | | | | | | | |
|---|---|---|---|---|---|---|---|---|---|---|---|---|---|---|
| envvpu3 | 7.33 | 9.67 | 28.33 | 81.00 | 19.67 | 84.00 | 22.68 | 9.33 | 7.00 | 8.67 | 17.33 | 36.67 | 88.00 | 54.00 |
| envvpu5 | 9.67 | 10.33 | 32.00 | 90.33 | 23.00 | 88.67 | 31.31 | 11.67 | 9.67 | 14.67 | 19.00 | 40.00 | 91.67 | 65.67 |
| envvpu9 | 8.67 | 9.33 | 31.00 | 86.00 | 18.00 | 86.33 | 27.33 | 10.00 | 7.00 | 10.67 | 20.00 | 38.00 | 87.67 | 56.67 |
| nef1 | 12.00 | 17.00 | 40.33 | 93.00 | 24.33 | 90.67 | 35.21 | 12.00 | 9.67 | 12.00 | 19.67 | 45.00 | 89.00 | 62.33 |
| nef12 | 12.67 | 11.67 | 35.67 | 91.00 | 25.00 | 89.00 | 42.24 | 13.33 | 6.33 | 11.00 | 19.00 | 39.00 | 87.67 | 59.00 |
| nef2 | 11.00 | 13.33 | 37.00 | 92.00 | 24.67 | 91.00 | 31.67 | 12.67 | 7.67 | 11.67 | 19.00 | 39.67 | 90.00 | 62.00 |
| nef3 | 11.00 | 12.67 | 37.00 | 91.00 | 26.33 | 92.33 | 34.00 | 13.67 | 9.00 | 12.67 | 21.33 | 43.00 | 88.67 | 64.33 |
| nef4 | 11.00 | 12.00 | 36.67 | 91.33 | 25.00 | 91.33 | 31.33 | 13.33 | 9.33 | 12.33 | 21.00 | 41.00 | 89.67 | 63.33 |
| nef5 | 12.00 | 13.00 | 40.33 | 91.33 | 29.33 | 92.67 | 35.00 | 16.33 | 10.67 | 14.33 | 24.33 | 45.67 | 89.00 | 67.67 |
| nef9 | 11.67 | 12.33 | 38.00 | 91.67 | 23.33 | 91.33 | 30.67 | 10.67 | 8.33 | 11.00 | 20.00 | 38.33 | 89.67 | 63.33 |
| rev1 | 10.00 | 11.67 | 32.67 | 90.00 | 22.00 | 89.67 | 31.33 | 12.33 | 7.67 | 12.00 | 19.00 | 38.67 | 89.67 | 61.00 |
| rev10 | 11.60 | 16.33 | 37.67 | 90.33 | 26.00 | 93.67 | 35.32 | 18.87 | 14.94 | 18.45 | 25.64 | 46.29 | 89.19 | 63.75 |
| rev11 | 11.00 | 8.67 | 35.00 | 89.67 | 30.33 | 92.33 | 31.67 | 15.00 | 10.00 | 13.67 | 27.63 | 45.33 | 90.67 | 68.48 |
| rev2 | 10.00 | 11.00 | 32.00 | 89.67 | 22.33 | 90.00 | 32.67 | 11.67 | 7.67 | 10.67 | 19.00 | 39.33 | 89.33 | 60.33 |
| rev3 | 10.33 | 12.33 | 29.67 | 91.67 | 21.00 | 89.67 | 31.43 | 10.00 | 8.67 | 11.33 | 18.33 | 37.00 | 88.67 | 63.00 |
| rev4 | 8.67 | 11.67 | 33.33 | 90.67 | 25.00 | 93.00 | 33.20 | 12.33 | 6.33 | 13.33 | 20.33 | 44.00 | 88.33 | 65.33 |
| rev5 | 9.00 | 11.00 | 29.67 | 90.33 | 23.67 | 91.67 | 42.43 | 13.33 | 8.00 | 12.67 | 22.33 | 43.67 | 87.33 | 62.67 |
| rev7 | 9.00 | 11.33 | 35.67 | 90.00 | 26.33 | 91.67 | 32.33 | 15.67 | 9.67 | 13.67 | 21.67 | 40.33 | 90.00 | 62.33 |
| rev8 | 10.00 | 11.67 | 35.00 | 91.00 | 24.67 | 91.67 | 32.08 | 14.00 | 9.33 | 14.67 | 22.67 | 42.67 | 91.00 | 63.33 |
| SORF1 | 12.00 | 14.00 | 33.33 | 90.33 | 26.33 | 92.00 | 29.76 | 13.67 | 11.00 | 16.67 | 23.00 | 37.33 | 89.33 | 64.00 |
| tat1 | 15.67 | 17.67 | 41.00 | 91.67 | 25.33 | 91.67 | 32.67 | 15.00 | 9.67 | 14.67 | 23.00 | 40.33 | 91.33 | 63.67 |
| tat2 | 15.00 | 17.67 | 40.67 | 91.67 | 28.33 | 92.33 | 33.67 | 16.00 | 10.67 | 15.00 | 23.33 | 43.00 | 90.00 | 65.67 |
| tat3 | 16.67 | 18.00 | 42.67 | 91.67 | 30.00 | 91.67 | 34.77 | 15.67 | 11.00 | 15.33 | 24.33 | 40.00 | 89.67 | 63.67 |
| tat4 | 18.01 | 17.09 | 50.58 | 93.05 | 33.07 | 90.31 | 39.89 | 20.77 | 15.18 | 22.33 | 31.67 | 47.94 | 90.13 | 74.81 |
| tat5 | 5.98 | 9.13 | 30.40 | 83.34 | 17.25 | 83.01 | 29.00 | 17.10 | 10.15 | 11.25 | 20.41 | 37.17 | 87.73 | 53.42 |
| us | 4.33 | 5.00 | 16.00 | 66.33 | 8.67 | 66.67 | 13.00 | 6.00 | 3.67 | 5.33 | 9.67 | 21.00 | 77.33 | 43.33 |
| vif1 | 20.00 | 19.00 | 51.33 | 95.33 | 40.33 | 95.33 | 47.83 | 21.67 | 15.67 | 16.00 | 28.67 | 44.00 | 88.67 | 71.33 |
| vif2 | 8.33 | 8.00 | 27.33 | 80.67 | 17.00 | 81.67 | 25.33 | 10.33 | 7.67 | 9.67 | 19.00 | 31.00 | 88.00 | 53.67 |

| | | | | | | | | | | | | | | |
|---|---|---|---|---|---|---|---|---|---|---|---|---|---|---|
| vpr1 | 20.33 | 17.67 | 47.00 | 92.33 | 34.00 | 91.67 | 31.09 | 21.33 | 14.33 | 13.00 | 27.00 | 43.33 | 93.67 | 68.33 |
| vpr2 | 24.48 | 17.03 | 47.23 | 93.24 | 34.02 | 92.19 | 47.05 | 19.74 | 18.19 | 21.23 | 36.13 | 49.17 | 91.18 | 67.63 |
| vpr3 | 6.67 | 6.67 | 24.00 | 77.33 | 14.33 | 77.33 | 23.00 | 9.00 | 6.33 | 8.33 | 14.33 | 30.00 | 85.67 | 52.33 |
| vpr4 | 10.45 | 10.04 | 33.81 | 90.38 | 22.91 | 85.46 | 29.29 | 12.18 | 9.84 | 15.12 | 22.33 | 35.52 | 88.84 | 61.83 |

**C - SupT1_20h_pi rep2**

| | **8436** | **8442** | **8487** | **8532** | **8555** | **8563** | **8572** | **8632** | **9024** | **9074** | **9163** | **9400** | **9428** | **9442** |
|---|---|---|---|---|---|---|---|---|---|---|---|---|---|---|
| envvpu1 | 7.00 | 9.33 | 27.67 | 83.33 | 15.67 | 82.33 | 32.67 | 7.33 | 7.00 | 10.00 | 16.33 | 33.67 | 89.67 | 56.67 |
| envvpu13 | 5.51 | 11.81 | 40.84 | 83.11 | 23.31 | 93.33 | 28.77 | 12.52 | 11.52 | 12.71 | 23.43 | 40.38 | 90.21 | 67.64 |
| envvpu17 | 8.33 | 8.00 | 30.33 | 82.33 | 15.33 | 84.00 | 34.27 | 5.33 | 8.67 | 11.33 | 17.33 | 35.00 | 90.67 | 54.00 |
| envvpu19 | 7.36 | 15.50 | 19.51 | 85.73 | 17.85 | 83.70 | 21.41 | 3.94 | 5.55 | 9.42 | 11.26 | 34.93 | 91.77 | 56.19 |
| envvpu2 | 7.67 | 9.00 | 26.67 | 81.33 | 16.00 | 81.00 | 27.90 | 8.00 | 6.67 | 7.67 | 16.00 | 33.67 | 87.33 | 59.33 |
| envvpu3 | 4.67 | 8.33 | 26.67 | 81.67 | 14.33 | 80.33 | 32.57 | 6.33 | 6.67 | 10.67 | 18.00 | 31.33 | 88.67 | 56.33 |
| envvpu5 | 10.00 | 13.67 | 33.67 | 92.00 | 21.33 | 89.67 | 45.98 | 9.33 | 9.33 | 15.33 | 24.67 | 43.33 | 92.33 | 65.33 |
| envvpu9 | 8.00 | 9.67 | 29.33 | 87.33 | 18.00 | 86.00 | 35.67 | 9.00 | 8.00 | 11.67 | 18.67 | 36.00 | 90.67 | 61.00 |
| nef1 | 17.33 | 20.67 | 50.67 | 93.00 | 24.33 | 90.33 | 40.24 | 11.67 | 12.00 | 17.33 | 25.33 | 42.00 | 93.33 | 64.33 |
| nef12 | 11.57 | 15.67 | 39.67 | 90.33 | 26.67 | 90.33 | 49.08 | 6.67 | 8.33 | 8.67 | 16.73 | 44.47 | 88.07 | 56.91 |
| nef2 | 11.00 | 13.67 | 38.33 | 92.00 | 21.00 | 90.67 | 42.67 | 11.00 | 7.67 | 11.00 | 18.33 | 39.67 | 91.67 | 61.67 |
| nef3 | 9.67 | 12.33 | 39.00 | 92.00 | 23.00 | 91.33 | 43.67 | 11.33 | 8.67 | 11.67 | 19.00 | 42.00 | 90.67 | 62.67 |
| nef4 | 10.00 | 11.33 | 37.33 | 91.33 | 22.67 | 91.00 | 42.67 | 10.67 | 8.00 | 10.67 | 17.67 | 40.67 | 90.67 | 62.00 |

| | | | | | | | | | | | | | | |
|---|---|---|---|---|---|---|---|---|---|---|---|---|---|---|
| nef5 | 10.67 | 12.00 | 41.33 | 92.33 | 24.33 | 91.67 | 46.33 | 12.33 | 8.67 | 11.00 | 20.67 | 43.67 | 90.33 | 64.33 |
| nef9 | 12.00 | 16.00 | 39.00 | 92.00 | 20.67 | 90.33 | 43.33 | 10.33 | 6.67 | 11.33 | 18.67 | 40.00 | 91.67 | 63.33 |
| rev1 | 9.67 | 11.33 | 34.67 | 91.00 | 20.67 | 89.33 | 45.67 | 10.33 | 7.67 | 11.00 | 19.67 | 39.67 | 92.33 | 61.67 |
| rev10 | 7.36 | 14.90 | 36.50 | 92.38 | 16.01 | 87.69 | 54.58 | 15.88 | 11.23 | 10.62 | 26.01 | 46.63 | 90.64 | 65.18 |
| rev11 | 8.94 | 12.32 | 34.53 | 90.17 | 20.57 | 91.44 | 40.44 | 12.65 | 8.76 | 16.33 | 16.61 | 37.49 | 87.40 | 66.10 |
| rev2 | 9.00 | 11.67 | 33.33 | 91.00 | 20.33 | 90.00 | 46.00 | 11.00 | 8.00 | 11.00 | 18.67 | 38.00 | 91.67 | 61.67 |
| rev3 | 8.67 | 11.33 | 29.33 | 88.67 | 20.00 | 85.67 | 42.78 | 8.67 | 9.67 | 11.33 | 16.67 | 37.33 | 90.33 | 66.33 |
| rev4 | 9.67 | 11.33 | 35.33 | 91.67 | 23.00 | 90.67 | 33.33 | 12.67 | 9.00 | 14.00 | 18.33 | 43.33 | 89.33 | 64.33 |
| rev5 | 12.33 | 11.67 | 41.33 | 90.33 | 19.33 | 91.67 | 49.08 | 11.33 | 10.00 | 14.00 | 20.99 | 41.33 | 92.00 | 67.01 |
| rev7 | 7.33 | 10.33 | 31.00 | 90.33 | 20.00 | 93.00 | 41.47 | 10.67 | 9.33 | 13.00 | 19.67 | 45.67 | 92.00 | 65.67 |
| rev8 | 8.00 | 10.00 | 33.67 | 90.67 | 22.33 | 88.67 | 47.60 | 11.00 | 6.33 | 9.33 | 18.00 | 37.33 | 90.00 | 59.67 |
| SORF 1 | 12.90 | 15.75 | 39.56 | 89.18 | 19.86 | 89.88 | 55.00 | 10.23 | 10.16 | 16.21 | 18.23 | 39.04 | 92.70 | 70.88 |
| tat1 | 14.33 | 17.00 | 40.67 | 93.00 | 21.33 | 91.33 | 42.33 | 11.67 | 9.00 | 12.67 | 22.00 | 40.67 | 92.67 | 65.00 |
| tat2 | 13.00 | 16.67 | 43.67 | 93.67 | 22.67 | 91.33 | 45.67 | 11.33 | 9.67 | 14.33 | 23.67 | 42.33 | 92.00 | 66.33 |
| tat3 | 17.22 | 17.31 | 38.92 | 92.27 | 24.60 | 92.52 | 40.71 | 9.86 | 8.90 | 12.84 | 24.96 | 45.84 | 92.98 | 68.09 |
| tat4 | 10.88 | 21.89 | 40.90 | 92.19 | 28.41 | 92.94 | 45.39 | 15.05 | 13.82 | 11.10 | 24.26 | 45.05 | 90.82 | 62.67 |
| tat5 | 9.10 | 9.26 | 33.33 | 76.50 | 18.85 | 85.86 | 36.79 | 8.72 | 6.38 | 10.99 | 20.31 | 37.02 | 90.14 | 61.92 |
| us | 4.00 | 5.33 | 18.33 | 66.33 | 10.33 | 68.33 | 20.33 | 5.33 | 4.33 | 6.00 | 10.00 | 23.00 | 79.33 | 43.00 |
| vif1 | 14.69 | 12.97 | 43.05 | 88.56 | 24.58 | 89.12 | 65.00 | 12.22 | 12.64 | 7.87 | 23.27 | 39.49 | 89.69 | 64.71 |
| vif2 | 7.00 | 8.67 | 26.67 | 79.67 | 15.00 | 79.00 | 29.33 | 7.00 | 6.33 | 9.33 | 15.67 | 31.67 | 87.67 | 56.67 |
| vpr1 | 13.90 | 14.56 | 44.85 | 91.54 | 20.35 | 91.78 | 52.28 | 13.95 | 12.57 | 14.77 | 22.17 | 34.28 | 89.81 | 62.40 |
| vpr3 | 6.00 | 7.67 | 25.00 | 78.67 | 14.67 | 78.67 | 30.67 | 8.67 | 6.00 | 7.67 | 16.67 | 31.33 | 86.00 | 53.67 |
| vpr4 | 9.92 | 10.10 | 32.13 | 84.33 | 20.74 | 87.87 | 44.67 | 10.92 | 7.21 | 13.30 | 27.57 | 34.37 | 93.30 | 70.14 |

**D - SupT1_30h_pi rep2**

| | **8436** | **8442** | **8487** | **8532** | **8555** | **8563** | **8572** | **8632** | **9024** | **9074** | **9163** | **9400** | **9428** | **9442** |
|---|---|---|---|---|---|---|---|---|---|---|---|---|---|---|
| envvpu1 | 8.00 | 8.67 | 29.00 | 83.33 | 16.33 | 82.67 | 31.00 | 7.33 | 6.67 | 7.33 | 16.00 | 31.67 | 87.33 | 54.67 |

| | | | | | | | | | | | | | | |
|---|---|---|---|---|---|---|---|---|---|---|---|---|---|---|
| envvpu10 | 10.04 | 13.43 | 42.38 | 84.45 | 20.74 | 84.34 | 28.64 | 10.04 | 9.16 | 9.37 | 15.80 | 44.52 | 90.33 | 52.34 |
| envvpu13 | 11.25 | 10.50 | 37.82 | 88.60 | 23.48 | 86.85 | 34.33 | 9.75 | 11.90 | 10.92 | 27.77 | 38.29 | 91.43 | 67.07 |
| envvpu17 | 7.00 | 9.00 | 25.67 | 82.00 | 16.67 | 83.00 | 25.00 | 8.00 | 6.00 | 7.33 | 14.00 | 34.67 | 90.67 | 53.67 |
| envvpu19 | 11.76 | 12.37 | 21.95 | 80.58 | 18.20 | 74.26 | 12.62 | 5.77 | 10.69 | 6.07 | 14.18 | 31.64 | 81.57 | 55.50 |
| envvpu2 | 6.33 | 11.00 | 33.33 | 82.00 | 17.33 | 82.67 | 31.83 | 7.00 | 6.33 | 8.00 | 16.33 | 33.33 | 86.33 | 55.33 |
| envvpu3 | 10.00 | 9.67 | 31.00 | 82.00 | 12.67 | 82.67 | 40.27 | 5.33 | 6.67 | 6.67 | 13.67 | 31.00 | 85.33 | 54.67 |
| envvpu5 | 11.00 | 13.00 | 34.33 | 87.67 | 22.33 | 89.67 | 38.43 | 10.33 | 8.67 | 11.33 | 19.67 | 38.33 | 87.67 | 64.00 |
| envvpu9 | 9.67 | 11.33 | 32.67 | 86.33 | 20.00 | 87.00 | 37.67 | 8.33 | 7.33 | 7.67 | 18.00 | 33.33 | 88.00 | 58.67 |
| nef1 | 13.33 | 16.33 | 39.67 | 93.33 | 25.67 | 90.33 | 42.67 | 9.00 | 10.67 | 10.33 | 21.67 | 44.00 | 90.67 | 60.67 |
| nef12 | 12.86 | 16.06 | 40.30 | 89.27 | 26.76 | 86.66 | 30.48 | 11.27 | 8.87 | 9.13 | 14.84 | 35.55 | 89.22 | 53.35 |
| nef2 | 11.67 | 14.00 | 37.67 | 90.67 | 23.00 | 90.33 | 44.00 | 10.33 | 8.00 | 9.67 | 18.00 | 37.33 | 88.67 | 60.00 |
| nef3 | 11.00 | 13.00 | 36.67 | 90.33 | 25.00 | 90.67 | 46.33 | 11.33 | 8.33 | 9.67 | 20.00 | 38.00 | 86.33 | 62.00 |
| nef4 | 10.67 | 12.33 | 37.00 | 90.67 | 26.33 | 90.67 | 47.33 | 11.33 | 9.33 | 9.33 | 20.33 | 37.67 | 89.00 | 60.33 |
| nef5 | 10.67 | 14.00 | 38.67 | 90.67 | 25.67 | 91.67 | 44.00 | 11.00 | 10.00 | 12.67 | 21.33 | 40.67 | 86.67 | 62.00 |
| nef9 | 9.33 | 12.33 | 38.67 | 90.33 | 22.00 | 90.67 | 44.72 | 10.67 | 7.33 | 10.67 | 17.00 | 37.00 | 88.00 | 60.67 |
| rev1 | 9.67 | 11.67 | 33.67 | 90.33 | 22.00 | 89.67 | 48.00 | 10.67 | 7.67 | 9.00 | 18.00 | 36.67 | 89.00 | 58.33 |
| rev10 | 13.59 | 13.33 | 45.58 | 85.73 | 28.45 | 92.12 | 51.27 | 11.54 | 9.16 | 15.19 | 31.03 | 42.81 | 89.40 | 64.77 |
| rev11 | 10.55 | 11.91 | 37.78 | 87.59 | 30.72 | 92.47 | 31.67 | 6.56 | 12.77 | 13.86 | 17.80 | 45.54 | 87.64 | 66.33 |
| rev2 | 10.00 | 11.33 | 32.33 | 90.00 | 22.67 | 89.33 | 44.33 | 9.67 | 8.00 | 10.67 | 18.67 | 35.67 | 88.00 | 58.67 |
| rev3 | 8.03 | 12.48 | 32.47 | 88.74 | 21.95 | 88.51 | 29.44 | 7.93 | 6.87 | 6.84 | 15.89 | 40.58 | 87.56 | 58.70 |
| rev4 | 9.33 | 12.33 | 33.67 | 89.67 | 25.00 | 89.00 | 45.07 | 9.33 | 10.33 | 11.00 | 18.67 | 37.67 | 86.33 | 61.67 |
| rev5 | 11.11 | 13.88 | 39.50 | 88.82 | 20.74 | 87.76 | 46.06 | 12.71 | 3.55 | 7.06 | 21.04 | 34.63 | 85.67 | 57.48 |
| rev7 | 11.00 | 12.00 | 35.33 | 88.67 | 25.67 | 90.33 | 41.65 | 11.67 | 10.00 | 10.00 | 19.00 | 38.67 | 89.00 | 62.33 |
| rev8 | 9.67 | 12.00 | 35.67 | 90.33 | 25.00 | 92.67 | 50.03 | 12.33 | 12.67 | 11.33 | 20.67 | 37.00 | 88.67 | 59.33 |
| SORF1 | 12.42 | 16.89 | 32.11 | 94.40 | 24.17 | 88.26 | 40.44 | 7.05 | 8.57 | 10.78 | 23.63 | 39.87 | 90.45 | 68.20 |
| tat1 | 16.00 | 17.33 | 41.67 | 90.33 | 23.67 | 90.00 | 47.00 | 12.00 | 10.67 | 11.00 | 22.67 | 36.67 | 89.00 | 62.67 |
| tat2 | 16.33 | 18.00 | 42.00 | 90.67 | 27.00 | 91.33 | 50.33 | 10.67 | 10.00 | 11.67 | 24.00 | 36.67 | 88.67 | 64.33 |
| tat3 | 16.33 | 20.67 | 45.33 | 91.67 | 33.33 | 91.00 | 42.08 | 13.00 | 12.00 | 13.33 | 24.00 | 39.67 | 88.39 | 61.75 |
| tat4 | 19.67 | 15.32 | 48.11 | 91.97 | 31.46 | 86.39 | 57.16 | 12.60 | 12.49 | 12.06 | 26.10 | 38.97 | 87.02 | 71.48 |

| | | | | | | | | | | | | | | |
|---|---|---|---|---|---|---|---|---|---|---|---|---|---|---|
| tat5 | 9.51 | 7.87 | 32.99 | 83.32 | 17.43 | 78.74 | 30.08 | 8.93 | 9.43 | 11.52 | 16.67 | 29.45 | 89.08 | 61.24 |
| us | 4.00 | 4.67 | 18.33 | 67.67 | 11.67 | 68.00 | 21.33 | 5.00 | 4.33 | 5.33 | 9.67 | 22.00 | 79.33 | 43.00 |
| vif1 | 13.63 | 10.94 | 39.15 | 88.35 | 25.39 | 88.06 | 49.40 | 7.70 | 7.30 | 9.21 | 19.00 | 36.98 | 89.76 | 61.41 |
| vif2 | 7.67 | 8.00 | 27.33 | 81.67 | 17.33 | 81.00 | 32.67 | 7.67 | 7.33 | 7.00 | 15.67 | 29.67 | 85.33 | 55.00 |
| vpr1 | 14.67 | 16.00 | 45.67 | 91.00 | 26.67 | 93.33 | 45.95 | 15.33 | 10.00 | 9.33 | 21.67 | 35.67 | 92.00 | 65.67 |
| vpr3 | 7.00 | 7.33 | 26.67 | 78.00 | 15.67 | 79.33 | 32.00 | 8.33 | 6.33 | 7.33 | 14.33 | 28.33 | 85.33 | 52.00 |
| vpr4 | 10.00 | 8.67 | 36.33 | 85.00 | 18.67 | 88.00 | 44.12 | 14.67 | 9.67 | 7.67 | 25.46 | 39.67 | 91.33 | 58.33 |

## E - CD4_48h_pi

| | 8436 | 8442 | 8487 | 8532 | 8555 | 8563 | 8572 | 8632 | 9024 | 9074 | 9163 | 9400 | 9428 | 9442 |
|---|---|---|---|---|---|---|---|---|---|---|---|---|---|---|
| envvpu1 | 5.00 | 8.00 | 25.67 | 79.67 | 14.67 | 79.67 | 30.67 | 5.33 | 5.00 | 7.33 | 13.33 | 26.00 | 86.33 | 48.67 |
| envvpu2 | 6.88 | 9.81 | 19.92 | 74.43 | 11.32 | 77.99 | 23.33 | 5.86 | 3.03 | 5.14 | 16.86 | 27.63 | 85.65 | 53.03 |
| envvpu3 | 4.56 | 9.51 | 22.62 | 75.73 | 12.23 | 74.77 | 36.21 | 7.90 | 5.92 | 11.27 | 11.24 | 26.61 | 85.33 | 46.48 |
| envvpu5 | 7.14 | 11.93 | 45.08 | 85.65 | 20.36 | 83.00 | 31.67 | 8.24 | 6.46 | 10.88 | 23.21 | 29.81 | 86.67 | 55.11 |
| envvpu9 | 5.67 | 6.67 | 29.67 | 83.33 | 15.00 | 81.67 | 31.26 | 6.67 | 7.33 | 11.00 | 14.00 | 26.67 | 86.67 | 50.33 |
| nef1 | 10.71 | 14.12 | 38.38 | 87.90 | 22.53 | 85.38 | 52.33 | 10.98 | 6.18 | 6.73 | 17.40 | 32.87 | 90.02 | 61.40 |
| nef2 | 9.00 | 12.00 | 36.00 | 89.33 | 20.33 | 88.67 | 41.67 | 8.33 | 6.67 | 10.00 | 15.33 | 32.33 | 87.67 | 54.67 |
| nef3 | 9.00 | 12.67 | 36.00 | 89.33 | 22.33 | 88.33 | 39.82 | 7.33 | 6.33 | 9.33 | 19.67 | 35.33 | 86.67 | 57.00 |
| nef4 | 8.33 | 13.67 | 37.67 | 88.33 | 21.33 | 87.67 | 39.67 | 8.67 | 6.67 | 8.67 | 16.00 | 31.67 | 88.33 | 55.67 |
| nef5 | 15.67 | 10.33 | 38.67 | 90.33 | 22.33 | 88.33 | 50.71 | 9.33 | 8.67 | 13.67 | 19.67 | 37.67 | 88.33 | 61.00 |
| nef9 | 7.44 | 11.73 | 39.31 | 85.17 | 27.79 | 88.61 | 48.33 | 6.30 | 9.77 | 9.36 | 13.97 | 31.99 | 85.01 | 52.10 |
| rev1 | 8.00 | 12.00 | 30.33 | 87.33 | 18.00 | 85.67 | 42.33 | 7.33 | 6.00 | 9.33 | 16.00 | 30.33 | 86.00 | 53.33 |
| rev11 | 10.41 | 10.87 | 40.27 | 89.91 | 31.81 | 91.85 | 52.50 | 14.30 | 12.47 | 13.49 | 20.85 | 44.22 | 89.89 | 58.00 |
| rev2 | 8.00 | 10.33 | 33.00 | 89.33 | 19.00 | 87.33 | 38.33 | 8.33 | 5.33 | 9.00 | 15.33 | 32.67 | 88.33 | 54.00 |
| rev3 | 8.67 | 10.00 | 29.67 | 87.00 | 14.67 | 85.33 | 46.68 | 6.33 | 5.67 | 10.00 | 17.67 | 29.67 | 85.67 | 51.67 |
| rev4 | 6.54 | 9.91 | 30.21 | 82.20 | 20.12 | 82.49 | 40.27 | 5.80 | 5.09 | 9.10 | 16.12 | 30.78 | 86.17 | 53.41 |
| rev5 | 11.88 | 10.11 | 28.55 | 82.50 | 24.56 | 84.00 | 48.33 | 10.19 | 8.26 | 12.17 | 20.49 | 31.64 | 86.95 | 56.69 |
| rev7 | 11.00 | 9.67 | 35.00 | 88.00 | 21.33 | 88.00 | 51.56 | 11.00 | 7.00 | 11.33 | 17.67 | 37.33 | 90.00 | 53.67 |
| rev8 | 8.00 | 10.33 | 33.33 | 90.00 | 22.00 | 88.00 | 41.19 | 9.33 | 6.67 | 9.00 | 19.67 | 33.67 | 88.00 | 55.33 |

| | | | | | | | | | | | | | | |
|---|---|---|---|---|---|---|---|---|---|---|---|---|---|---|
| tat1 | 11.67 | 16.67 | 38.67 | 88.67 | 19.67 | 88.00 | 38.67 | 10.00 | 7.33 | 11.33 | 17.00 | 31.67 | 89.67 | 54.00 |
| tat2 | 12.00 | 15.67 | 41.67 | 90.67 | 23.67 | 89.33 | 40.12 | 9.00 | 8.67 | 15.00 | 19.00 | 32.33 | 89.00 | 61.67 |
| tat3 | 12.41 | 15.42 | 39.63 | 90.03 | 21.29 | 91.69 | 42.78 | 9.31 | 10.10 | 14.32 | 14.13 | 29.71 | 89.46 | 52.37 |
| tat5 | 6.56 | 9.33 | 29.17 | 80.74 | 14.61 | 83.35 | 36.52 | 7.36 | 5.60 | 8.60 | 17.60 | 22.48 | 85.54 | 54.73 |
| us | 3.67 | 4.67 | 17.67 | 64.33 | 10.33 | 69.33 | 23.67 | 5.00 | 4.00 | 4.00 | 7.33 | 19.00 | 78.33 | 39.00 |
| vif2 | 6.67 | 5.67 | 24.00 | 76.67 | 16.67 | 77.33 | 36.54 | 9.67 | 4.67 | 7.67 | 13.00 | 24.33 | 82.33 | 51.00 |
| vpr1 | 7.48 | 11.70 | 36.88 | 92.20 | 20.21 | 87.73 | 40.27 | 10.35 | 9.30 | 14.51 | 22.44 | 35.42 | 84.85 | 56.12 |
| vpr3 | 7.33 | 9.33 | 28.67 | 76.33 | 15.67 | 79.00 | 33.33 | 7.00 | 6.33 | 7.00 | 15.00 | 22.00 | 83.00 | 46.33 |
| vpr4 | 9.58 | 16.33 | 37.28 | 88.04 | 19.93 | 91.14 | 58.94 | 9.19 | 11.48 | 9.34 | 16.39 | 33.22 | 85.62 | 53.77 |

**F - CD4_72h_pi**

| | **8436** | **8442** | **8487** | **8532** | **8555** | **8563** | **8572** | **8632** | **9024** | **9074** | **9163** | **9400** | **9428** | **9442** |
|---|---|---|---|---|---|---|---|---|---|---|---|---|---|---|
| envvpu1 | 4.67 | 8.00 | 26.67 | 78.00 | 11.33 | 77.00 | 21.33 | 4.67 | 4.33 | 8.67 | 12.33 | 25.67 | 88.00 | 52.33 |
| envvpu17 | 3.04 | 5.98 | 23.47 | 79.26 | 10.34 | 77.79 | 10.83 | 2.76 | 4.93 | 9.04 | 16.20 | 30.97 | 86.44 | 49.67 |
| envvpu2 | 9.81 | 5.26 | 33.25 | 79.40 | 8.52 | 79.24 | 31.67 | 6.47 | 3.75 | 9.94 | 15.54 | 26.23 | 86.61 | 56.40 |
| envvpu3 | 7.00 | 7.33 | 25.67 | 78.33 | 12.67 | 76.67 | 21.22 | 4.33 | 5.00 | 8.00 | 10.33 | 23.33 | 85.67 | 47.67 |
| envvpu9 | 6.45 | 6.54 | 22.60 | 83.44 | 14.22 | 82.11 | 28.89 | 3.37 | 5.13 | 9.50 | 13.86 | 26.36 | 83.67 | 49.67 |
| nef1 | 10.19 | 11.43 | 33.13 | 86.54 | 11.09 | 77.04 | 23.33 | 6.09 | 3.35 | 8.38 | 7.84 | 34.98 | 89.44 | 54.64 |
| nef12 | 5.22 | 14.82 | 35.96 | 92.74 | 7.08 | 88.78 | 31.67 | 0.71 | 1.73 | 10.30 | 15.33 | 28.93 | 92.42 | 67.41 |
| nef2 | 7.33 | 11.00 | 33.33 | 89.00 | 13.67 | 86.00 | 32.33 | 6.33 | 4.67 | 9.00 | 13.00 | 29.67 | 92.00 | 58.00 |
| nef3 | 7.67 | 10.33 | 33.00 | 89.33 | 14.33 | 86.67 | 34.60 | 8.67 | 4.00 | 8.67 | 15.33 | 30.33 | 92.33 | 60.00 |
| nef4 | 6.33 | 11.00 | 33.33 | 87.33 | 16.67 | 86.00 | 31.96 | 7.67 | 6.67 | 11.00 | 16.67 | 31.00 | 90.33 | 58.00 |
| nef5 | 14.43 | 14.90 | 29.77 | 90.81 | 17.71 | 88.27 | 37.22 | 14.41 | 8.11 | 17.00 | 11.50 | 38.55 | 92.33 | 56.31 |
| nef9 | 4.33 | 10.00 | 29.67 | 88.67 | 15.67 | 86.00 | 27.37 | 7.67 | 3.00 | 9.00 | 15.67 | 24.33 | 94.67 | 59.67 |
| rev1 | 6.33 | 9.33 | 29.00 | 86.00 | 14.33 | 84.67 | 32.95 | 6.33 | 4.33 | 7.67 | 12.67 | 26.00 | 91.00 | 56.67 |
| rev2 | 7.33 | 10.00 | 29.00 | 87.00 | 13.67 | 83.67 | 31.33 | 6.00 | 5.00 | 9.67 | 13.00 | 28.33 | 91.67 | 57.00 |
| rev3 | 6.00 | 10.33 | 33.00 | 88.00 | 13.33 | 83.00 | 33.01 | 7.67 | 4.33 | 9.67 | 12.67 | 26.00 | 91.00 | 56.33 |
| rev4 | 5.27 | 9.10 | 36.86 | 88.56 | 8.62 | 82.52 | 15.72 | 5.38 | -0.33 | 6.33 | 11.00 | 26.69 | 92.42 | 59.45 |
| rev5 | 4.30 | 15.41 | 36.27 | 89.67 | 13.95 | 87.63 | 45.00 | 3.37 | 6.21 | 10.24 | 17.81 | 34.24 | 90.67 | 65.05 |

| | | | | | | | | | | | | | | |
|---|---|---|---|---|---|---|---|---|---|---|---|---|---|---|
| rev8 | 7.62 | 9.22 | 26.06 | 87.77 | 13.13 | 81.19 | 28.64 | 5.19 | 4.04 | 8.19 | 15.77 | 24.06 | 91.15 | 57.47 |
| tat1 | 9.00 | 14.67 | 35.33 | 88.00 | 14.67 | 86.33 | 32.67 | 7.00 | 6.00 | 12.67 | 15.67 | 27.00 | 92.67 | 59.67 |
| tat2 | 9.33 | 14.67 | 34.67 | 90.00 | 16.67 | 88.00 | 33.57 | 7.33 | 7.33 | 12.00 | 18.00 | 32.67 | 91.67 | 63.00 |
| tat5 | 4.34 | 7.47 | 29.48 | 73.75 | 14.33 | 79.96 | 22.06 | 7.59 | 7.59 | 9.82 | 16.89 | 22.01 | 88.27 | 57.73 |
| us | 2.67 | 4.67 | 16.33 | 60.00 | 7.33 | 62.33 | 16.33 | 4.00 | 3.00 | 5.33 | 8.67 | 17.33 | 77.67 | 38.67 |
| vif1 | 9.57 | 14.21 | 32.94 | 75.81 | 21.43 | 81.91 | 28.33 | 11.70 | 14.35 | 8.08 | 22.00 | 29.61 | 81.69 | 48.69 |
| vif2 | 7.67 | 6.67 | 23.33 | 79.00 | 15.33 | 77.33 | 23.50 | 5.33 | 10.00 | 9.00 | 16.67 | 21.00 | 86.33 | 50.67 |
| vpr1 | 12.31 | 8.51 | 38.77 | 80.33 | 16.22 | 83.73 | 36.11 | 7.49 | 8.46 | 8.05 | 20.18 | 26.76 | 84.21 | 58.97 |
| vpr3 | 5.33 | 8.67 | 23.00 | 73.67 | 13.00 | 74.33 | 24.84 | 5.00 | 5.00 | 7.00 | 16.67 | 22.00 | 84.00 | 48.33 |

**Table S8.** Rates of methylation of the 14 $m^6A$ sites detected in all transcripts of the splicing isoforms found in the samples indicated. **(A-F)** Rates for the isoforms represented by ≥ 100 full-length reads (Materials and Methods).

| Isoform | | SupT1_20h_pi rep1 | | | |
|---|---|---|---|---|---|
| CS - i4 | PS + i4 | %methyl (5613) | %methy l (5654) | %methyl (5703) | %methyl (5887) |
| Vif1 | | 35.2 | 2.4 | 11.4 | 18.6 |
| | Vif2 | 9.3 | 0.7 | 3.3 | 2 |
| Vpr1 | | 58 | 9.3 | 24 | 25.3 |
| | Vpr3 | 12.3 | 1 | 4 | 0.3 |
| Vpr2 | | 37.3 | 4.6 | 10 | 15.9 |
| | Vpr4 | 5.7 | 2.3 | 3.7 | 0 |
| Tat1 | | | | | 18.3 |
| | Tat5 | | | | 28.7 |
| Tat2 | | | | | 12.3 |
| Tat3 | | | | | 12.3 |
| Tat4 | | | | | 9.5 |
| SORF1 | | | | | 16.7 |
| US | | 1.3 | 0 | 0 | 0 |

| Isoform | | SupT1_20h_pi rep2 | | | |
|---|---|---|---|---|---|
| CS - i4 | PS + i4 | %methyl (5613) | %methy l (5654) | %methyl (5703) | %methyl (5887) |
| Vif1 | | 15.7 | 1.2 | 5.9 | 5.5 |
| | Vif2 | 7.3 | 0.3 | 1.7 | 1.3 |
| Vpr1 | | 40.1 | 4.7 | 15.8 | 14.8 |
| | Vpr3 | 8.7 | 0.3 | 2 | 1 |
| Vpr2 | | 32 | 6.7 | 19.2 | 16.3 |
| | Vpr4 | 13.3 | 0.3 | 0.9 | 2.1 |
| Tat1 | | | | | 17 |
| | Tat5 | | | | 24.3 |
| Tat2 | | | | | 11.7 |
| Tat3 | | | | | 16.6 |
| Tat4 | | | | | 16.6 |
| SORF1 | | | | | 16.7 |
| US | | 1.0 | 0 | 0 | 0.3 |

| Isoform | | SupT1_30h_pi rep1 | | | |
|---|---|---|---|---|---|
| CS<br>- i4 | PS<br>+ i4 | %methyl (5613) | %methyl (5654) | %methyl (5703) | %methyl (5887) |
| Vif1 | | 21 | 4 | 5.7 | 10.7 |
| | Vif2 | 8.7 | 0 | 2.0 | 1 |
| Vpr1 | | 40.3 | 5 | 12 | 13.7 |
| | Vpr3 | 10.3 | 0.3 | 1.3 | 0.3 |
| Vpr2 | | 39.0 | 8.0 | 18.5 | 15 |
| | Vpr4 | 12.6 | 0.5 | 6.1 | 2.0 |
| Tat1 | | | | | 16.7 |
| | Tat5 | | | | 18.8 |
| Tat2 | | | | | 13 |
| Tat3 | | | | | 13.3 |
| Tat4 | | | | | 18.3 |
| SORF1 | | | | | 16 |
| US | | 1.3 | 0 | 0 | 0 |

| Isoform | | SupT1_30h_pi rep2 | | | |
|---|---|---|---|---|---|
| CS<br>- i4 | PS<br>+ i4 | %methyl (5613) | %methyl (5654) | %methyl (5703) | %methyl (5887) |
| Vif1 | | 18.7 | 1.4 | 9.4 | 7.0 |
| | Vif2 | 7 | 0.7 | 1.7 | 1 |
| Vpr1 | | 42.3 | 7.7 | 17 | 13.3 |
| | Vpr3 | 9.3 | 1 | 2 | 0.3 |
| Vpr2 | | 41.0 | 5.5 | 13.8 | 6.4 |
| | Vpr4 | 10 | 2.1 | 3 | 1.3 |
| Tat1 | | | | | 16.7 |
| | Tat5 | | | | 19.7 |
| Tat2 | | | | | 12.0 |
| Tat3 | | | | | 14.3 |
| Tat4 | | | | | 14.3 |
| SORF1 | | | | | 16.1 |
| US | | 1.0 | 0.3 | 0.3 | 0 |

| Isoform | | CD4_48h_pi | | | |
|---|---|---|---|---|---|
| CS - i4 | PS + i4 | %methyl (5613) | %methyl (5654) | %methyl (5703) | %methyl (5887) |
| Vif1 | | 15.1 | 5.3 | 5.2 | 10.8 |
| | Vif2 | 6.7 | 0.3 | 1.3 | 2 |
| Vpr1 | | 43.5 | 5.1 | 8.8 | 14.0 |
| | Vpr3 | 8.7 | 0.7 | 1 | 0.7 |
| Vpr2 | | 42.2 | 12.3 | 18.7 | 33.0 |
| | Vpr4 | 8.3 | 0 | 1.2 | 2.8 |
| Tat1 | | | | | 19.3 |
| | Tat5 | | | | 23.5 |
| Tat2 | | | | | 13.7 |
| Tat3 | | | | | 13.5 |
| Tat4 | | | | | 16.6 |
| SORF1 | | | | | 15.3 |
| US | | 1.0 | 0 | 0 | 0.3 |

| Isoform | | CD4_72h_pi | | | |
|---|---|---|---|---|---|
| CS - i4 | PS + i4 | %methyl (5613) | %methyl (5654) | %methyl (5703) | %methyl (5887) |
| Vif1 | | 9.8 | 1.0 | 4.5 | 6.0 |
| | Vif2 | 8 | 1.0 | 2.3 | 1.3 |
| Vpr1 | | 27.1 | 4.4 | 13.9 | 9.1 |
| | Vpr3 | 8 | 0.7 | 3 | 1 |
| Vpr2 | | 32.7 | 0 | 0 | 0 |
| | Vpr4 | 13.3 | 0 | 3.4 | 3.6 |
| Tat1 | | | | | 13 |
| | Tat5 | | | | 21.1 |
| Tat2 | | | | | 8 |
| Tat3 | | | | | 9.1 |
| Tat4 | | | | | 20.4 |
| SORF1 | | | | | 16.7 |
| US | | 1.3 | 0.3 | 0 | 0.3 |

**Table S9.** Methylation rates for the A5613, A5654, A5703 and A5887 $m^6A$ sites in the Vif, Vpr, Tat and SORF1 isoforms, as measured in the samples indicated. The first two columns indicate the isoforms completely spliced (CS, lacking intron 4, - i4) or partially spliced (PS, containing intron 4, + i4); US, unspliced transcript.

**A**

**SupT1_20h_pi rep1**

| | US | | | | PS | | | | CS | | | |
|---|---|---|---|---|---|---|---|---|---|---|---|---|
| | 0 | 1 | 2 | 3 | 0 | 1 | 2 | 3 | 0 | 1 | 2 | 3 |
| 0 | 19 | 26 | 11 | 1 | 34 | 75 | 39 | 7 | 86 | 82 | 72 | 12 |
| 1 | 16 | 61 | 24 | 3 | 65 | 313 | 274 | 54 | 149 | 876 | 1066 | 290 |
| 2 | 9 | 80 | 75 | 22 | 109 | 679 | 864 | 261 | 561 | 4310 | 6522 | 2136 |
| 3 | 7 | 33 | 47 | 9 | 54 | 540 | 854 | 313 | 655 | 5241 | 9068 | 3503 |
| 4 | 1 | 8 | 16 | 1 | 26 | 208 | 433 | 202 | 284 | 2878 | 5581 | 2679 |
| 5 | 0 | 3 | 4 | 2 | 5 | 45 | 115 | 84 | 100 | 861 | 2005 | 1203 |
| 6 | 0 | 0 | 0 | 0 | 0 | 13 | 24 | 14 | 18 | 162 | 427 | 303 |
| 7 | 0 | 0 | 0 | 0 | 0 | 2 | 1 | 0 | 1 | 19 | 50 | 57 |
| 8 | 0 | 0 | 0 | 0 | 0 | 0 | 0 | 0 | 0 | 2 | 4 | 4 |

**B**

**SupT1_20h_pi rep2**

| | US | | | | PS | | | | CS | | | |
|---|---|---|---|---|---|---|---|---|---|---|---|---|
| | 0 | 1 | 2 | 3 | 0 | 1 | 2 | 3 | 0 | 1 | 2 | 3 |
| 0 | 341 | 503 | 167 | 18 | 194 | 396 | 220 | 34 | 147 | 143 | 124 | 29 |
| 1 | 366 | 1076 | 644 | 95 | 293 | 1373 | 1320 | 287 | 251 | 1508 | 1962 | 570 |
| 2 | 252 | 1280 | 1298 | 282 | 346 | 2877 | 3872 | 1212 | 722 | 6740 | 11069 | 4025 |
| 3 | 78 | 566 | 737 | 191 | 213 | 1826 | 3268 | 1234 | 739 | 6966 | 13411 | 5547 |
| 4 | 13 | 135 | 222 | 77 | 50 | 605 | 1268 | 615 | 332 | 3078 | 6830 | 3278 |
| 5 | 3 | 24 | 41 | 15 | 12 | 137 | 279 | 183 | 74 | 719 | 1696 | 982 |

| | | | | | | | | | | | | |
|---|---|---|---|---|---|---|---|---|---|---|---|---|
| 6 | 0 | 5 | 7 | 5 | 1 | 19 | 26 | 30 | 4 | 75 | 247 | 168 |
| 7 | 0 | 0 | 0 | 1 | 0 | 3 | 4 | 0 | 0 | 4 | 19 | 18 |
| 8 | 0 | 0 | 0 | 0 | 0 | 0 | 0 | 0 | 0 | 0 | 0 | 0 |

**C**
**SupT1_30h_pi rep1**

| | US | | | | PS | | | | CS | | | |
|---|---|---|---|---|---|---|---|---|---|---|---|---|
| | 0 | 1 | 2 | 3 | 0 | 1 | 2 | 3 | 0 | 1 | 2 | 3 |
| 0 | 71 | 110 | 50 | 4 | 134 | 259 | 165 | 21 | 138 | 170 | 194 | 53 |
| 1 | 91 | 238 | 156 | 24 | 251 | 1125 | 961 | 208 | 375 | 2112 | 2709 | 881 |
| 2 | 52 | 308 | 278 | 71 | 316 | 2232 | 2757 | 804 | 1318 | 9691 | 14648 | 5491 |
| 3 | 29 | 134 | 157 | 31 | 156 | 1460 | 2359 | 938 | 1304 | 10289 | 17984 | 7611 |
| 4 | 5 | 28 | 52 | 12 | 58 | 481 | 955 | 452 | 596 | 4745 | 9458 | 4762 |
| 5 | 2 | 3 | 7 | 2 | 13 | 128 | 240 | 142 | 135 | 1211 | 2667 | 1634 |
| 6 | 0 | 0 | 1 | 0 | 0 | 19 | 43 | 22 | 17 | 170 | 442 | 349 |
| 7 | 0 | 0 | 0 | 0 | 0 | 1 | 1 | 4 | 1 | 19 | 52 | 47 |
| 8 | 0 | 0 | 0 | 0 | 0 | 0 | 0 | 0 | 0 | 0 | 3 | 5 |

**D**
**SupT1_30h_pi rep2**

| | US | | | | PS | | | | CS | | | |
|---|---|---|---|---|---|---|---|---|---|---|---|---|
| | 0 | 1 | 2 | 3 | 0 | 1 | 2 | 3 | 0 | 1 | 2 | 3 |
| 0 | 496 | 740 | 257 | 25 | 265 | 432 | 264 | 42 | 122 | 139 | 143 | 31 |
| 1 | 609 | 1681 | 1048 | 158 | 440 | 1736 | 1520 | 300 | 237 | 1301 | 1541 | 423 |

| | | | | | | | | | | | | |
|---|---|---|---|---|---|---|---|---|---|---|---|---|
| 2 | 428 | 2110 | 1999 | 389 | 558 | 3630 | 4691 | 1300 | 854 | 5717 | 8325 | 2608 |
| 3 | 160 | 945 | 1174 | 284 | 319 | 2405 | 4023 | 1292 | 792 | 5991 | 9961 | 3600 |
| 4 | 22 | 215 | 344 | 94 | 105 | 847 | 1596 | 643 | 328 | 2685 | 5025 | 2192 |
| 5 | 6 | 32 | 53 | 27 | 17 | 152 | 371 | 160 | 28 | 584 | 1332 | 649 |
| 6 | 0 | 3 | 3 | 3 | 0 | 11 | 60 | 16 | 11 | 68 | 202 | 134 |
| 7 | 0 | 0 | 0 | 0 | 1 | 0 | 1 | 3 | 0 | 7 | 14 | 11 |
| 8 | 0 | 0 | 0 | 0 | 0 | 0 | 0 | 0 | 0 | 1 | 0 | 0 |

**E**
**CD4_48h_pi**

| | US | | | | PS | | | | CS | | | |
|---|---|---|---|---|---|---|---|---|---|---|---|---|
| | 0 | 1 | 2 | 3 | 0 | 1 | 2 | 3 | 0 | 1 | 2 | 3 |
| 0 | 81 | 95 | 35 | 7 | 94 | 151 | 84 | 16 | 51 | 78 | 52 | 23 |
| 1 | 90 | 253 | 134 | 12 | 123 | 562 | 370 | 59 | 120 | 623 | 613 | 125 |
| 2 | 64 | 293 | 277 | 45 | 172 | 1015 | 1097 | 211 | 320 | 2203 | 2606 | 736 |
| 3 | 14 | 156 | 155 | 26 | 103 | 688 | 746 | 215 | 275 | 2048 | 2930 | 901 |
| 4 | 2 | 30 | 34 | 7 | 24 | 208 | 306 | 106 | 104 | 790 | 1242 | 453 |
| 5 | 2 | 10 | 9 | 4 | 9 | 36 | 56 | 19 | 18 | 158 | 322 | 152 |
| 6 | 0 | 0 | 1 | 0 | 1 | 5 | 10 | 6 | 4 | 17 | 42 | 29 |
| 7 | 0 | 0 | 0 | 0 | 0 | 0 | 1 | 2 | 0 | 0 | 3 | 4 |
| 8 | 0 | 0 | 0 | 0 | 0 | 0 | 0 | 1 | 0 | 0 | 0 | 0 |

**F**
**CD4_72h_pi**

| | US | | | | PS | | | | CS | | | |
|---|---|---|---|---|---|---|---|---|---|---|---|---|
| | 0 | 1 | 2 | 3 | 0 | 1 | 2 | 3 | 0 | 1 | 2 | 3 |
| 0 | 148 | 250 | 73 | 4 | 78 | 165 | 77 | 10 | 55 | 111 | 91 | 16 |
| 1 | 129 | 418 | 214 | 32 | 108 | 513 | 425 | 68 | 168 | 1000 | 1021 | 200 |
| 2 | 82 | 419 | 366 | 68 | 116 | 902 | 943 | 256 | 402 | 3123 | 4221 | 1138 |
| 3 | 26 | 178 | 170 | 48 | 65 | 510 | 762 | 189 | 273 | 2264 | 3585 | 1129 |
| 4 | 7 | 31 | 43 | 12 | 9 | 126 | 211 | 78 | 67 | 700 | 1273 | 508 |
| 5 | 0 | 3 | 6 | 1 | 5 | 15 | 40 | 16 | 10 | 118 | 244 | 141 |
| 6 | 0 | 2 | 0 | 0 | 1 | 0 | 4 | 2 | 4 | 10 | 21 | 28 |
| 7 | 0 | 0 | 0 | 0 | 0 | 0 | 0 | 1 | 0 | 0 | 2 | 1 |
| 8 | 0 | 0 | 0 | 0 | 0 | 0 | 0 | 0 | 0 | 0 | 1 | 0 |

**Table S10.** Single-molecule analysis of methylated $m^6A$ sites illustrating the correlation between the methylation of clusters 1 and 2 of the same transcript, by sample. **(A-F)** Number of reads in the indicated samples with defined numbers of methylated As in cluster 1 (by line: 0 to 8 $m^6As$) and in cluster 2 (by column: 0 to 3 $m^6As$). US, unspliced; PS partially spliced; CS, completely spliced. One $m^6A$ of one read is considered methylated if the probability of base modification is ≥ 0.5 (value in modkit extract, Materials and Methods).

| Position | Sequence | SupT1_20h_pi rep1 | SupT1_20h_pi rep2 | SupT1_30h_pi rep1 | SupT1_30h_pi rep2 | CD4_48h_pi | CD4_72h_pi |
|---|---|---|---|---|---|---|---|
| | | % methyl. | % methyl. | % methyl. | % methyl. | % methyl. * | % methyl. * |
| 2L125 | UGACU | 35.3 | 30.7 | 33.3 | 35 | 38.4 | 24.4 |
| 2L183 | AGACU | 11.7 | 9 | 13.7 | 9 | 6.5 | NS |
| 2L272 | GAACU | 27.7 | 30.7 | 30.3 | 29 | 21.9 | 26 |
| 2L509 | GAACU | 32.7 | 37.3 | 35 | 32.7 | 26.8 | 19.7 |
| 2L537 | GGACU | 90.3 | 91.3 | 89.3 | 90 | 94.1 | 79.2 |
| 2L551 | GGACU | 66.3 | 66.7 | 68.7 | 70 | 69.2 | 64 |

**Table S11.** Rates of methylation of the six $m^6A$ sites detected in the 2-LTR transcripts. The names of the sites refer to the position of the corresponding A in the HIV-1 reference genome. * indicates that the sequence context of the A9074 in the 2-LTR transcript (AGACU, Figure 6B) differs from the context of the same position, 9074, in the reference genome (GGACU). NS, p-value > $10^{-5}$.
